# Chromatin-associated intronic RNAs from long genes form introsomes that shape nuclear architecture in neuronal cells

**DOI:** 10.1101/2025.03.13.641826

**Authors:** Wenjing Kang, Wing Hin Yip, Xiaoze Li-Wang, Quentin Verron, Britta A.M. Bouwman, Simona Pedrotti, Andrea Abou Yaghi, Mitsuyoshi Murata, Lorenzo Salviati, Erik Wernersson, Yali Zhang, Marika Oksanen, Francesca Mastropasqua, Xufeng Shu, Rodrigo Pracana, Jay W. Shin, Takeya Kasukawa, Chi Wai Yip, Masaki Kato, Hazuki Takahashi, Kristiina Tammimies, Nicola Crosetto, Piero Carninci, Magda Bienko

## Abstract

Cell differentiation towards neurons is accompanied by widespread changes in three-dimensional (3D) genome organization and gene expression. Chromatin-associated RNAs have been proposed to be important regulators of such changes; however, the type, abundance, and role of these RNAs during neuronal differentiation remain largely unexplored. Here, we integrate multi-omic data generated in the frame of the Functional ANnoTation Of the Mammalian genome (FANTOM6) to chart 3D genome, RNA-DNA contactome, and transcriptome changes occurring during in vitro differentiation of human induced pluripotent stem cells to neural stem cells and neurons. We reveal a previously unreported phenomenon, in which intronic RNAs engage in long-distance contacts with DNA loci distributed all over the genome. These trans-contacting intronic RNAs (TIRs) are produced from exceptionally long (mean length: 750 kilobases, kb) protein-coding genes that carry ultra-long introns and are selectively expressed in neurons. We show that TIRs do not undergo rapid co-transcriptional degradation but rather accumulate in the nucleus of neuronal cells, forming large ‘dot clouds’ around their source loci and spreading across the nucleus, as visualized by single-molecule RNA fluorescence in situ hybridization. TIRs engage in contacts with a set of genomic regions (TIR-contacted regions or TIRCs) that carry much shorter (mean length: ∼30 kb) neuronally expressed genes forming high-connectivity hubs. We also show that the expression of genes within TIRCs contacted by the same set of TIRs is highly co-varied, and that TIR source genes, especially their introns, are enriched in genetic risk loci for neurodevelopmental and neuropsychiatric disorders. Our findings point to a functional role of the persistence of long intronic RNAs in the nucleus of neuronal cells and might contribute to explain why neurons uniquely express many ultra-long genes. We propose a model in which TIRs form pan-nuclear scaffolds—which we propose to name introsomes—that constitute a dynamically self-renewing regulatory layer, where transcription itself continuously regenerates the very scaffold that organizes genome function in neurons, and that might be involved in the pathogenesis of neurodevelopmental and neuropsychiatric disorders.

## Introduction

The three-dimensional (3D) spatial organization of the genome in the cell nucleus provides a multi-scale framework for spatio-temporal control of key genomic functions such as DNA replication, transcription, and repair^1^. Substantial 3D genome reorganization occurs as cells differentiate during embryonic development and in postnatal tissues. While pluripotent stem cells harbor an open 3D genome configuration that supports low-intensity genome-wide transcription, as cells exit pluripotency and undergo lineage commitment, their 3D genome undergoes drastic reorganization accompanied by the activation of lineage-specific transcriptional programs^2,3^. Multi-scale 3D genome rewiring is particularly evident during brain development, with chromatin loops and topologically associating domain (TAD) boundaries undergoing cell type-specific remodeling during neuronal lineage specification, both in vitro and in vivo^4–7^.

A notable property of many of the genes that become specifically activated in neurons and encode for proteins essential for neuronal functions—including numerous transmembrane ion channels and synaptic proteins—is that these genes are substantially longer than the average gene in the same organism^8^. This exceptional length of genes specifically expressed in neuronal cells is conserved across the eukaryotic kingdom, reaching megabase range in humans^9–11^. It has been hypothesized previously that the length of neuron-specific genes might have evolved to enlarge the gene regulatory landscape of neurons, by providing an expanded repertoire of new regulatory elements, alternative isoforms, and exons^10,11^. However, this hypothesis does not account for the fact that the length of neuron-specific genes is, to a large extent, attributable to their introns^10^. Why introns have evolved to such lengths within genes implicated in neuronal functions remains puzzling, especially considering the tremendous energetic cost and topological challenges associated with the replication and transcription of megabase-long genes. Interestingly, in human and mice, genes expressing the longest transcripts are enriched in pro-longevity genes^12^ and tissue aging has been linked to a progressive decrease in the expression of long genes, particularly in the brain^12,13^.

Disruption of the 3D genome as well as DNA damage and mutations affecting long neuronally expressed genes have all been linked to various neurodevelopmental and neuropsychiatric disorders, including schizophrenia and autism. During prenatal development, genes implicated in schizophrenia cluster in networks of highly expressed genes^14^. Concordantly, genetic variants associated with schizophrenia and bipolar disorder risk have been found to be enriched in regulatory elements that partake in cell type-specific contacts with their target genes, which in turn display coordinated expression in neuronal cells and encode regulators of neuronal connectivity and chromatin remodeling^15–18^. Similarly, chromatin loops acquired during human cortical development have been associated with genomic risk loci for autism identified in genome-wide association studies^6^. Genes implicated in autism and schizophrenia are also hotspots of DNA double-strand break formation in neuronal cells, possibly related to DNA torsional stress associated with the topological reorganization of these loci during neurodevelopment^19,20^.

A growing body of evidence indicates that chromatin-associated RNAs (caRNAs) play key roles in regulating genome function and 3D organization, including in neuronal cells^21,22^. caRNAs can neutralize the positive charges on histone tails and compete with DNA binding, preventing chromatin compaction in a sequence-independent manner, as exemplified by CoT-1 RNA, which comprises a diverse mix of repetitive transcripts including LINE-1 RNA^23,24^. Besides several well-studied ‘architectural’ long non-coding RNAs (lncRNAs) such as *MALAT1*, *XIST*, and *NEAT1*, other caRNAs exert regulatory roles in gene expression, RNA processing, heterochromatin assembly, and segregation of active (A) and inactive (B) chromatin^25^. By interacting with RNA-binding proteins (RBPs), caRNAs have been proposed to form mesh-like structures that serve as scaffolds for chromosome territories (CTs) and/or support transcriptionally active compartments within them^26^. Moreover, various nuclear processes involving caRNAs occur within subnuclear condensates formed through phase separation. RNA—especially long, nascent RNA and lncRNAs—can trigger phase separation through a combination of electrostatic forces and RBP binding^27–29^. Indeed, it has been shown that newly synthesized RNA can induce tunable, transcription-dependent microphase separation by nucleating at transcriptionally active sites across the nucleus (named transcription pockets or microgels)^30,31^. It is increasingly believed that the ability of caRNAs and RBPs to form mesh-/gel-like structures underlies the formation of the so-called nuclear matrix, a term first introduced almost five decades ago^32,33^. Factors implicated in the formation of this structure include the splicing factors RBFOX2 and TDP-43 and classical nuclear matrix proteins such as MATR3 and heterogeneous nuclear ribonucleoproteins (HNRNPs), including HNRNPU (also known as scaffold attachment factor A, SAF-A)^26,31^. However, the precise RNA and protein constituents of the nuclear matrix remain poorly defined. It is also unclear whether the repertoire of scaffold RNAs and proteins varies across cell types and how this RNA-protein meshwork mechanistically shapes 3D genome organization. Of note, mutations in SAF-A/HNRNPU have been associated with neurodevelopmental disorders including autism^34^, suggesting an important role of the nuclear matrix in neurodevelopment.

To illuminate the interplay between caRNAs, the nuclear matrix, and the 3D genome in neuronal cells, here we perform multi-omic profiling of the 3D genome, RNA-DNA contactome, and transcriptome across in vitro neuronal differentiation, in the frame of the sixth edition of the Functional ANnoTation Of the Mammalian genome (FANTOM6) project. We identify and characterize a novel group of intronic RNAs that are transcribed from ultra-long, neuronally expressed genes and that accumulate in neuronal nuclei while engaging in genome-wide trans interactions with multiple DNA loci. We thoroughly characterize the DNA loci from which these Trans-Interacting Intronic RNAs (TIRs) are produced and visualize the spatial organization of the most abundant TIRs in neurons, revealing that they form elaborate pan-nuclear patterns that extend through a large portion of the nucleus. Finally, we show that TIRs are important for the co-regulation of gene expression in TIR-contacted genomic regions and provide evidence suggesting a possible role of TIRs in the pathogenesis of neurodevelopmental and neuropsychiatric disorders. Our study establishes the first atlas of the RNA-DNA contactome during human neurogenesis and highlights TIRs as novel structural constituents of the nucleus of neuronal cells. Our findings support a model in which RNA species generated by transcription, but not translated into proteins, do not merely reflect gene activity but actively shape the nuclear context in which future transcription occurs.

## Results

### Multi-omic profiling of human iPSC in vitro differentiated to cortical neurons

One of the aims of the FANTOM6 consortium is to unravel the interplay between coding and non-coding RNAs and genome structure-function during cell differentiation. Towards this goal, we performed multi-omic profiling of human male induced pluripotent stem cells (iPSC), iPSC-derived neural stem cells (NSC), and cortical neurons (NEU) (**Experimental Methods**, sections 1 and 2), using the following assays: (i) RNA and DNA Interacting Complexes Ligated and sequenced (RADICL-seq)^35^ to profile RNA-DNA contacts; high-throughput chromosome conformation capture (Hi-C)^36^ to measure DNA-DNA contacts; Genomic Loci Positioning by Sequencing (GPSeq)^37^ to map genome-wide radial positions in the nucleus; pseudo-bulk single-cell Assay for Transposase-Accessible Chromatin using sequencing (scATAC-seq)^38^ for measuring chromatin accessibility; total RNA sequencing (RNA-seq); and Cap Analysis of Gene Expression (CAGE)^39^ for promoter identification and gene expression profiling (**Fig. 1a and Experimental Methods**, sections 3–8).

**Figure 1.**
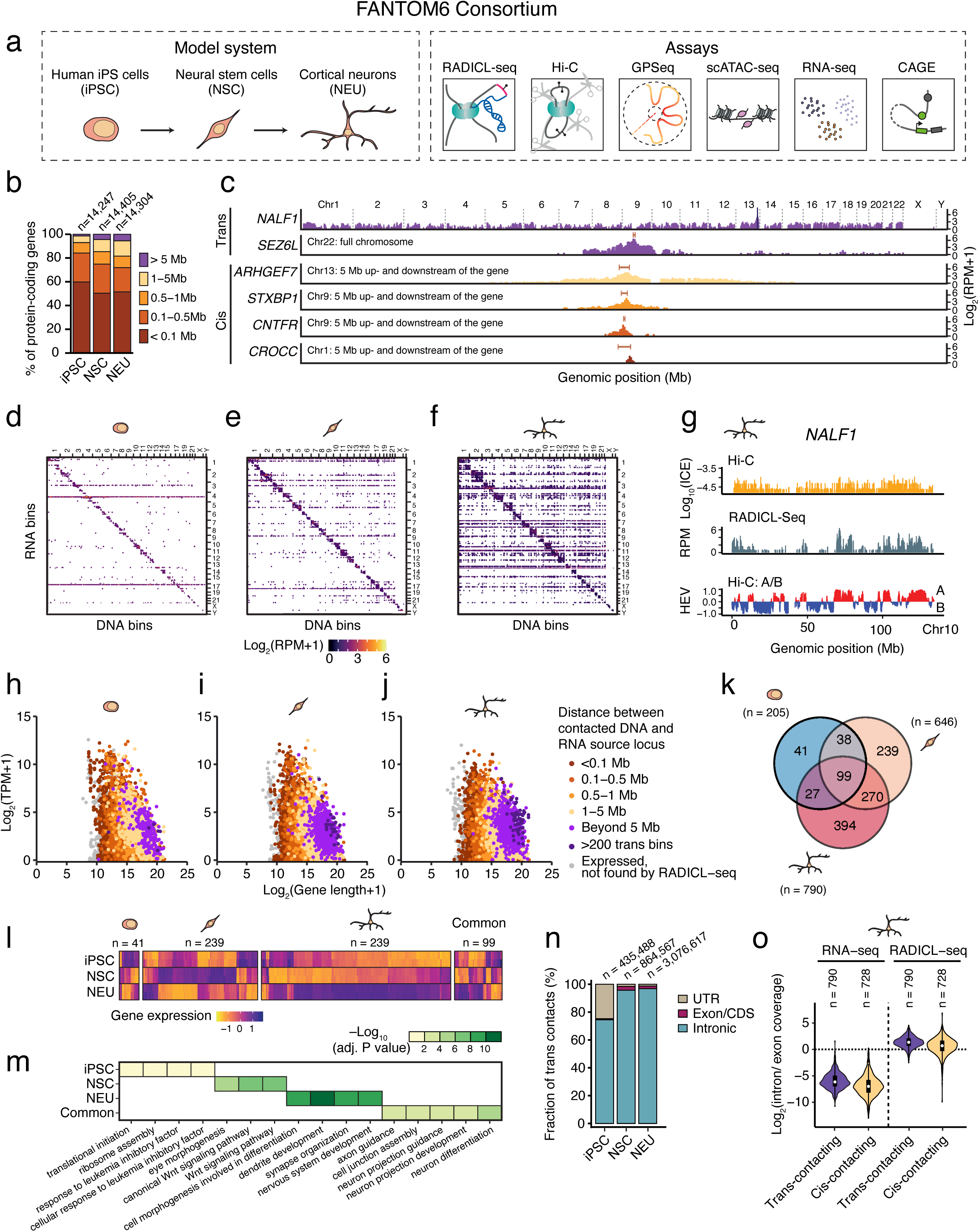
Identification of cis– and trans-contacting RNAs by RADICL-seq. (**a**) Scheme of the model system (left box) and datasets generated from them (right box) in the frame of the sixth edition of the Functional ANnoTation Of the Mammalian genome project (FANTOM6), which were analyzed in this study. (**b**) Proportion of protein-coding genes producing RNAs that engage in RNA-DNA contacts as detected by RADICL-seq, for different genomic distances between the RNA source locus and the contacted loci. *n*, number of protein-coding genes. (**c**) Representative RNA cis and trans contact profiles (*NALF1* and *SEZ6L*: 100 kb resolution; all other genes: 25 kb resolution) revealed by RADICL-seq (trans defined as any contact ≥ 5 Mb from the source gene). Red segments mark the position of the RNA source gene. RPM, reads per million. (**d-f**) RNA-DNA contact matrices (100 kb resolution) for iPSC (d), NSC (e) and NEU (f). Only trans contacts of RNA from protein-coding genes are shown. (**g**) Top track: profile of inter-chromosomal DNA-DNA contacts along chromosome (chr) 10 using the *NALF1* gene locus on chr13 as viewpoint, as measured by Hi-C in NEU. Middle track: trans RNA-DNA contacts of *NALF1* RNA along chr10, detected by RADICL-seq in NEU. Bottom track: A/B chromatin compartments identified along chr10, based on Hi-C data in NEU. For all tracks, the resolution is 100 kb. ICE, iterative correction and eigenvector decomposition. HEV, first Hi-C eigenvector. (**h-j**) Relationship between the length (in base pairs) and the expression level of all human protein-coding genes, in the three cell types profiled by RADICL-seq. Each dot represents a gene. Dots are colored based on whether the RNA transcribed from them was detected by RADICL-seq and on the farthest distance between the source gene and the contacted DNA loci. TPM, transcripts per million. (**k**) Venn diagram showing the number (*n*) of protein-coding genes producing trans-contacting RNAs in the three cell types profiled by RADICL-seq. (**l**) Gene expression levels measured by RNA-seq for cell type-specific and common protein-coding gene sets producing trans-contacting RNAs. Each column in the RNA-seq heatmap panel represents one gene. Gene expression (DESeq rlog) was scaled gene by gene across iPSC, NSC, and NEU. (**m**) Significant gene ontology terms for the genes included in the plot shown in l. (**n**) Proportion of trans contacts mediated by RNAs derived from protein-coding genes, for different segments of the gene body to which the RNA sequence aligns. UTR, untranslated region, either at the 5′ or 3′ end of the gene. CDS, coding DNA sequence. *n*, number of trans RNA-DNA contacts detected by RADICL-seq. (**o**) Ratio between the intronic and exonic maximum RNA coverage of the trans-contacting and cis-contacting genes (as control) expressed at comparable levels in NEU, as calculated from RNA-seq (left) or RADICL-seq (right) data. *n*, number of genes. Violins extend from minimum to maximum, boxplots extend from the 25^th^ to the 75^th^ percentile, white dots represent the median, whiskers extend from –1.5×IQR to +1.5×IQR from the closest quartile. IQR, inter-quartile range. A link to the Source Data and code to regenerate the plots displayed in this figure is provided in the Data Availability and Code Availability statements.

We generated libraries from two biological replicates per cell type (iPSC, NSC, NEU) and assay (except for scATAC-seq, which was performed on one batch of single cells) and sequenced them on different Illumina platforms (**Supplementary Table 1**). We analyzed RADICL-seq, GPSeq, and CAGE data using custom pipelines, and leveraged state-of-the-art pipelines for Hi-C, RNA-seq, and scATAC-seq (**Computational Methods**). We confirmed that biological replicates were highly correlated (**Supplementary Fig. 1a-e**); hence we merged them for subsequent analyses.

To confirm that iPSCs were properly differentiated to neurons, we identified differentially expressed genes (DEG) and performed gene ontology (GO) analysis on the RNA-seq datasets (**Computational Methods**, sections 1 and 2). Among genes upregulated in NEU compared to iPSC, we found many genes involved in synapse organization, regulation of neurotransmitter transport, and other neuron-specific processes (**Supplementary Fig. 1f**). Conversely, genes downregulated in NEU compared to iPSC included genes involved in cell division and chromatin organization (**Supplementary Fig. 1g**), which is expected as cells exit the cell cycle during neuronal differentiation. Proper iPSC differentiation to neurons was further confirmed by immunofluorescence staining for well-established markers of neuronal cells, including NeuN, MAP2 and beta-Tubulin (**Supplementary Fig. 1h, i and Experimental Methods**, section 9). Altogether, these results confirmed the validity of our model system, allowing us to proceed to in-depth analyses of all the other datasets.

### RADICL-seq uncovers multiple RNAs that contact DNA far from their locus of origin

We first analyzed RADICL-seq data by binning the identified RNA-DNA contacts in 100 kilobase (kb) genomic windows (bins) and generating RNA-DNA contact matrices (**Computational Methods**, section 3). We classified each RNA-DNA contact as cis or trans, depending on whether the RNA fragment contacted a DNA locus within 5 megabase (Mb) (on either side) from the RNA-transcribing locus (‘cis contacts’), or farther along the same chromosome or on other chromosomes (collectively named ‘trans contacts’). We chose a 5 Mb threshold to exceed the typical size of TADs (1–2 Mb) and thus avoid calling trans contacts within the same TAD containing the RNA-producing locus (hereafter, ‘source’).

In all three cell types, ∼99% of all RADICL-seq reads originated from RNAs transcribed from either protein-coding (pc) genes or from lncRNA genes (**Supplementary Fig. 2a**). Most of the RNA-DNA contacts occurred in cis, in the immediate vicinity of their source (**Fig. 1b, c—**excluding contacts involving the lncRNA *MALAT1***—**and **Supplementary Fig. 2b-j**). Among trans contacts, more than 60% involved *MALAT1*, which formed genome-wide contacts in all the three cell types analyzed (**Supplementary Fig. 2e, g, i**). A few other lncRNAs, including *GAS5, RMRP,* the splicing factor *RNVU1-7*, and the small nucleolar RNAs (snoRNAs) *U8* and *SNHG14* also formed genome-wide contacts, predominantly in iPSC (**Supplementary Fig. 2e, g, i**).

Excluding contacts involving *MALAT1,* most of the remaining trans contacts involved RNA transcribed from pc genes (**Fig. 1d-f and Supplementary Fig. 2c, f, h, j**). Whereas only a small fraction of RNAs transcribed from pc genes engaged in trans contacts in iPSC (1.4%, *n*=205), the proportion significantly increased in NSC (4.5%, *n*=646) and in NEU (5.5%, *n*=790) (**Fig. 1b**). Among trans-contacting RNAs produced by pc genes in NEU, we identified dozens of transcripts that engage in trans contacts with DNA loci on different chromosomes than the one carrying their source locus (**Supplementary Fig. 2f, h, j**). Moreover, five of the top-10 genes producing trans-contacting RNAs in NEU (*NALF1*, *LSAMP*, *CNTNAP2*, *PCDH9*, and *NRG3*) are expressed at moderate-to-high levels almost exclusively in the brain (**Supplementary Fig. 3a-e**). Notably, trans contacts did not simply recapitulate long-range Hi-C contacts but rather displayed a unique pattern with more clearly distinguishable peaks and valleys compared to Hi-C profiles, mainly overlapping with the A chromatin compartment (**Fig. 1g and Computational Methods**, section 4).

### Trans-contacting RNAs originate from highly expressed, cell type-specific, long pc genes

We then focused on characterizing the trans-contacting RNAs that originate from pc genes. In all three cell types, those RNAs were transcribed from relatively long genes with cell type-specific expression and function, as revealed by DEG and GO analyses (**Fig. 1h-m, Supplementary Fig. 3f-i**). Other cell types profiled by RADICL-seq within the FANTOM6 consortium could also be separated into distinct clusters based on both the repertoire of genes producing trans-contacting RNAs (**Supplementary Fig. 4a**) and on the genomic regions contacted by these RNAs (**Supplementary Fig. 4b, c**). In all the cell types analyzed, the source genes of trans-contacting RNAs were consistently longer and more highly expressed than the average length of pc genes (**Supplementary Fig. 4d, e and Computational Methods**, section 5). However, we only observed these widespread patterns of trans contacts in NEU cells (**Supplementary Fig. 4f**). The source genes of trans-contacting RNAs showed moderate-to-high expression levels (**Supplementary Fig. 4e**). Moreover, pc genes with comparable expression levels to the top-10 source genes, whose RNAs were not detected by RADICL-seq, were linked with housekeeping or mitochondrial functions based on GO analysis, as opposed to cell type-specific functions (**Supplementary Fig. 5a-e**). These results indicate that, across all the cell types profiled by RADICL-seq, but more prominently in NEU, a set of cell type-specific, moderate-to-highly expressed, and longer-than-average pc genes generate RNAs that engage in RNA-DNA contacts with multiple regions on different chromosomes.

### Trans-contacting RNAs are mainly composed of intronic sequences

To further investigate the nature of the identified trans-contacting RNAs, we annotated the RADICL-seq reads and found that more than 95% of the reads from NSC and NEU aligned to introns (**Fig. 1n**). To understand whether these reads originate from spliced-out introns or from intronic segments of nascent RNA or pre-mRNA, we aligned all the RADICL-seq and RNA-seq reads from NSC and NEU to the corresponding source genes and computed the intron-over-exon coverage ratio (**Computational Methods**, section 6). Despite the presence of many intronic reads in our RNA-seq data, the coverage was higher for exons (**Fig. 1o**). In contrast, the gene coverage of RADICL-seq reads was slightly skewed towards intronic sequences (**Fig. 1o**). Of note, the trans-contacting RNAs identified by RADICL-seq in other cell lines also mapped predominantly to introns (**Supplementary Fig. 5f**). Importantly, we did not observe a strong enrichment of CAGE peaks in the introns of the source genes producing trans-contacting RNAs (**Supplementary Fig. 5g-j and Computational Methods**, section 5). This suggests that the observed pattern of RADICL-seq reads is unlikely caused by alternative transcription start sites.

In sum, these results indicate that trans-contacting RNAs produced by pc genes are predominantly intronic sequences, and that the abundance of pc gene-derived trans-contacting RNAs significantly increases during neuronal differentiation. We refer to these trans-contacting RNAs as trans-contacting intronic RNAs, or TIRs, and define as high-confidence TIRs those interacting with 200 or more 100 kb genomic bins (**Fig. 1h-j**). By applying this threshold, in NEU we identified 55 high-confidence TIRs (**Supplementary Table 2**), which we further characterized in depth.

### The number of TIR trans contacts does not linearly scale with the length of introns

We then analyzed the relationship between intron length and the number and diversity of trans contacts of the top-55 TIR-producing genes identified in NEU. To this end, for each of those genes separately, we assessed the relationship between the length of each individual intron of a given gene and the number of trans contacts formed by the TIRs transcribed from the same gene (**Computational Methods**, section 7). We found that TIRs derived from longer introns tend to engage in disproportionately more trans contacts per nucleotide as compared to TIRs originating from shorter introns (**Fig. 2a and Supplementary Fig. 6a**). In contrast, the diversity of genomic regions contacted by the TIRs derived from the introns of the top-55 TIR source genes rapidly saturated for increasing intron lengths (**Fig. 2b and Supplementary Fig. 6a**). This suggests that trans contacts are not distributed randomly across the genome but are constrained to a defined repertoire of genomic loci.

**Figure 2.**
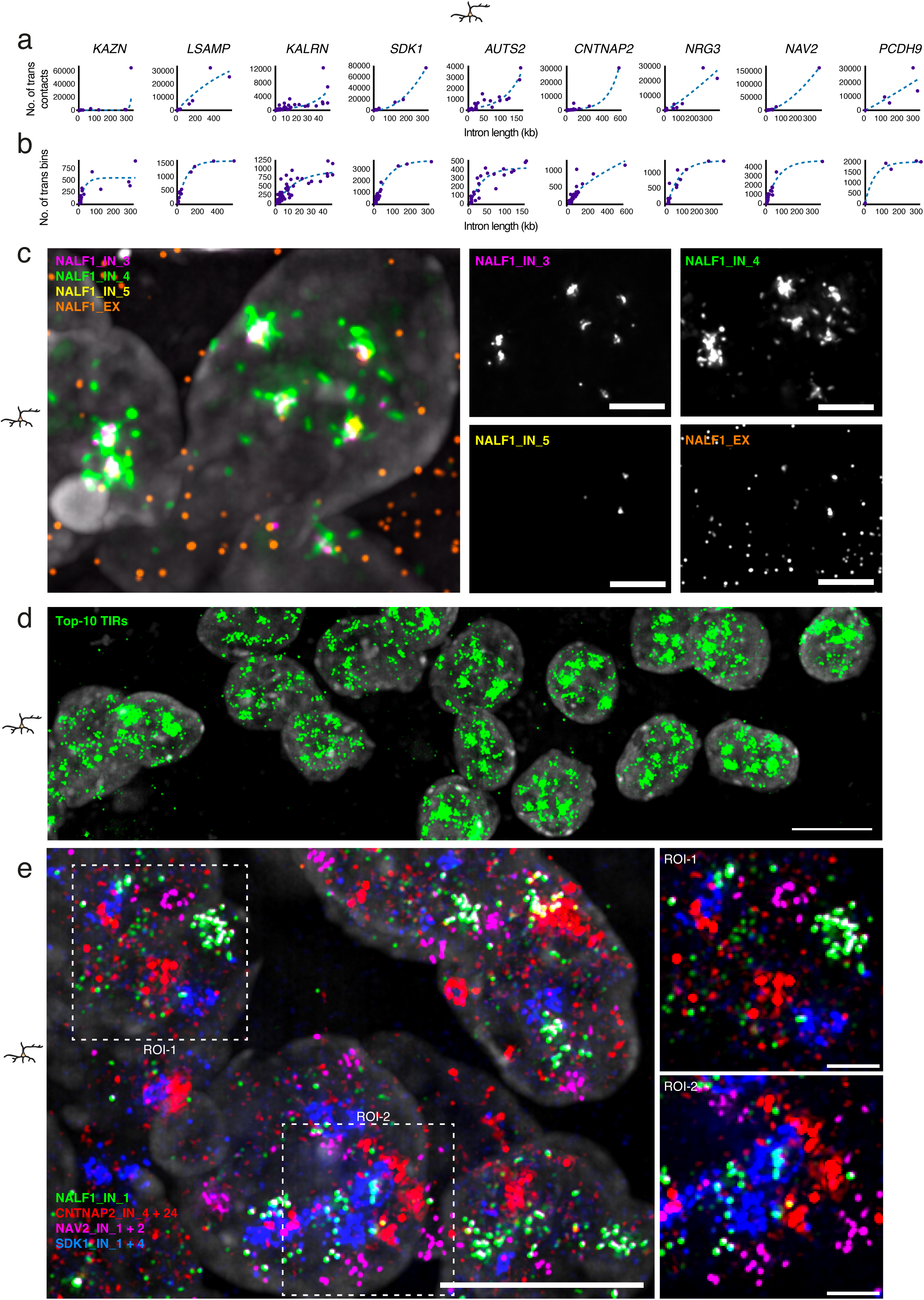
Trans-contacting Intronic RNAs (TIRs) form an intricate nuclear meshwork. (**a**) Relationship between intron length and the total number of trans contacts formed by the RNAs transcribed from the indicated genes in NEU, as detected by RADICL-seq. The top-10 TIR source genes are shown. kb, kilobases. (**b**) As in (a) showing the relationship between intron length and the number of distinct 100 kb genomic bins contacted by TIRs. (**c**) Maximum intensity projection of a z-stack widefield microscopy image exemplifying the nuclear pattern detected in NEU by RNA FISH using probes targeting all the exons (EX) or various intronic (IN) regions of *NALF1*. Gray, DNA stained by Hoechst 33342. Scale bars, 5 μm. The insets on the right show the signal produced by each probe separately. (**d**) As in (c) showing the nuclear pattern of TIRs originating from the top-10 TIR source genes detected in NEU, using a pool of the intron-targeting probes shown in **Supplementary** Fig. 6c with the exclusion of CAMTA1_IN probes and labelling all the probes in the pool with the same color. Scale bar, 10 μm. (**e**) As in (c) using RNA FISH probes targeting different intronic regions of *NALF1*, *SDK1*, *CNTNAP2*, and *NAV2*. Scale bars, 10 μm (large image) and 2 μm (ROI insets). ROI, region of interest. See **Supplementary Table 3** for the list of oligos composing the FISH probes shown in this figure. The images shown in (c-e) were deconvolved using Deconwolf (https://deconwolf.fht.org/). A link to the Source Data and code to regenerate the plots displayed in this figure is provided in the Data Availability and Code Availability statements.

### TIRs form large ‘clouds’ in the nucleus of neuronal cells

Next, we sought to investigate how TIRs are spatially distributed in the nucleus. To visualize the spatial distribution of TIRs in single cells, we performed RNA fluorescence in situ hybridization (RNA FISH) (**Experimental Methods**, section 10) leveraging an updated version of the iFISH probe design pipeline that we previously developed^40^ (https://ifish4u.fht.org/) and using our Deconwolf software^41^ (https://deconwolf.fht.org/) for image deconvolution and dot picking (**Computational methods,** section 8). We designed RNA FISH probes targeting either intronic or exonic regions of selected TIRs displaying the most abundant and widespread trans contacts in NEU, as measured by RADICL-seq (**Supplementary Fig. 6b, c and Supplementary Table 3**). As controls, we designed RNA FISH probes targeting 11 pc genes with expression levels comparable or higher than those of the selected TIR source genes, but whose transcripts did not engage in trans contacts (**Supplementary Table 3**). When probing for TIR intronic sequences in NEU, we observed many RNA FISH signals (fluorescence dots) in the nucleus, typically forming 1–2 large dot clusters (‘clouds’) per nucleus, most likely marking the location of the transcribing source loci (**Fig. 2c, Supplementary Fig. 7a-c, Supplementary Fig. 8a, b, Supplementary Fig 9a-c, and Supplementary Fig. 10a-e**). In contrast, probing for the introns of TIR control genes in NEU typically yielded only 1–2 individual dots per nucleus, without any detectable dot cloud (**Supplementary Fig. 11a-e**). Furthermore, RNA FISH with probes targeting the exons of the same TIR source or control genes yielded numerous dots, including nuclear dots (some of which co-localized with intron dot clouds) as well as multiple dots in the cytoplasm, representing exported mature mRNAs (**Fig. 2c and Supplementary Fig. 7-10**). Notably, despite the fact that the NEU-specific TIR source gene –CNTNAP2 – was found expressed in NSC at comparable levels as in NEU, its long introns did not form as large dot clouds in NSC cells as in NEU (**Supplementary Fig. 12a-c**).

For some of the TIRs analyzed, in addition to the FISH dot clouds, we also observed multiple intronic dots scattered throughout the nucleus in NEU, potentially representing spliced-out intronic fragments (**Fig. 2c and Supplementary Fig. 7-10**). Indeed, RNA FISH with probes targeting both the exons and one selected intron of the same TIR source gene revealed almost no colocalization between the sparse intronic dots and exonic dots (**Fig. 2c and Supplementary Fig. 7-10**). Furthermore, while the dots corresponding to two different introns of the same TIR source gene largely overlapped within the same dot cloud, the sparse intronic dots originating from the same TIR source gene did not colocalize (**Fig. 2c and Supplementary Fig. 7-10**). Of note, the number of intronic and exonic FISH dots per cell were not correlated across different TIR source genes (Pearson’s correlation coefficient, PCC: –0.015) (**Supplementary Fig. 12d**). These results indicate that the sparse intronic dots detected by RNA FISH most likely correspond to spliced-out introns. Importantly, when considering the top-10 TIRs detected by RADICL-seq in NEU, we found a strong positive correlation (PCC: 0.8) between the number of intronic RNA FISH dots per cell and the corresponding number of RADICL-seq contacts (**Supplementary Fig. 12e**), indicating high consistency between RNA FISH and RADICL-seq results.

We then visualized all the top-10 TIRs detected in NEU by pooling together all the RNA FISH intronic probes and labeling them with the same fluorophore (**Supplementary Table 3**). Despite the fact that each of these probes targets only a small fraction of the introns of the corresponding TIR source gene (**Supplementary Fig. 6c**), we observed a widespread localization pattern occupying a substantial portion of the nucleus of NEU cells (**Fig. 2d**). By labeling distinct TIRs in separate colors, the TIR clouds originating from different source genes partially colocalized, with highly variable localization patterns between individual cells (**Fig. 2e and Supplementary Fig. 13**). Interestingly, when staining four consecutive segments of a single, exceptionally long (∼600 kb) intron of *NALF1* with four different fluorophores (**Supplementary Fig. 14a**), we found that many of the sparse FISH dots (i.e., not included in dot clouds) were often co-localized, with numerous 2-, 3-, or even 4-color co-localization events detected (**Supplementary Fig. 14b-f, Supplementary Fig. 15a-f, and Computational methods,** section 8). This indicates that these sparse intronic dots observed correspond to exceptionally long RNA molecules, given that each single-color probe targets ∼120–200 kb segments of the intron. We note that the detection sensitivity of the third segment of the intron was lower compared to the other segments (**Supplementary Fig. 15f**), possibly causing an underestimation of the co-localization rate.

Altogether, these results demonstrate that, in NEU cells, long neuronally expressed genes generate exceptionally long intronic RNAs that do not get rapidly co-transcriptionally degraded but, instead, accumulate in the nucleus, forming a rich variety of localization patterns. Large clouds of intronic RNAs most likely surround the corresponding source loci and represent both nascent and partially spliced transcripts, as well as (partially) spliced(-out) introns. In addition to these large dot clouds, multiple sparse intronic dots are also detected, possibly representing spliced-out (often very long) introns that ‘escaped’ co-transcriptional degradation. Notably, the total mass of intronic RNAs produced by just the top-10 TIR source genes detected in NEU seem to occupy a large fraction of the volume of the nucleus of these cells.

### TIR clouds do not overlap with splicing speckles

Since TIR source genes are typically very long and correspond to developmentally upregulated genes composed of long introns and (often) numerous exons, thus requiring extensive splicing, we wondered whether the observed TIR clouds are proximal to or overlap with nuclear speckles, which represent the main splicing hub in the nucleus^42,43^. To test this, we first leveraged our RADICL-seq data to identify contacts between the lncRNA, *MALAT1*—a speckle marker^44,45^—and the top-10 TIR source genes identified in NEU. We computed a speckle proximity score based on the *MALAT1* RNA contact frequency calculated from RADICL-seq data (**Computational Methods**, section 3). The top-10 TIR source genes were marked by low scores (**Fig. 3a, b**), suggesting that they do not frequently associate with speckles. To corroborate this finding, we performed RNA FISH targeting *MALAT1* RNA together with all the top-10 TIRs identified in NEU, by pooling together all the RNA FISH intronic probes and labeling them with the same fluorophore (**Supplementary Table 3**). This confirmed the absence of obvious co-localization between *MALAT1* RNA and the TIR dot clouds (**Fig. 3c and Supplementary Fig. 16a-d**), suggesting that TIR clouds occupy a nuclear neighborhood distinct from speckles.

**Figure 3.**
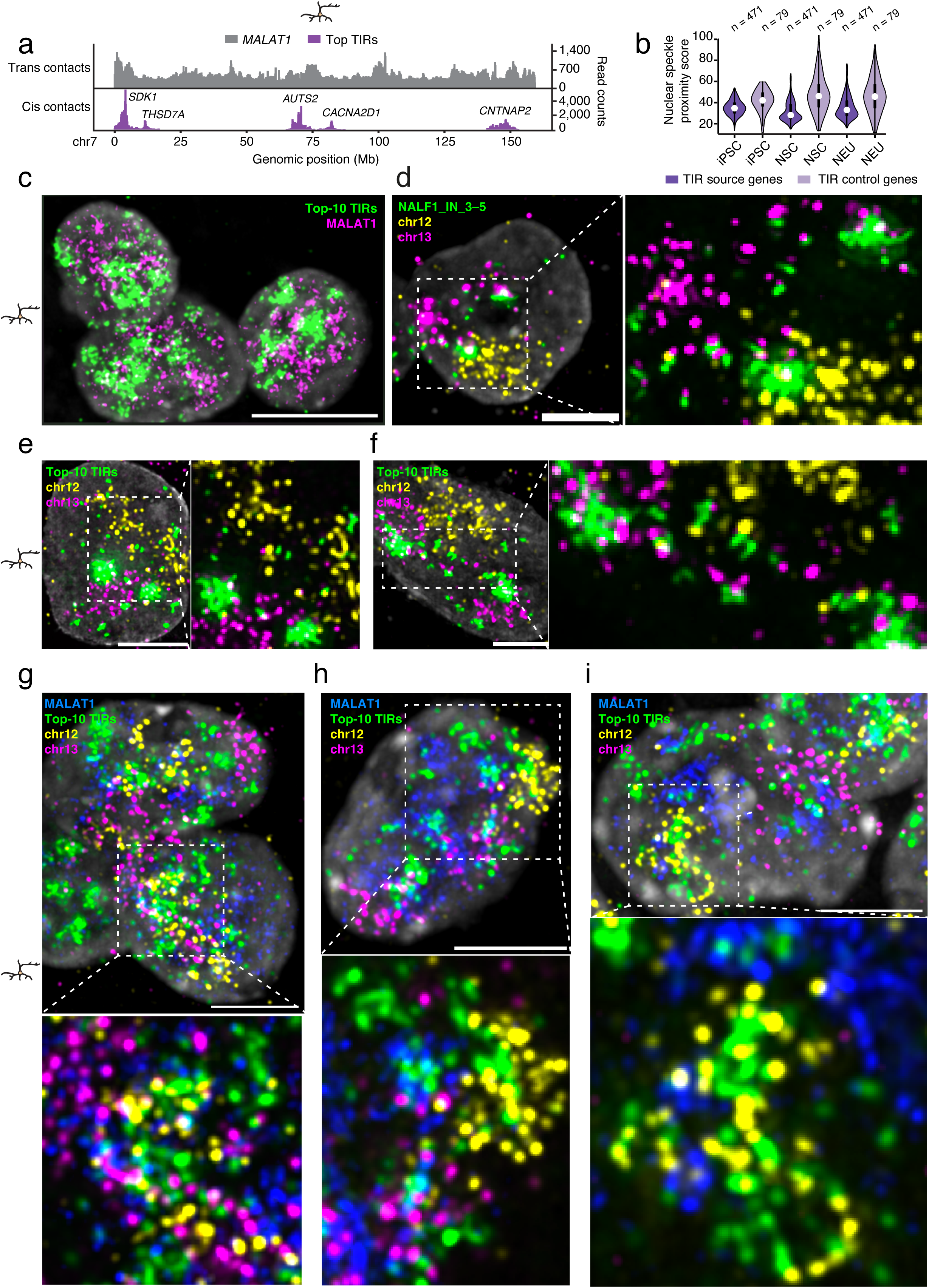
TIR clouds are distinct from nuclear speckles and extend beyond the chromosome territory of the source gene. (**a**) Profiles (100 kb resolution) along chromosome (chr) 7 of *MALAT1* RNA trans contacts (top track) and of cis contacts for the indicated TIRs (bottom track) identified by RADICL-seq in NEU. (**b**) Distribution of the nuclear speckle proximity score inferred from the normalized (based on reads per million) *MALAT1* trans contacts for the top-55 TIR source genes and an equal number (55) of TIR control genes. *n*, number of 100 kb bins overlapping with the gene body of top-55 TIR source gene and of the 55 TIR control genes. Violins extend from minimum to maximum, boxplots extend from the 25^th^ to the 75^th^ percentile, white dots represent the median, whiskers extend from –1.5×IQR to +1.5×IQR from the closest quartile. IQR, inter-quartile range. (**c**) Maximum intensity projection of a z-stack widefield microscopy image exemplifying the pattern of *MALAT1* RNA (marking nuclear speckles, magenta) and of intronic RNAs (green) originating from the top-10 TIR source genes, as detected by RNA FISH in NEU. Intronic RNAs were detected using a pool of the intron-targeting probes shown in **Supplementary** Fig. 6c with the exclusion of CAMTA1_IN probes and labelling all the probes in the pool with the same color. Gray, DNA stained by Hoechst 33342. Scale bar, 10 μm. (**d**) Maximum intensity projection of a z-stack widefield microscopy image exemplifying the spatial distribution of *NALF1* TIRs visualized by RNA FISH with respect to the chromosome territory (CT) of chr12 and chr13 (where *NALF1* is located) visualized by chromosome spotting DNA FISH probes. Scale bar, 5 μm. The white-dash square region is magnified on the right. (**e, f**) As in (d) but using the pool of RNA FISH probes targeting the top-10 TIRs shown in (c) instead of the NAL1_IN probes. (c). (**g-i**) As in (e, f) with the addition of an RNA FISH probe targeting *MALAT1* RNA. See **Supplementary Table 3** for the list of oligos composing the FISH probes shown in this figure. The images shown in (c-g) were deconvolved using Deconwolf (https://deconwolf.fht.org/). A link to the Source Data and code to regenerate the plots displayed in this figure is provided in the Data Availability and Code Availability statements.

### TIR source genes undergo structural reorganization during neuronal differentiation

Prompted by these observations, we sought to further characterize the chromatin neighborhood of the TIR clouds. Given their shape, we wondered whether the clouds are confined to their chromosome territory (CT) of origin, similarly to *XIST* RNA on the inactive X chromosome^46,47^. To explore this, we performed combined DNA&RNA FISH using RNA FISH probes against *NALF1* intronic RNA and chromosome-spotting DNA FISH probes^40^ targeting chromosome (chr) 13, which harbors the *NALF1* locus (**Supplementary Table 3**). For comparison, we also visualized the CT of chr12 (**Supplementary Table 3**), which, in NEU, engages in extensive trans contacts with the top-55 TIR genes, while it does not contain any of the top-10 TIR source genes (hence, it is devoid of cis contacts from the top-10 TIR source genes identified in NEU). In NEU, *NALF1* TIR clouds were not confined to the *NALF1* locus and often extended beyond the chr13 CT, typically showing minimal overlap with it (**Fig. 3d-f and Supplementary Fig. 17a-c**). In some nuclei, the *NALF1* intronic cloud was located considerably away from the core of chr13 CT and, interestingly, some of the sparse *NALF1* TIR dots colocalized with chr12 (**Fig. 3d-f and Supplementary Fig. 17a-c**). We then visualized all the top-10 TIRs identified in NEU together, co-staining for both chr12 and chr13 (note that chr13 carries three TIR source genes, while chr12 none). This revealed a high extent of mingling between the CT of chr12 and the cloud formed by the top-10 TIRs; this mingling was much less pronounced in the case of chr13 (**Fig. 3g-i and Supplementary Fig. 18a-d**). Of note, the top-10 TIRs cloud and *MALAT1* RNA did not co-localize (**Fig. 3g-i and Supplementary Fig. 18a-d**), further confirming that TIRs and speckles are spatially separated.

To further assess the chromatin environment of TIR clouds, we examined Hi-C maps from iPSC, NSC, and NEU, assessing how the chromatin contacts of TIR source genes change during neuronal differentiation (**Computational Methods**, section 4). We observed that a substantial portion of the distal contacts of TIR source genes (but not of TIR control genes) with DNA was lost within a 30 Mb region around these loci (excluding the immediate vicinity of each locus) during differentiation (**Supplementary Fig. 19a-l and Supplementary Fig. 20a**). Moreover, the Hi-C eigenvector values of TIR source genes tended to increase from iPSC to NEU, suggestive of i) weakening of the connectivity with the surrounding B compartment; ii) a shift from B to A; or iii) occasionally, strengthening of the connectivity within the A compartment (**Supplementary Fig. 20b, c**). This is reminiscent of what has been previously described for transcriptional loops (TLs)^48^ and gene melting^49^, which represent phenomena associated with highly expressed long genes. Indeed, the gene body of TIR source genes, visualized by DNA FISH, appeared to have an ‘extended’ configuration in many cells (**Supplementary Fig. 20d-f and Experimental Methods**, section 10), similarly to what has been reported for TL-forming genes such as the thyroglobulin gene^48^, even though the extended configuration was less pronounced in the case of TIR genes. Moreover, in contrast to the structural changes associated with TLs^48^ and gene melting^49^, we observed well-defined TAD structures around the TIR source loci in iPSC and even more pronounced in NEU (**Supplementary Fig. 21a-h**). This discrepancy might be explained by the much more modest expression of the TIR source genes in comparison to TL-forming genes.

We also noticed that the clearly patterned structure of Hi-C contacts in the vicinity of TIR source genes loci often resembled the RADICL-seq profile of the same locus (**Supplementary Fig. 22a-i**). To quantitatively compare RADICL-seq and Hi-C contact maps, we calculated the RADICL-seq residues in NEU and correlated them with the Hi-C observed-over-expected (O/E) values in the three cell types (**Supplementary Fig. 22a-j and Computational Methods**, section 9). As expected, the NEU RADICL-seq residues were positively correlated with the Hi-C O/E values from the same cell type (**Supplementary Fig. 22k, l**), suggesting that the chromatin structure in the vicinity of the TIR source genes demarcates the RNA-DNA connectivity detected by RADICL-seq. Of note, for many TIR source genes, we found that RNA-DNA contacts extended beyond multiple TAD distances, which was not the case for TIR control genes (**Supplementary Fig. 22m**). Altogether, these results indicate that the formation of intronic RNA clouds around TIR source genes is accompanied by a profound reorganization of both the local chromatin topology and the long-range contactome of those genes, as they become upregulated during neuronal differentiation. We speculate that the long introns of TIR source genes might play a role in this process.

### TIRs are enriched in RNAs bound to the nuclear matrix protein, HNRNPU and in the insoluble portion of the nucleus

The spatial pattern and nuclear abundance of the top TIRs found in NEU prompted us to consider whether these RNAs might play a role in the formation of the nuclear matrix^22,26,50^. This hypothesis is corroborated by recent reports demonstrating that intronic RNAs are enriched in the chromatin-depleted nuclear insoluble fraction, which is also enriched in protein constituents of the nuclear matrix, such as HNRNPU^26^. To test the possible association between TIRs and the nuclear matrix, we reanalyzed the results of an RNA pull-down experiment in which formaldehyde crosslinking followed by ribonucleoprotein immunoprecipitation (fRIP-seq) was applied to identify RNAs bound to HNRNPU (SAF-A) in neuronal cells differentiated from neuroepithelial stem cells^51^ (**Computational Methods**, section 10). We found that TIRs were significantly enriched among HNRNPU-bound RNAs in those cells, suggesting a propensity for TIRs to engage in ribonucleoprotein complexes with HNRNPU (**Fig. 4a**). Of note, among HNRNPU-bound RNAs, we also detected a significant enrichment in genes associated with autism, intellectual disability, and epilepsy, suggesting shared regulatory relevance. As expected, the enrichment of TIRs among HNRNPU-bound RNAs was stronger in neurons compared to neuroepithelial stem cells, consistent with TIR source genes becoming upregulated in NEU cells.

**Figure 4.**
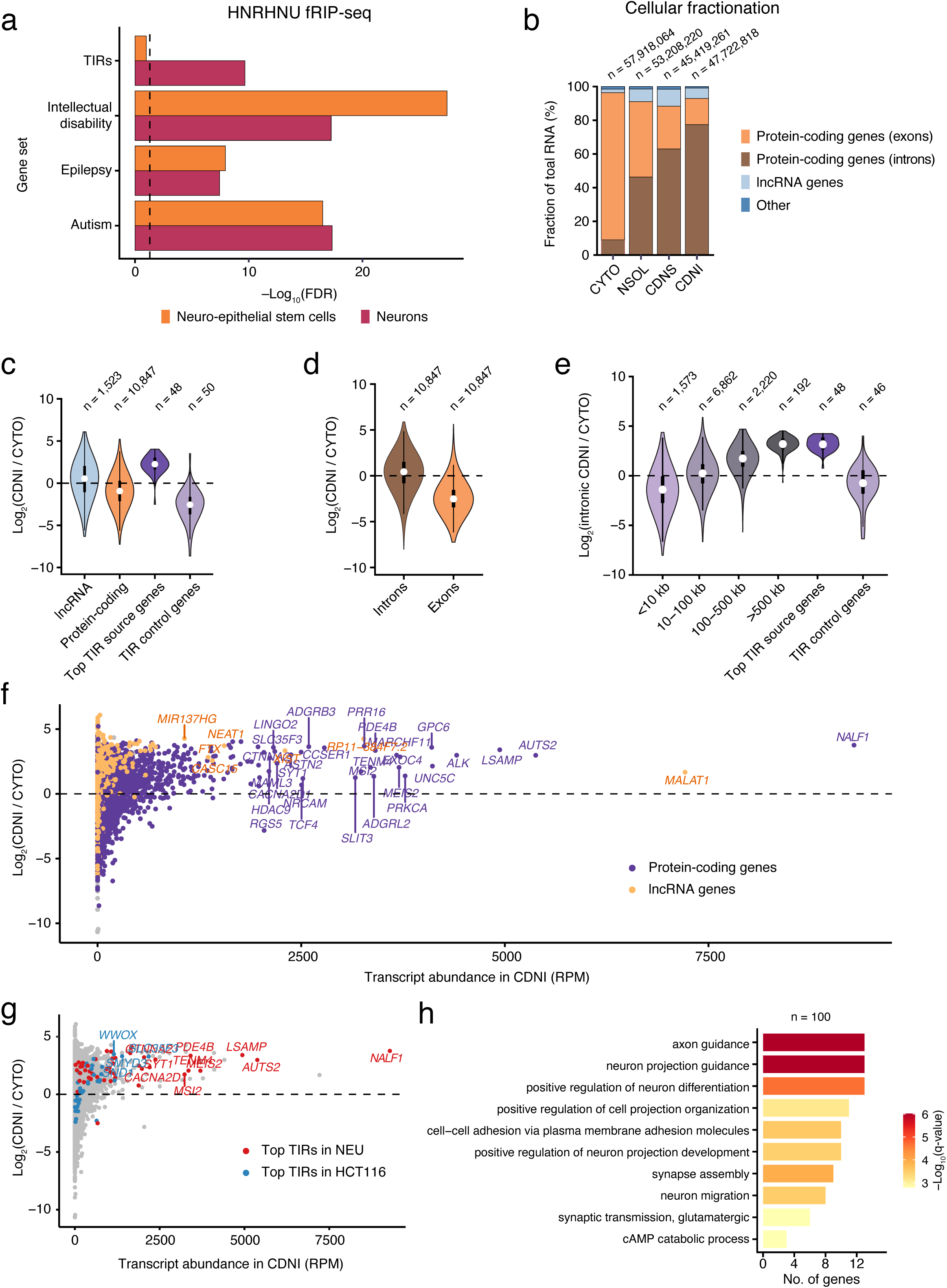
TIRs are enriched in the nuclear matrix. (**a**) Enrichment of the top-55 TIR source genes identified by RADICL-seq in NEU and of genes associated to autism, intellectual disability, and epilepsy risk among RNAs associated with the nuclear matrix protein HNRNPU, as detected by RNA-seq following formaldehyde crosslinking followed by ribonucleoprotein immunoprecipitation (fRIP-seq) against HNRNPU applied to neuro-epithelial stem cells (NES) and neurons (NEU) differentiated from them, relative to random expectation. The dashed vertical line indicates the significance threshold of false-discovery rate (FDR) lower than 0.05 (−Log10(FDR) > 1.3) based on hypergeometric testing. (**b**) Percentages of Total RNA-seq reads mapping to exonic or intronic regions of protein-coding genes, lncRNA genes, or other annotated genes, in four subcellular RNA fractions obtained from SH-SY5Y cells. CYTO, cytosolic fraction. NSOL, nuclear soluble fraction. CDNS, chromatin-depleted nuclear soluble fraction. CDNI, chromatin-depleted nuclear insoluble fraction. *n*, number of reads. (**c**) Distributions of the gene enrichment in the CDNI fraction relative to the CYTO fraction, quantified as the fold change in RNA counts between the CDNI and the CYTO fraction, for the indicated gene sets, in SH-SY5Y cells. *n*, number of genes in each set. RNA reads were quantified at the gene level. (**d**) As in (c) summing all the RNA reads mapping to intronic or exonic regions of protein-coding genes, in SH-SY5Y cells. (**e**) As in (c) summing all the RNA reads mapping to intronic regions for gene sets of different length in addition to the top TIR source and control genes, in SH-SY5Y cells. (**f**) Relationship between RNA abundance and gene enrichment in the CDNI fraction relative to the CYTO fraction, in SH-SY5Y cells. Each dot represents one gene. The dashed horizontal line represents Log_2_(CDNI/CYTO) = 0. RNA reads were quantified at the gene level. RPM, reads per million. (**g**) As in (f) with the top TIR genes identified in NEU and in HCT116 cells highlighted. (**h**) Enriched Gene Ontology (GO) biological process terms based on the most abundant genes (*n*) in the CDNI fraction relative to the CYTO fraction, in SH-SY5Y cells. In (c-e), violins extend from minimum to maximum, boxplots extend from the 25^th^ to the 75^th^ percentile, white dots represent the median, whiskers extend from –1.5×IQR to +1.5×IQR from the closest quartile. IQR, inter-quartile range. A link to the Source Data and code to regenerate the plots displayed in this figure is provided in the Data Availability and Code Availability statements.

We then tested the hypothesis that TIRs might be enriched in the chromatin-depleted nuclear insoluble fraction. To this aim, we leveraged a previously described cellular fractionation protocol^26^, which allows biochemical separation of four different cellular fractions: cytosolic (CYTO), nuclear soluble (NSOL), chromatin-depleted nuclear soluble (CDNS), and chromatin-depleted nuclear insoluble (CDNI), followed by total RNA-seq on each fraction (**Experimental Methods**, section 11). To account for the large input material (>20×10^6^ cells) required for the assay, we used the SH-SY5Y neuroblastoma cell line as a proxy of neuronal lineage. As a control, we also profiled the human colorectal cancer cell line, HCT116, for which RADICL-seq data are available in the FANTOM6 consortium (**Supplementary Table 1**). In both cell lines, we detected a marked enrichment of intronic RNAs derived from pc genes, in the CDNS and CDNI fractions compared to the CYTO and NSOL fractions, with ∼80% and ∼60% of all the RNAs in the CDNI fraction being intronic RNA in SH-SY5Y and HCT116, respectively (**Fig. 4b, Supplementary Fig. 23a, and Computational Methods**, section 11). By examining different types of RNA in the CDNI fraction, we found a weak enrichment of lncRNAs, depletion of transcripts encoding for pc genes in general, and, notably, a strong enrichment of RNAs originating from the same top-55 TIR source genes identified by RADICL-seq in NEU cells (**Fig. 4c and Supplementary Fig. 23b**). By further discriminating between exons and introns, we found that intronic RNAs derived from pc gene genes were enriched in the CDNI fraction, while exonic RNAs were depleted (**Fig. 4d and Supplementary Fig. 23c-e**).

The enrichment of intronic RNAs in the CDNI fraction was length-dependent at both the gene and intron level: in the first case, genes longer than 500 kb showed the highest enrichment in the CDNI fraction (**Fig. 4e and Supplementary Fig. 23f**), consistent with the pattern observed for TIRs. At the intron level, the analysis of individual introns of pc genes revealed a steep increase of the CDNI enrichment for short introns (<25 kb), followed by saturation for longer introns (**Supplementary Fig. 23g-j**), indicating a length-dependent but capped enrichment of intronic RNAs in the CDNI fraction. Surprisingly, despite *MALAT1* (the top trans-contacting RNA in NEU cells based on RADICL-seq) was expressed in SH-SY5Y cells at higher levels than *NALF1*, the latter was more enriched in the CDNI fraction (**Fig. 4f and Supplementary Fig. 23k**). Notably, among the top-55 TIRs identified in NEU, *AUTS2* and *LSAMP* were two of the three most abundant pc-gene RNAs detected in the CDNI fraction of SH-SY5Y cells (**Fig. 4f and Supplementary Fig. 23k**). Conversely, the HCT116 CDNI fraction was enriched in the TIRs identified by RADICL-seq in HCT116 cells but not in those identified in NEU cells, including *NALF1*, *AUTS2* and *LSAMP* (**Fig. 4g and Supplementary Fig. 23l, m**), consistent with the fact that these genes are specific for the neuronal lineage. Of note, the TIRs identified by RADICL-seq in HCT116 cells engaged in less widespread trans contacts compared to those identified in NEU cells (**Supplementary Fig. 23n**). The abundance of pc-gene RNAs in the HCT116 CDNI fraction was moderately correlated with the extent of genome-wide trans contacts formed by those RNAs, as detected by RADICL-seq (**Supplementary Fig. 23o**).

Next, we performed a GO analysis on the genes whose transcripts were enriched in the CDNI fraction of SH-SY5Y and HCT116 cells. In the former, most of the GO terms were related to neuronal processes/functions, whereas in the case of HCT116 cells most of the GO terms were linked to epithelial cells (**Fig. 4h and Supplementary Fig. 24a**), reflecting the different tissue of origin of these cell lines. In contrast, the most abundant RNAs detected in the CYTO fraction were shared between HCT116 and SH-S5Y5 cells and were associated to GO terms related to housekeeping functions (**Supplementary Fig. 24b, c**). Altogether, these results indicate that the TIRs detected by RADICL-seq are enriched in the chromatin-depleted insoluble nuclear fraction, further suggesting that TIRs might be constituents of the nuclear matrix/scaffold.

### TIRs contact specific genomic regions enriched in short neuronally expressed genes

Having thoroughly characterized the source genes of TIRs, we turned our attention to the DNA loci contacted by TIRs (**Computational Methods**, section 3). The genome-wide inter-chromosomal contact patterns of the top-10 TIRs identified by RADICL-seq in NEU appeared overall similar, even though not identical (PCC: 0.54–0.89 at 1 Mb; 0.29–0.75 at 100 kb), showing many broad trans contact peaks shared between different TIRs (**Fig. 5a and Supplementary Fig. 25a-c**). This indicates that different TIRs contact a common set of distant genomic regions. Accordingly, we found that the total number of trans contacts per genomic bin, formed collectively by all the TIRs identified by RADICL-seq, strongly correlated (PCC: 0.93 at 100 kb) with the number of distinct TIRs (produced by distinct source genes) contacting each bin (**Supplementary Fig. 25a**).

**Figure 5.**
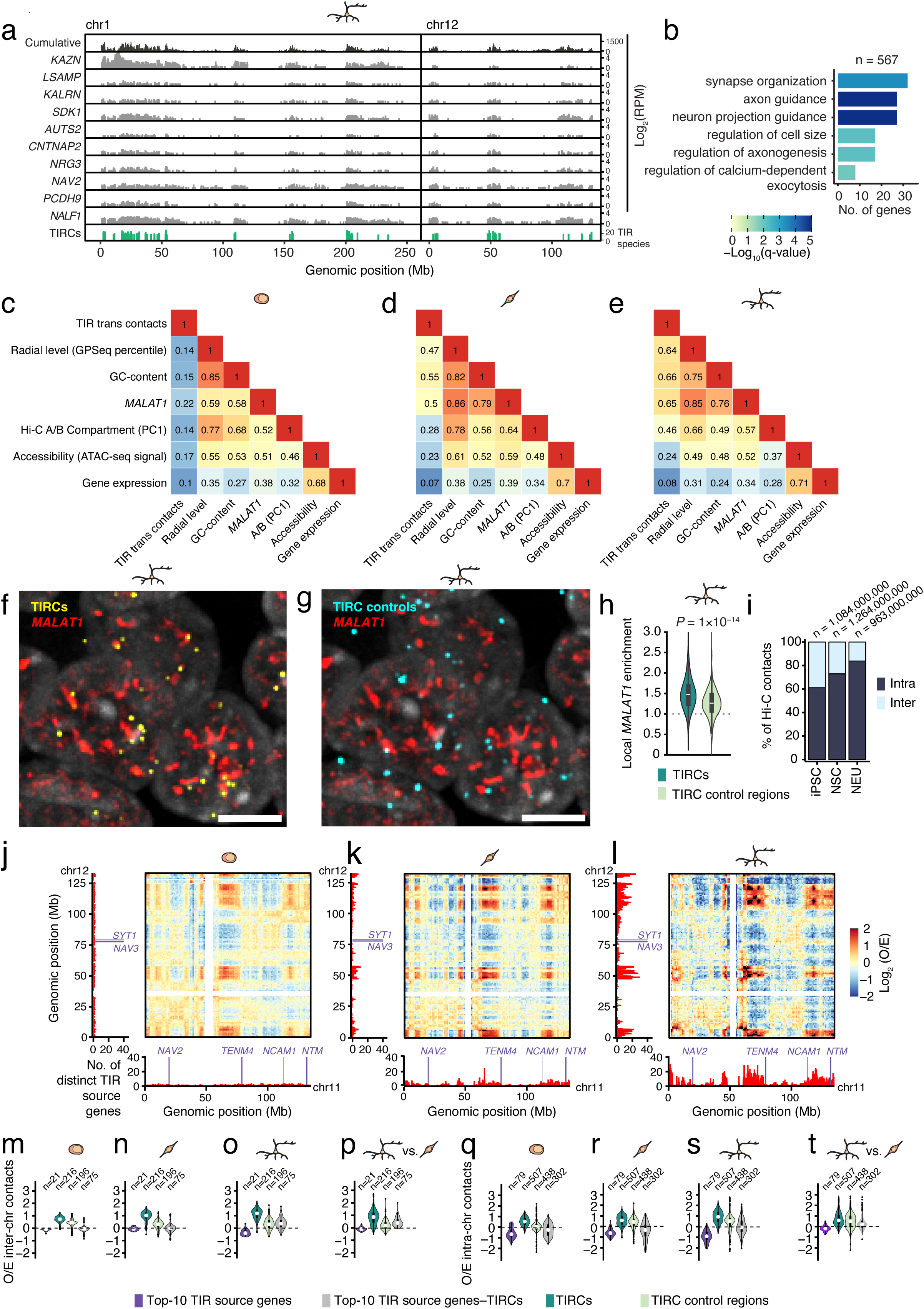
TIRCs are in proximity to speckles and form high-connectivity hubs. (**a**) Top track (dark gray): cumulative trans contact counts (100 kb resolution) along chromosome (chr) 1 and 12 for the top-10 TIR source genes identified in NEU. Light-gray tracks: trans contacts of each of the indicated top-10 TIR source genes. Bottom track (green): TIR contact regions (TIRCs) corresponding to 100 kb bins contacted by at least 16 distinct TIRs in NEU. RPM, reads per million. Only *CHiCANE* significant RADICL RNA-DNA pairs were considered (see **Computational Methods**, section 3). (**b**) Enriched gene ontology (GO) terms in biological processes based on 567 expressed genes (*n*) (among 916 protein-coding genes) within the 557 TIRCs identified in NEU. (**c-e**) Pairwise correlation (Spearman’s correlation coefficient) between the indicated genomic features (100 kb resolution) in iPSC (c), NSC (d), and NEU (e). TIR trans contacts: number of distinct genes producing TIRs contacting each genomic bin. Radial level: average GPSeq percentile in each genomic bin. GC-content: percentage of G and C in each genomic bin. *MALAT1*: raw count of contacts between *MALAT1* lncRNA and each genomic bin. PC1: Hi-C matrix principal component 1. Accessibility, mean height of pseudo-bulk ATAC-seq peaks per genomic bin (see Methods). (**f**) Maximum intensity projection of a z-stack widefield microscopy image exemplifying the spatial proximity between *MALAT1* lncRNA (marking speckles) and selected TIRCs, in one representative NEU cell nucleus, as detected by RNA and DNA FISH, respectively. Gray, DNA stained by Hoechst 33342. Scale bar, 5 μm. (**g**) As in (f) for selected TIRC control genes. (**h**) Quantification of the spatial proximity between *MALAT1* lncRNA and selected TIRCs or TIRC control regions, as exemplified in (f, g). (**i**) Percentage of intra– and inter-chromosomal DNA-DNA contacts identified by Hi-C in iPSC, NSC, and NEU. *n*, total number of Hi-C contacts. (**j-l**) Observed-over-expected (O/E) ratio of DNA-DNA inter-chromosomal contacts (1 Mb resolution) between chr11 and 12, detected by Hi-C. The bar plots on the left and bottom of each Hi-C matrix show the total number of TIRs (from distinct source genes) that engage in trans contacts with each 100 kb genomic bin, for chr12 and 11, respectively. The TIR source genes from the top-55 TIRs list that are located on chr11 or 12 are shown in purple. (**m**) Log2 of the O/E ratio of DNA-DNA inter-chromosomal contacts detected by Hi-C (1 Mb resolution) in iPSC, within each of the groups of genomic regions shown. *n*, number of genomic regions. (**n**) As in (m) for NSC. (**o**) As in (n) for NEU. (**p**) As in (m-o) but showing the difference between NEU and iPSC of the Log2 of the O/E ratio of DNA-DNA inter-chromosomal contacts. (**q-t**) As in (m-p) for intra-chromosomal contacts in trans (DNA-DNA distance > 5 Mb; 100 kb resolution). In all the violin plots in the figure, violins extend from minimum to maximum, boxplots extend from the 25^th^ to the 75^th^ percentile, white dots represent the median, whiskers extend from – 1.5×IQR to +1.5×IQR from the closest quartile. IQR, inter-quartile range. A link to the Source Data and code to regenerate the plots displayed in this figure is provided in the Data Availability and Code Availability statements.

We then focused on regions contacted by many different TIRs and identified 557 bins (100 kb) contacted by at least 16 different TIRs in NEU (**Fig. 5a**, bottom track **and Supplementary Fig. 25a**). We hereafter refer to these regions as TIR-contacted regions or TIRCs. To validate the trans contacts identified by RADICL-seq, we performed combined DNA&RNA FISH in NEU cells, using RNA FISH probes targeting the top-10 TIRs and DNA FISH probes targeting the top-10 shared TIRCs alongside an equal number of control DNA loci lacking TIR-mediated contacts (**Supplementary Table 3 and Experimental Methods**, section 10). Consistent with the RADICL-seq results, TIRs were significantly more co-localized with TIRCs than with control genomic regions defined as 100 kb genomic bins selected based on the absence of trans RNA contacts and on radial positioning similar to that of TIRCs in iPSC (**Supplementary Fig. 25d, e**). This indicates that these trans contacts reflect a genuine biological phenomenon rather than representing noise.

Next, we performed GO analysis of the expressed genes localized within TIRCs. In NEU, these genes were enriched in GO terms related to neuronal functions (**Fig. 5b**). Notably, the enrichment was higher for TIRCs than for *MALAT1* contact regions in NEU or for genes within the A compartment (**Supplementary Fig. 26a, b**). Among genes related to neuronal functions, TIRCs contained genes specifically upregulated in NEU, similarly to the TIR source genes (**Supplementary Fig. 26c**). In comparison, housekeeping genes within TIRCs were not upregulated during neuronal differentiation (**Supplementary Fig. 26c**). However, we found that, in contrast to the TIR source genes, neuronally expressed TIRC genes are exceptionally short and have a significantly higher GC-content (**Supplementary Fig. 26d, e**). Altogether, these results indicate that, in neurons, TIRs tend to engage with multiple shared regions along the genome, which are enriched in short, active genes associated with neuronal functions.

### TIRCs show increased inter-chromosomal connectivity in neurons

We then analyzed the identified TIRCs by intersecting them with various genomic annotations (**Fig. 5c-e and Supplementary Fig. 27a**). In NEU, the genome-wide profiles of the trans contacts formed by the top-55 TIRs identified in these cells and the genome-wide profile of *MALAT1* trans contacts were overall similar (Spearman’s correlation coefficient, SCC: 0.65), although the latter displayed less distinct peaks (**Supplementary Fig. 27b**). In NSC and even more in NEU, but not in iPSC, NEU-specific TIRCs had a higher speckle proximity score compared to the TIR source genes and TIRC control regions, and TIRCs were enriched in the A compartment (**Supplementary Fig. 27a, c-e**). To further investigate the relationship between TIRCs and speckles, we performed combined DNA&RNA FISH in NEU using an RNA FISH probe targeting *MALAT1* lncRNA (as a proxy for speckles) and DNA FISH probes targeting 16 selected TIRCs (**Supplementary Table 3 and Experimental Methods**, section 10). We then measured the extent of co-localization between DNA and RNA FISH dots and found that TIRCs were significantly closer to speckles in comparison to the control regions (**Fig. 5f-h and Computational Methods**, section 8). Of note, DNA accessibility, as measured by scATAC-seq, was slightly lower inside TIRC genes in comparison to TIRC control regions (**Supplementary Fig. 27a and Computational Methods**, section 12), suggesting that TIRs-TIRCs contacts are not driven by a particularly elevated accessibility of TIRCs in neurons.

The 3D contacts of TIRCs, as assessed by Hi-C, also changed dramatically during cell differentiation towards the neuronal lineage. The ratio between intra– and inter-chromosomal contacts was substantially higher in NEU compared to NSC and iPSC (**Fig. 5i**); however, despite a global decrease of inter-chromosomal contacts in NEU, inter-chromosomal contacts between different NEU-specific TIRCs progressively intensified during differentiation (**Fig. 5j-l**). Notably, we detected enhanced interactions between TIRCs located on different chromosomes (**Fig. 5m-p**) as well as between TIRCs residing on the same chromosome but separated by more than 5 Mb (**Fig. 5q-t**). We also observed a moderate increase in interactions between TIRC and TIR source loci (**Fig. 5m-t**). In contrast, interactions among genomic regions harboring the top-10 TIR source genes were reduced (**Fig. 5m-t**). Overall, these results indicate that, in neuronal cells, TIRCs form a high-connectivity chromatin neighborhood, which is in close spatial proximity with speckles.

### TIR source genes move closer to TIRCs during NEU differentiation

Next, we sought to understand the spatial context of TIRs-TIRCs interactions. To this end, we leveraged our GPSeq method^37^ for mapping genome radiality and our latest GPSeq protocol^52^ to assess if and how the genome is radially reorganized during neuronal differentiation (**Experimental Methods**, section 5 and **Computational Methods**, section 13). On a global scale, we identified multiple regions that clearly moved towards the nucleus center during the transition from iPSC to NEU, as well as regions that moved towards the nuclear periphery (**Fig. 6a and Supplementary Fig. 28a**). GO term analysis of the inward-moving regions revealed a significant enrichment in genes associated with neuronal functions, whereas the regions moving outwards were enriched in genes associated with skin development (**Supplementary Fig. 28b, c**), possibly reflecting the fact that the iPSC used in this study were derived from skin fibroblasts. These radial changes were linked to gene expression changes: genes within inward-moving regions were upregulated in NEU compared to iPSC, whereas genes in regions moving towards the nuclear periphery were downregulated (**Supplementary Fig. 28d**).

**Figure 6.**
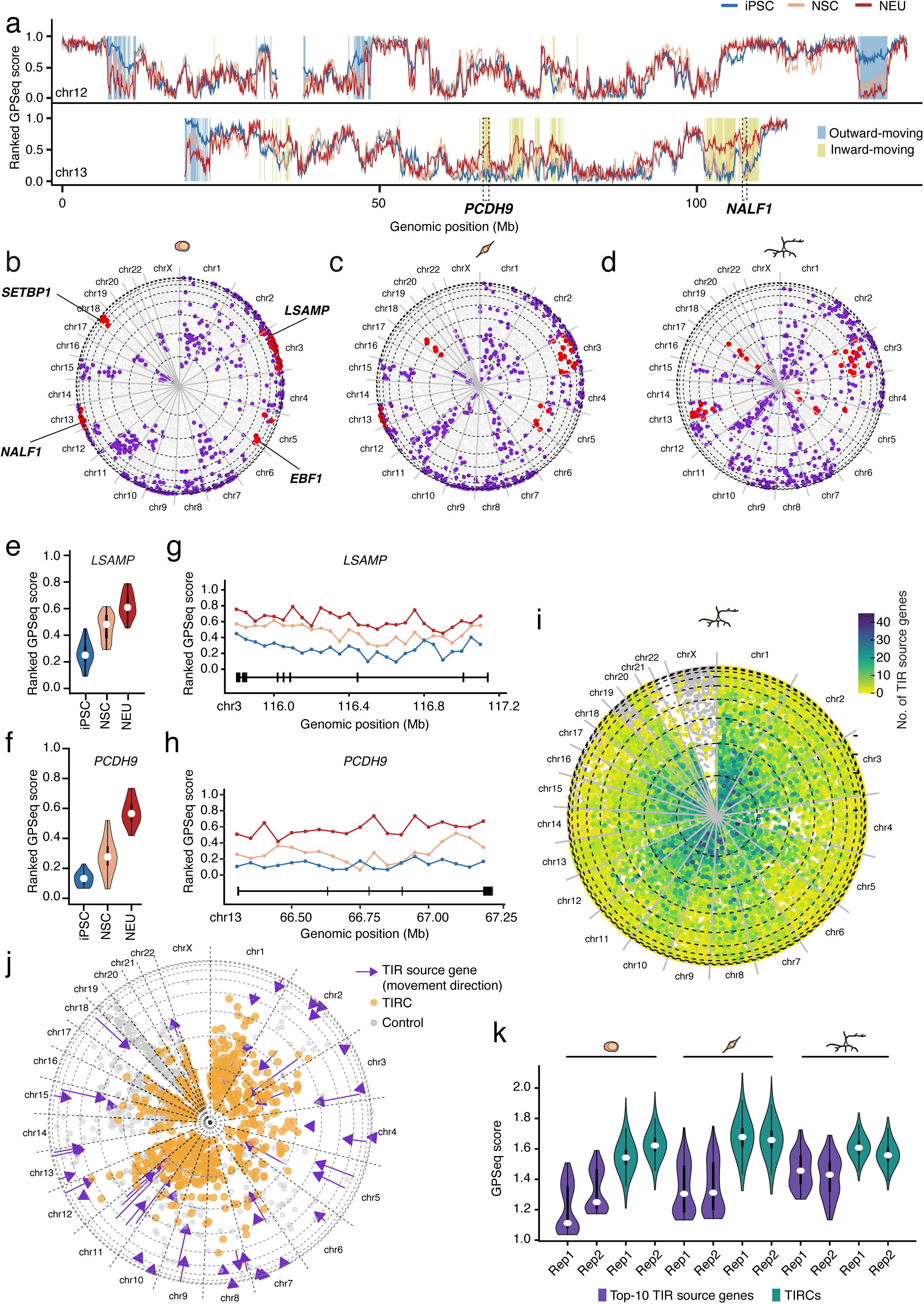
TIR source genes relocate radially during neuronal differentiation. (**a**) GPSeq percentiles (100 kb resolution) along chromosome (chr) 12 and 13 in iPSC, NSC, and NEU. Dashed boxes indicate the gene bodies of two of the top-55 TIR source genes identified in NEU. The regions on chr12 and 13 included in the top-1000 regions moving outward or in top-1000 regions moving inward from iPSC to NEU are shown. (**b-d**) ‘Pizza plots’ showing the radial placement (100 kb bins) of the top-55 TIR source genes (in purple and red) in iPSC (b), NSC (c) and NEU (d), as measured by GPSeq. Four representative TIR source genes are indicated. Purple dots represent 100 kb genomic bins overlapping with the top-55 TIR source genes detected in NEU. Gray dots in the background represent the remaining 100 kb genomic bins. The outermost and innermost dashed circle of each pizza plot represent the center and periphery of the cell nucleus, respectively. Dashed circles from center to periphery represent GPSeq score deciles. Radii separate individual chromosomes. Each dot represents a 100 kb bin. Dots occupying the same radial position within the same chromosomal sector are jittered for visualization. (**e, f**) Distribution of the GPSeq score percentile of 50 kb genomic bins overlapping the indicated two of the top-10 TIR source genes identified in NEU, in iPSC, NSC, and NEU. (**g, h**) As in (e, f) showing the distribution of GPSeq score percentiles along the gene body of the indicated TIR source genes. The black line vertical bars below the GPSeq score percentiles schematically represent the exons of the corresponding gene, whereas the horizontal black line crossing the vertical bars represents the gene body. (**i**) ‘Pizza plots’ showing the number of TIRs (expressed as number of distinct source genes) contacting 100 kb genomic bins radially placed based on GPSeq data from NEU. (**j**) ‘Pizza plots’ showing the radial re-positioning of the 557 TIRCs identified in NEU (orange dots), of the 557 TIRC control regions (grey dots), and of the top-55 TIR source genes identified in NEU (purple arrowheads), as assessed by GPSeq. The purple arrows indicate the magnitude and direction of the change in radial position of the TIR source genes from iPSC to NEU. (**k**) Distributions of the GPSeq scores of 100 kb genomic bins overlapping with the top-10 TIR source genes and TIRCs identified in NEU, in iPSC, NSC, and NEU. A link to the Source Data and code to regenerate the plots displayed in this figure is provided in the Data Availability and Code Availability statements.

We then set out to analyze the radial location and the possible repositioning of the TIR source genes within the nucleus, during neuronal differentiation. We found that the TIR source genes identified by RADICL-seq in NEU were located more peripherally than the genes producing RNAs involved only in cis RNA-DNA contacts (**Supplementary Fig. 29a**). As cells differentiated to neurons, the radial position of the top-10 TIR source genes identified in NEU changed significantly, with many genes showing a gradual relocation, moving towards the nuclear center during the iPSC-to-NSC transition and, further, during the NSC-to-NEU transition (**Fig. 6b-f and Supplementary Fig. 29b-i**). Notably, the radial position change differed along the long gene body of each of the top-10 TIR source genes, with some portions of the gene body radially relocating more compared to other parts of the same gene (**Fig. 6g, h and Supplementary Fig. 29j-q**). As a result of this process, the spread of the top-55 TIR genes across multiple radial layers was substantially higher in NEU in comparison to iPSC or NSC (**Supplementary Fig. 29r**).

Lastly, we assessed the radial arrangement of TIRCs. We found that, while RADICL-seq contacts were mapped all along the nuclear center-periphery axis, TIRCs tended to occupy more central parts of the nucleus in NEU (**Fig. 6i and Supplementary Fig. 30a-c**). In fact, upon neuronal differentiation, the TIR source genes moved progressively closer to TIRCs—radially—with both gene groups ending up close to the nuclear center, in NEU (**Fig. 6j, k**). These results indicate that, upon cell differentiation to neurons, the genome undergoes substantial radial reorganization, with TIR source genes moving closer to the transcriptionally active center of the nucleus and becoming spatially close to TIRCs.

### Expression of TIR source genes and TIRC genes strongly co-varies between single cells

Having found that TIRCs are part of high-connectivity hubs and that TIR source gene loci move radially towards TIRCs during neuronal differentiation, we hypothesized that this spatial configuration might favor co-regulation of the expression of TIR source genes and of TIRC genes, therefore resulting in co-variation in their expression. To test this hypothesis, we leveraged a method for calculating gene expression co-variation from scRNA-seq data^53^ and applied it to publicly available scRNA-seq datasets from motor neurons derived from SOD1 A4V/+ iPSC^54^ (**Computational Methods**, section 14). Using this approach, we calculated a co-variation enrichment score (CES) for different pairs of genes: i) between genes inside TIRCs; ii) between different TIR source genes; and iii) between TIRC and TIR source genes. Intriguingly, for genes located in TIRCs, the CES score progressively increased with increasing numbers of shared TIRC-interacting with TIRs, reaching the highest values for genes inside TIRCs contacted by the same set of 7–8 different TIRs (**Fig. 7a**). TIR source and TIRC genes were associated with substantially higher CES values compared to genes inside TIRC control regions (**Fig. 7b**). Importantly, TIR-TIRC gene pairs also showed relatively high CES scores compared to the control gene set (**Fig. 7b**), indicating that their expression also co-varies. Altogether, these results suggest that the expression of TIR source genes and the expression of TIRC genes might be linked by a common regulatory mechanism, and we hypothesize that TIRs play a role in mediating this regulation.

**Figure 7.**
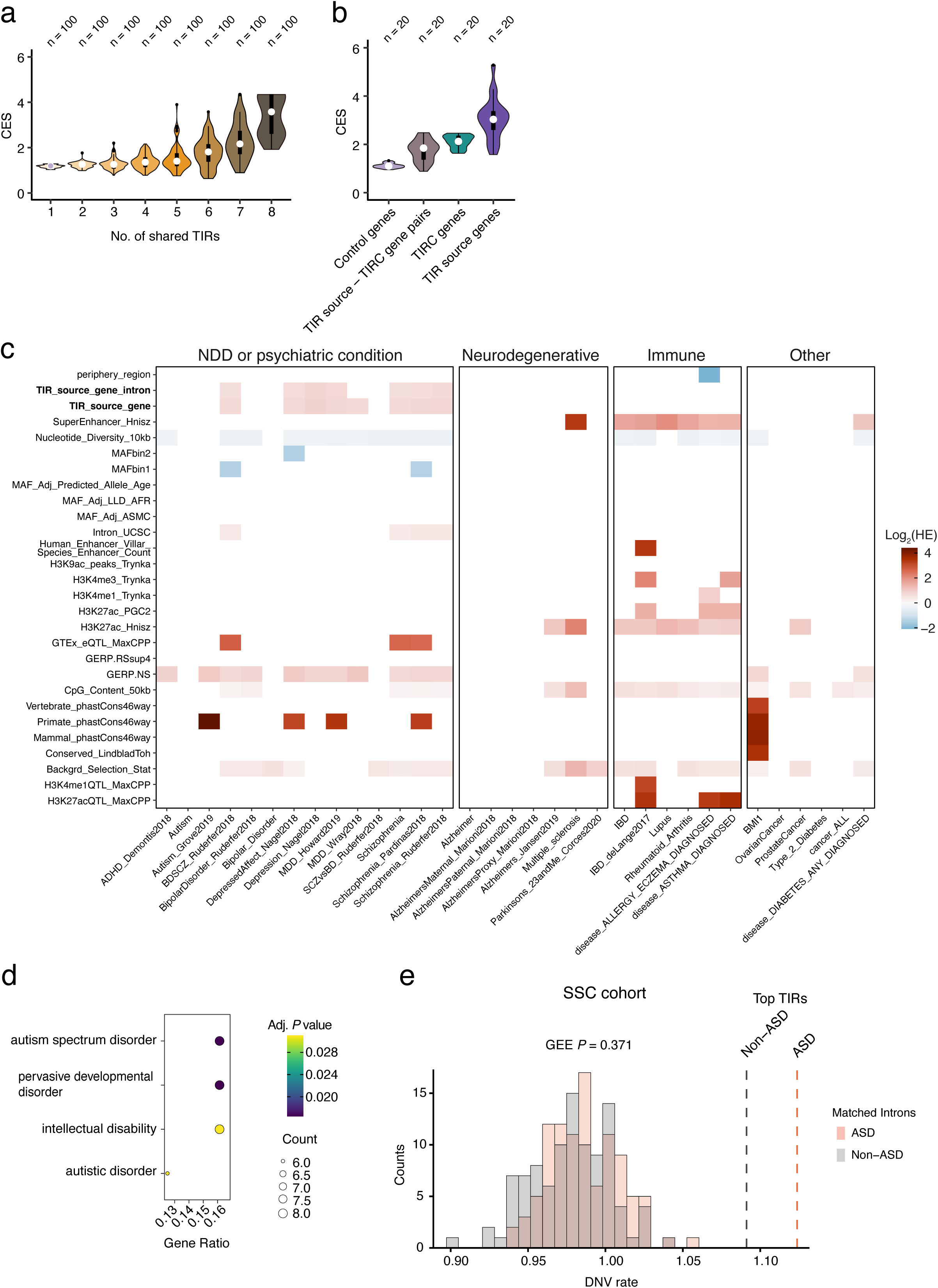
TIR source genes and TIRCs are associated with neurodevelopmental disorders. (**a**) Distribution of the gene covariation enrichment score (CES) for different sets of expressed genes located inside the genomic regions (100 kb resolution) co-contacted by TIRs produced from the indicated number of the top-10 TIR source genes identified in NEU. *n*, number of random samplings used to calculate the CES (see **Computational Methods**, section 14). (**b**) As in (a) for covariation within or between the indicated gene groups. *n*, number of random samplings used to calculate the CES. (**c**) Heritability enrichment (HE) of single-nucleotide polymorphisms (SNPs) identified across multiple genome-wide association studies (GWAS) grouped by trait analyzed (x-axis) within the annotation groups listed on the y-axis. Significant HE is shown (P < 0.00001). Traits are based on the S-LDSC results from 33 GWAS (see **Supplementary Table 4**). NDD, neurodevelopmental disorder. (d) Disease ontology enrichment analysis of the top-55 TIR source genes identified in NEU. (**e**) Counts of significance level (p-values) in 100 GEE models assessing the association between DNV count and intron set types (top-55 TIR introns versus matched introns from NSC-expressed genes). The dashed vertical lines represent DNV event rates in the introns of the top-55 TIR genes. GEE, Generalized Estimating Equation. In all the violin plots in the figure, violins extend from minimum to maximum, boxplots extend from the 25^th^ to the 75^th^ percentile, white dots represent the median, whiskers extend from –1.5×IQR to +1.5×IQR from the closest quartile. IQR, inter-quartile range. A link to the Source Data and code to regenerate the plots displayed in this figure is provided in the Data Availability and Code Availability statements.

### The introns of TIR source genes are enriched in neuropsychiatric disorder-associated loci

Lastly, we explored whether TIRs or TIRCs are somehow related to neurodevelopmental or neuropsychiatric conditions, given that many genes involved in these disorders include very long genes specifically expressed in neurons. To this end, we applied stratified linkage disequilibrium score regression (S-LDSC)^55^ to test whether the heritability of single-nucleotide polymorphisms (SNPs) associated with a given disorder is enriched in TIR source genes, TIRCs or in 96 annotated genomic regions from the so-called Baseline model^56^ (**Supplementary Table 4 and Computational Methods**, section 15). As control, we applied S-LDSC to partition the SNP heritability of 14 disease traits unrelated to neurodevelopmental and neuropsychiatric disorders, including 7 neurodegenerative disorders, 6 immune-related diagnoses and 6 other diagnoses (**Supplementary Table 4**). S-LDSC computes a heritability enrichment (HE) score, which indicates whether a genomic region of interest is associated with a greater proportion of the heritability of a given trait, as compared to the proportion of heritability expected based on the SNPs contained in the same region. We found that the introns of TIR source genes as well as regions with high neutral rate score based on Genomic Evolutionary Rate Profiling (GERP.NS)^57^ had the most significant and consistent HE scores for neuropsychiatric disorders, but not for control traits (**Supplementary Fig. 31**). Specifically, we observed a significant HE score (P < 0.00001) inside the introns of the TIR source genes for 7 out of 14 neurodevelopmental and neuropsychiatric conditions versus 0 out of 19 control traits (**Fig. 7c**). Moreover, disease ontology analysis of the top-55 TIRs identified by RADICL-seq in NEU revealed terms linked to autism and intellectual disability (**Fig. 7d**).

To further corroborate the link between TIRs and neurodevelopmental/neuropsychiatric disorders, we analyzed de novo variants (DNV) from two autism cohorts including neurotypical sibling controls (**Computational Methods**, section 16). We found that the introns of the top-55 TIR source genes were enriched in DNVs, in comparison to introns matched for GC-content and length (**Fig 7e, Supplementary 32a-c, and Supplementary Table 5**). In both cohorts, we observed a difference in the mean DNV rate between autism and non-autism, with the former having higher DNV rates in the top-55 TIR introns, although the differences did not reach statistical significance (**Fig 7e and Supplementary 32a-c**). Altogether, these results demonstrate that TIR source genes are enriched in genomic loci associated with neurodevelopmental and neuropsychiatric conditions and suggest that the exceptionally long introns of TIR genes harbor functional regulatory sequences that might be implicated in the pathogenesis of these disorders.

## Discussion

Our study provides a comprehensive portrait of the RNA-DNA contactome, 3D genome, and transcriptome—and their dynamic changes—during in vitro differentiation of human iPSC to neurons. Through integrative omic analyses, we have identified a novel class of chromatin-associated RNAs (caRNAs) that form widespread contacts with DNA, extending far from the locus where they are transcribed. Contrary to our expectation, the majority of trans-contacting caRNAs that we identified are not derived from lncRNAs, with the exception of *MALAT1*, a well-known constituent of nuclear speckles that forms widespread genomic contacts^45,58–60^. Instead, except for *MALAT1* lncRNA, most of the trans-contacting caRNAs are transcribed from the introns of protein-coding genes involved in cell type-specific functions; we therefore refer to this group of caRNAs as TIRs, for Trans-contacting Intronic RNAs. We detected TIRs across multiple cell types profiled by RADICL-seq in FANTOM6, including three cancer lines (MCF7, K562, HCT116). However, neuronal cells display by far the highest number and diversity of TIRs, which progressively increase during neuronal differentiation. This is possibly related to the observation that TIRs derive from very long genes (which typically carry long introns), which are among the most highly expressed genes in neurons^8^. Remarkably, more than a decade ago, it was shown that fetal human brain cells harbor high levels of intronic RNA derived from long genes associated with neuronal functions^61^. While this observation was linked to co-transcriptional and alternative splicing—both of which are common for long, synapse-associated genes^62^—the exact relevance of those intronic RNAs has remained elusive ever since.

Leveraging high-resolution RNA FISH, we discovered that, in neurons, TIRs form cloud-like agglomerates around their source loci, in addition to spreading across the nucleus. Despite the fact that we only visualized a small fragment of each intron transcribed from the top-10 TIR source genes detected in NEU, this was sufficient to uncover a meshwork extending through a large portion of the nucleus of these cells. Moreover, TIRs produced by different source genes were intermixed, forming a continuous network across the whole nucleus. We propose that this meshwork corresponds to the chromatin-depleted nuclear insoluble (CDNI) fraction that we have profiled using an orthogonal approach and that is mainly composed of intronic RNAs, in line with a previous report^26^. The fact that the CDNI fraction is enriched in intronic RNAs and depleted of exonic sequences strongly suggests that the intronic RNAs found in the CDNI fraction correspond to spliced-out RNA rather than being part of nascent RNA molecules.

At this point, we do not know whether TIRs contact DNA or other RNAs directly (via complementary interactions or through triplex formation) or indirectly (e.g., through RNA-binding proteins). However, irrespectively of whether TIRs engage in specific or unspecific contacts with DNA or other RNA types, we postulate that the sole mass of TIRs accumulating in the nucleus of neuronal cells might, on its own, influence nuclear physiology. Based on the above considerations, we name this previously uncharted nuclear compartment as ***introsomes*** and propose the model shown in **Figure 8** to explain their formation. Briefly, in cells with broadly open chromatin—as in embryonic stem cells^40^—very long genes carrying long introns are mostly silenced and typically located at the nuclear periphery; hence, introsomes do not form in those cells or, if they do, they contact chromatin in a seemingly random fashion (based on the absence of significant levels of genome-wide trans contacts in our iPSC RADICL-seq data). As an increasing number of long genes start being expressed as differentiation progresses, transcription leads to their decompaction, in turn causing their repositioning away from the nuclear periphery. When the introns of those long genes are then spliced, they accumulate in the nucleus instead of undergoing rapid degradation. The function of these introns seems intrinsically linked to their own turnover: they are produced, exert a structuring role in cis and in trans, and are then degraded, yet continuously replenished through ongoing transcription of the same long genes. This results in the emergence of a meshwork whose biophysical properties are dictated by the length and sequence composition (A/U-rich in neurons) of the intronic RNAs composing it, as well as by the type of RNA-binding proteins recruited to them, such as SAF-A, as we have shown here. In the absence of cellular division, the accumulation of TIRs continues, leading to the formation of a widespread network of intronic RNAs bridging all the DNA loci from which they are actively transcribed. This meshwork possibly creates a sponge-like environment that brings in proximity various protein complexes required for transcription, splicing, and/or repair. The TIR meshwork might correspond to the principal component of the interchromatin compartment (IC) previously proposed as part of the Active Nuclear Compartment (ANC) – Inactive Nuclear Compartment (INC) Model^63^, in line with a recent report^64^. In fact, the accumulation of large intronic RNA masses around multiple TIR source loci distributed across the nucleus might underlie the formation of the IC. We propose that, in neuronal cells, this scaffold reshapes the global 3D genome structure, leading to more defined inter-chromosomal connectomes by connecting TIR source loci with regions (TIRCs) that are preferentially located at the outer surface of chromosomal territories. The ‘consolidation’ of the 3D genome structure in neuronal cells might be critical for (co-)regulation of genes residing in TIRCs, many of which are short, neuron-specific genes activated during differentiation, whose expression co-varies both among themselves and with that of TIR source genes, at the single cell level. 3D genome consolidation might be additionally favored by radial reorganization of chromatin as stem cells differentiate towards neurons, as revealed by our GPSeq data.

**Figure 8.**
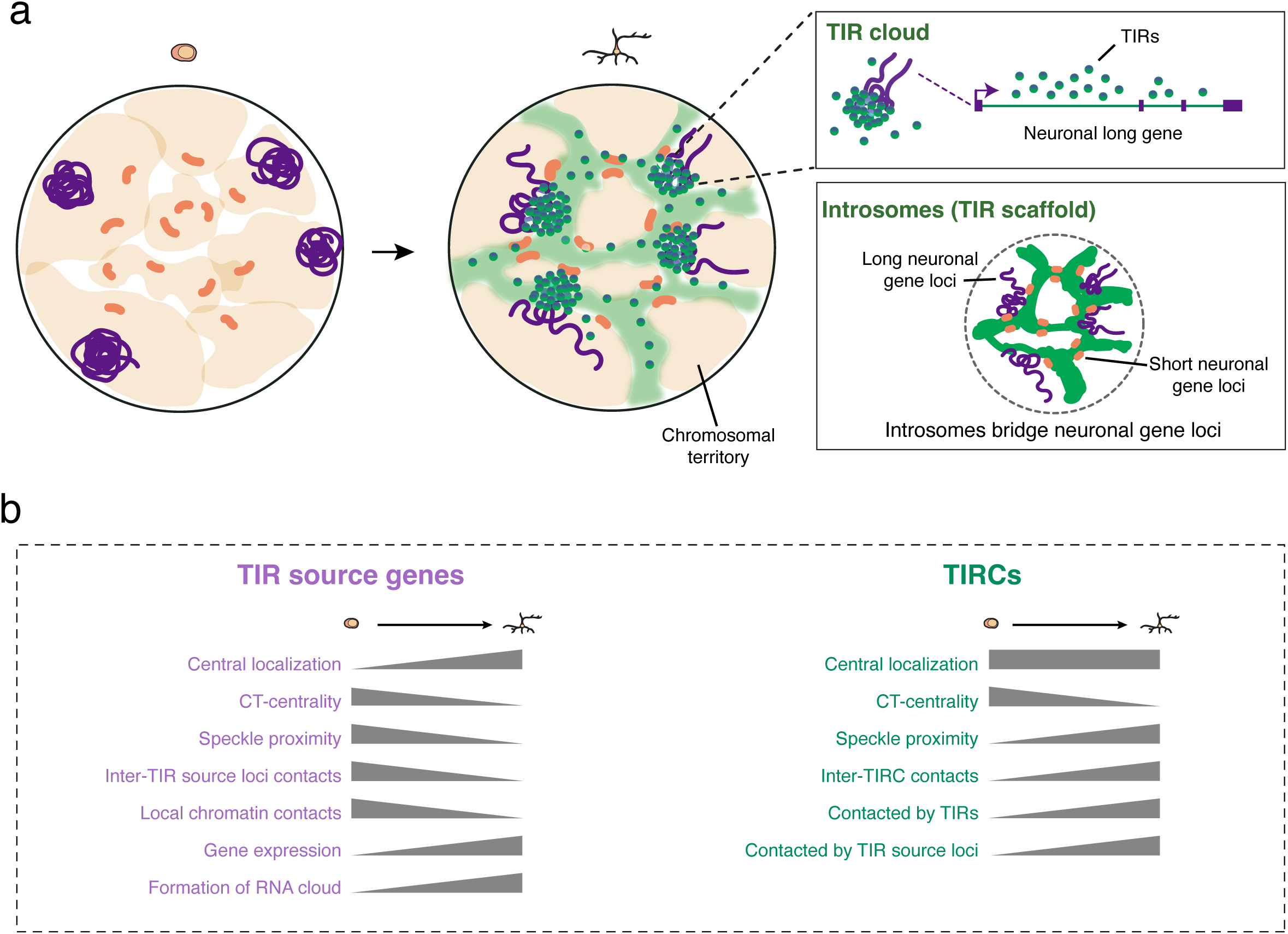
Proposed model of introsomes formation and their main features. (**a**) In iPSC, very long genes containing long introns encoding for trans-contacting intronic RNAs (TIRs) are transcriptionally silent and are typically positioned near the nuclear periphery. Upon differentiation to NEU cells, activation of TIR source genes leads to transcription-associated chromatin decompaction, resulting in the repositioning of these genes away from the nuclear periphery as well as from the interior towards the surface of chromosomal territories (CT). In parallel, expression of TIR source genes leads to the accumulation of TIR clouds around the source locus, with additional sparse intronic RNAs scattered throughout the nucleus. Collectively, TIRs originating from multiple source loci assemble into a widespread RNA meshwork—named **introsomes**—that fills in the inter-chromosomal space. Introsomes create a sponge-like nuclear environment that recruits protein complexes, bridging distant TIR source genes together with TIR-contacted regions (TIRCs) that are enriched in short genes specifically expressed in neurons. This process establishes a neuron-specific nuclear architecture that enables coordinated co-regulation of genes essential for neuronal functions and for the maintenance of neuronal cell identity. (**b**) Distinctive features of TIR source genes and TIRCs, and schematic representation of their changes during neurodifferentiation.

Based on recent studies suggesting that nascent RNAs form microgel-like liquid-liquid phase separated entities in the nucleus^30,31^, we speculate that introsomes follow the same principle. In fact, the biochemical properties of long intronic RNAs make them ideal candidates for nucleating phase separated condensates, especially considering a recent study that identified the length of RNAs as well as their sequence composition as key factors predisposing them to form condensates^65^. Of note, the A/U-rich RNA cluster identified in that study^65^ increased in strength during cellular differentiation.

Although in the present study we have profiled a limited number of cell lines, our results suggest the possibility that different cell types harbor different introsome repertoires, as the top TIR source genes differ across cell lines. Based on our data, the RNA composition of introsomes is largely different from the composition of the soluble RNA fraction that is commonly profiled by RNA-seq and strongly depends on the type of (long) introns that are expressed in each cell. We propose that introsomes represent a specialized, differentiation-dependent component of the nuclear matrix, whose composition—particularly the length distribution and A/U richness of its intronic RNA constituents—shapes the cell-type specificity of the nuclear matrix. In turn, the composition and abundance of introsomes might significantly affect the morphology and biophysical properties of the nucleus. We propose that introsomes might not be required in pluripotent cells, which lack cell type-specific transcriptional programs, but become prominent in terminally differentiated cells where they help establish and maintain the stable nuclear architecture required for sustained lineage-specific gene expression. We further speculate that each cell type is characterized by a distinctive intronic RNA length-to-species-diversity ratio—with neurons harboring introsomes dominated by a small number of ultra-long intronic RNAs, while other cell types, such as HCT116, harbor introsomes composed of a larger number of long but not ultra-long intronic RNA species. Further studies will be needed to address these exciting questions.

Our observations that, in neuronal cells, TIRs originate from ultra-long, neuronally expressed genes located at the nuclear periphery are possibly related to the finding of mega-enhancer bodies in cerebellum neurons during mouse development, as reported in a recent pre-print^66^. In this study, long neuronal genes were found to localize to a unique nuclear sub-compartment located at the nuclear periphery and relatively far from speckles, like we observe here for the TIR source genes. The authors speculated that the peripheral localization of mega-enhancer bodies might be driven by physical constraints linked to the large size of the genes contacted by these enhancers, for which the nuclear center might not provide sufficient space, especially considering that many long genes are expressed simultaneously during neuronal differentiation, in line with previously reported transcription loops formed by highly-expressed long genes^48^. Notably, the study describing mega-enhancer bodies also reported the identification of two different A sub-compartments: one carrying long neuronal genes forming mega-enhancer bodies, and the other comprising gene-dense genomic regions localized close to speckles in the nuclear center. We speculate that these two sub-compartments correspond, respectively, to the TIR source genes and TIRCs that we have identified in this study. We propose that TIRs somehow mediate the communication between these two sub-compartments since they both harbor genes that become specifically activated during neuronal differentiation. Furthermore, it has been reported that exceptionally long genes localize close to Polycomb bodies, away from nuclear speckles^67^. The authors of this study proposed that the separation of long genes from speckles might protect them from premature internal splicing before the long introns of these genes are fully transcribed, further supporting our observations. Lastly, it has also been reported that transcriptionally active long neuronal genes are associated with RNA polymerase II (RNAPII)-enriched regions that form a distinct nuclear compartment located far from nuclear speckles^68^. Future studies will need to assess whether TIRs contribute to the formation of mega-enhancer bodies and assess how TIR source genes are spatially related to Polycomb bodies and RNAPII compartments in different cell types.

Functionally, our study provides several lines of evidence supporting an important role of TIRs in neurobiology. First, the expression of TIR source and TIRC genes significantly co-varies, especially among TIRC genes contacted by common sets of TIRs, as revealed by our gene expression co-variation analysis. This suggests that TIRs somehow co-regulate the expression of multiple TIR source genes and, possibly, TIRC genes. We speculate that, in neuronal cells, introsomes provide a scaffold that spatially organizes TIRC genes, leading to the formation of highly connected TIRC networks, which in turn are essential for proper gene (co)-regulation inside these genomic regions.

Remarkably, we found that TIR source genes are significantly enriched in SNPs associated with several neurodevelopmental and neuropsychiatric conditions, including bipolar disorder and schizophrenia. Indeed, our disease ontology analysis revealed that the source genes of the top TIRs identified in neurons are enriched in terms linked to autism and intellectual disability and are typically ultra-long genes similarly to those associated with Rett syndrome, a rare neurodevelopmental disorder associated with mutations in the *MECP2* gene^69^. Moreover, the previously reported significant enrichment of intronic RNA in the human fetal brain in comparison to the adult brain^61^ further implicates TIRs in neurodevelopment. We also observed enrichment of de novo variants within TIR introns when compared with non-TIR introns matched for GC-content and length. The de novo mutation rate within the intronic sequence of TIR source genes was higher in autism but not significant in comparison to unaffected siblings, suggesting that TIR source genes may reside in genomic contexts with intrinsically higher mutational input or detectability. However, the slightly greater enrichment observed in autism, together with increased evolutionary constraint and enrichment for GWAS heritability of neurodevelopmental/neuropsychiatric traits, strongly suggests that these introns harbor functionally relevant regulatory elements. The convergence of rare de novo variation and common-variant heritability within the same noncoding regions is consistent with a recent study showing that rare and common variants perturb shared regulatory architecture underlying neurodevelopmental and psychiatric liability^70^. We also found that TIRs interact with one of the components of the nuclear matrix—HNRNPU—whose mutations are linked to severe neurodevelopmental disorders in humans, including autism^34^. Although, at present, we do not know whether TIRs are expressed at normal levels and form clouds in neuronal cells derived from patients with autism or other neurodevelopmental/neuropsychiatric conditions, we speculate that certain condition-associated variants might reduce the levels of or change the biochemical properties of the intronic sequences harbored in TIRs, or affect their interaction with RBPs, in turn altering their ability to cross-regulate the expression of many genes involved in neurodevelopment. Future studies integrating splice prediction, chromatin annotation, and experimental perturbations will be necessary to determine whether individual variants within TIR source gene introns directly alter gene regulation or splicing in neurodevelopmental contexts.

In conclusion, our study provides—to the best of our knowledge—the most comprehensive analysis of the RNA-DNA contactome in a human neurogenesis model, and highlights TIRs as a novel class of caRNAs that shape the 3D genome architecture of neuronal cells. We provide evidence of the existence of a new nuclear compartment—**introsomes**—which, in neurons, is mainly composed of RNAs derived from very long introns, potentially explaining why many neuronally expressed genes have evolved to carry ultra-long introns. Our work suggests that introsomes might not simply represent static structural elements of the nucleus, but rather dynamic self-renewing scaffolds that are continuously regenerated through active transcription and degradation, organizing the space between chromosomes and, as a consequence, globally arranging the genome to support the execution and maintenance of cell type-specific gene expression programs. This introduces, in our view, a form of transcription-dependent nuclear plasticity, where genome organization is not only dynamic, but directly coupled to the transcriptional activity of functionally relevant genes. What remains unanswered is why neurons require such nuclear matrix for their physiology.

## Acknowledgements

We acknowledge the Laboratory for Genotyping Development at the RIKEN Center for Integrative Medical Sciences (IMS) in Yokohama, Japan, the SciLifeLab National Genomics Infrastructure in Stockholm, Sweden and the National Facility for Genomics at Human Technopole in Milan, Italy for support with sequencing. The Hi-C, RNA-seq, and GPSeq data pre-processing was enabled by resources provided by the National Academic Infrastructure for Supercomputing in Sweden (NAISS) at UPPMAX (project no. 2024/22-849) funded by the Swedish Research Council (grant no. 2022-06725). We thank Nobuyuki Takeda and Teruaki Kitakura (RIKEN) for their support of the IT infrastructure for the FANTOM6 collaboration, and Emi Ito (RIKEN) for administrative support. This was a collaborative work with FANTOM6 Consortium. We thank all consortium members for their insights and suggestions. We additionally thank: Kevin Creamer (J.B. Lawrence lab) for guidance and technical assistance with the chromatin fractionation protocol; Francesco Sebastiano Rusconi and Elena Battaglioli (University of Milan) for critically reading the manuscript and providing valuable feedback; Nicola Pirastu (Human Technopole) for providing valuable suggestions on accessing and analyzing publicly available GWAS datasets. We also thank the Simons Foundation for granting us access to the SPARK/SSC cohorts and to all the participants in these studies.

W.H.Y. was supported by a PhD fellowship from the European Union’s Horizon 2020 Research and Innovation programme under the Marie Skłodowska-Curie Actions Innovative Training Network (MSCA ITN ‘Cell2Cell’ – grant no. 860675). B.A.M.B. was supported by a postdoctoral scholarship from the Karolinska Institutet Strategic Programme in Neurosciences (StratNeuro). Y.Z. was supported by a China Scholarship Council. This work was funded through research grants from:

- The Ministry of Education, Culture, Sports, Science and Technology (MEXT) Japan to RIKEN IMS in support of FANTOM6;
- The Swedish Brain Foundation (Hjärnfonden, grant no. PS2023-0023) to W.K.;
- Karolinska Institutet (KI Consolidator Grants 2025), the Swedish Research Council (Consolidator Grant no. 2023-02111), and the Swedish Foundation for Strategic Research (grant no. FFL18-0104) to K.T.;
- The Swedish Research Council (VR, grant no. 2022-00721) and Karolinska Institutet (KI Consolidator Grants 2020) to N.C.;
- The RIKEN Center for Integrative Medical Sciences; the Japanese Ministry of Education, Culture, Sports, Science and Technology (MEXT) to P.C.;
- The Swedish Research Council (grant. no. 2020-02657), Karolinska Institutet (KI Consolidator Grants 2020), and the European Union (ERC, RADIALIS, GA n. 101088408. Views and opinions expressed are those of the authors only and do not necessarily reflect those of the European Union or the European Research Council Executive Agency. Neither the European Union nor the granting authority can be held responsible for them.) to M.B.;
- The European Union (Next Generation EU, MISSION 4, COMPONENT 2, ‘From Research to Business’, INVESTMENT 1.4, ‘Strengthening research infrastructures and creation of national R&D champions’ in relation to the project identified by code CN00000041_S6_LA_006, titled ‘National Center for Gene Therapy and Drugs based on RNA Technology and CUP CNR B83C22002860006’) to P.C. and M.B.

## Author Contribution Statement

*Conceptualization: W.K., P.C., M.B. Data curation: W.K., R.P., W.H.Y., Q.V., Y.Z, M.O, F.M., B.A.M.B., L.S., K.T., M.B., Sample preparation: W.H.Y., X.L., K.Y., M.M., M.K. Sequencing data acquisition: W.H.Y., X.S., M.K., C.W.Y., M.G. FISH validation and analysis: X.L., A.A.Y., Q.V. Knockdown experiments and analysis: K.Y., C.W.Y., R.P., Formal analysis: W.K., R.P., Q.V., L.S., Y.Z, M.O. Funding acquisition: K.T., N.C., P.C., M.B. Investigation: W.K., W.H.Y., Q.V., B.A.M.B., K.T., N.C., P.C., M.B. Project administration: B.A.M.B., M.O., F.M., N.C., M.B. Supervision: T.K., H.T., J.W.S., K.T., P.C., N.C., M.B. Visualization: W.K., L.S., Y.Z., Q.V., B.A.M.B., N.C., M.B. Writing: W.K., B.A.M.B., N.C., M.B. with contributions from all the authors*.

## Methods

### Ethical regulation statement

The collection of samples and data related to the SPARK and SSC autism cohorts described in Computational Methods, section 16 adhered to the ethical standards of the Helsinki Declaration, with informed consent obtained from all the study participants and legal guardians or parents for participants unable to provide consent themselves. The Swedish Ethical Committee approved the analysis of these data to be performed in Sweden (ethical permit number: dnr 2020–00400).

### Experimental Methods

#### 1. Cell culture

All the omic assays depicted in **Figure 1a** were performed on a previously described human induced pluripotent stem cell line (hiPSC-i3N)^71^, which carries a doxycycline-inducible NGN2 expression system integrated at the AAVS1 safe harbor locus, as well as on neural stem cells (NSC) and cortical neurons (NEU) differentiated from hiPSC-i3N. The latter was previously derived from the commercially available WTC-11 iPSC line (Coriell, cat. no. GM25256) and generously made available to us by Dr. Michael E. Ward at the National Institutes of Health, Bethesda MA, USA. We cultured hiPSC-i3N cells in StemFit Basic04 Complete Type medium (Ajinomoto, cat. no. SF041-001) on cell culture vessels coated with iMatrix-511 (Nippi, cat. no. AMS.892 012). We changed the medium every other day and passaged the cells when they reached 70–80% confluence. To passage the cells, we rinsed them with 1x DPBS and incubated them with StemPro Accutase Cell Dissociation Reagent (Thermo Fisher Scientific, cat. no. A1110501) at 37 °C for 7 minutes. After centrifugation, we resuspended the cells in the StemFit medium supplemented with 10 μM Y-27632 (STEMCELL Technologies, cat. no. 72304, or Sigma-Aldrich, cat. no. SCM075) and seeded them on iMatrix-511 coated cell culture vessels. We refreshed the medium the next day, after cell attachment.

For biochemical fractionation experiments, we used SH-SY5Y cells (ATCC, cat. no. CRL-2266) and HCT116 cells (ATCC, cat. no. CCL-247). Briefly, we cultured SH-SY5Y cells in Iscove’s Modified Dulbecco’s Medium (Euroclone, cat. no. ECM0192L) supplemented with 10% fetal bovine serum (Gibco, cat. no. A56707-01), 4 mM L-Glutamine (Gibco, cat. no. 25030-081), 1% Penicillin/Streptavidin (Gibco, cat. no. 15140-122) and 1% Insulin-Transferrin-Selenium (Gibco, cat. no. 41400-045). For HCT116, we used Dulbecco’s Modified Eagle Medium (Euroclone, cat. no. ECM0090L) supplemented with 10% fetal bovine serum and 1% Penicillin/Streptavidin.

All the cell lines used in this study were cultured at 37 °C in 5% CO_2_ air and periodically confirmed to be negative for Mycoplasma contamination. For cryopreservation of iPSC and NSC we used STEM-CELLBANKER (Zenogen, cat. no. 11924). None of the cultures used to produce the data described in this study is included in the International Cell Line Authentication Committee (ICLAC) database of misidentified cell lines.

#### 2. Differentiation of iPSC to neural stem cells and cortical neurons

To differentiate iPSC-i3N to cortical neurons (NEU), we first differentiated them to NSC using PSC Neural Induction Medium (Thermo Fisher Scientific, cat. no. A1647801). We seeded 10^6^ cells (Day 0) on a 10-cm cell culture dish coated with iMatrix-511 (Nippi, cat. no. AMS.892 012) and cultured the cells in StemFit Basic04 containing 10 μM of Y-27632. The next day (Day 1), we replaced the medium with Neural Induction Medium to initiate the differentiation to NSC. We subsequently refreshed the medium every second day, on Days 3 and 5, adding larger volumes of medium to account for the increased cell density. On Day 6, we dissociated the differentiated NSC with StemPro Accutase Cell Dissociation Reagent (Thermo Fisher Scientific, cat. no. A1110501) and plated them on iMatrix-coated 10-cm culture dishes for further expansion. For this purpose, we seeded 1 million NSC per 10-cm dish (Day 0) in Neural Expansion Medium, composed of 49% Neurobasal Medium (Thermo Fisher Scientific, cat. no. 21103049), 49% Advanced DMEM / F-12 (Thermo Fisher Scientific, cat. no. 12634010) and 2% Neural Induction Supplement (Thermo Fisher Scientific, cat. no. A1647801), supplemented with 10 μM of Y-27632. We then replaced the medium with fresh Neural Expansion Medium without Y-27632 on Days 1 and 3. At Day 5, we either passaged the mature NSC for differentiation to neurons (see below) or harvested them for the omic assays depicted in **Figure 1a**.

To further differentiate NSC to NEU, we followed a previously described protocol^72^. Briefly, we seeded the NSC on cell culture dishes coated with poly-L-ornithine (Sigma Aldrich, cat. no. P3655) and cultured them in Differentiation Medium I, composed (for ∼100 mL) of 48.9 mL DMEM/F-12 medium (Thermo Fisher Scientific, cat. no. 31331093), c48.5 mL Neurobasal Medium (Thermo Fisher Scientific, cat. no. 21103049), 500 μL Non-Essential Amino Acids (NEAA) (Thermo Fisher Scientific, cat. no. 11140050), 500 μL GlutaMAX (Thermo Fisher Scientific, cat. no. 35050061), 500 μL N-2 Supplement (Thermo Fisher Scientific, cat. no. 17502001), 1 mL B-27 Supplement (Thermo Fisher Scientific, cat. no. 17504044), 2.5 μg/mL Human Insulin (Sigma, cat. no. I9278), 2 μM DAPT (Sigma, cat. no. D5942), 50 μM 2-Mercaptoethanol (Thermo Fisher Scientific, cat. no. 21985023), 2 μg/mL Doxycycline hyclate (Sigma, cat. no. D9891), and 5 μg/mL Mouse Laminin (Sigma, cat. no. L2020), supplemented with 10 μM of Y-27632 for the first 24 hours. After three days of induction, we switched the medium to Differentiation Medium II, containing 48.5 mL DMEM/F-12 medium (Thermo Fisher Scientific, cat. no. 31331093), 48.5 mL Neurobasal Medium (Thermo Fisher Scientific, cat. no. 21103049), 500 μL NEAA, 500 μL GlutaMAX™, 500 μL N-2 Supplement, 1 mL B-27 Supplement, 2.5 μg/mL Human Insulin, 2 μM DAPT, 10 ng/mL BDNF (Peprotech 450-02), 10 ng/mL GDNF (Peprotech 450-10), 10 ng/mL NT-3 (Peprotech 450-03), 50 μM 2-Mercaptoethanol, 2 μg/mL Doxycycline hyclate, and 0.5 μg/mL Mouse Laminin. On days 6 and 9 we replaced half of the medium with a fresh batch without doxycycline. Lastly, on Day 10, we harvested fully differentiated cortical neurons for the omic assays depicted in **Figure 1a**.

For DNA and RNA FISH experiments on NSC, we seeded NSC on iMatrix-coated, 18-mm glass coverslips (Marienfeld, cat. no. 630-2200) placed in 12-well plates and cultured the cells for 2-3 days in Neural Induction Medium, supplemented with 10 μM Y-27632 during the first 24 hours, before fixation. For FISH on NEU, we first seeded NSC on poly-L-ornithine coated coverslips and differentiated them as described above.

#### 3. RADICL-seq

We performed RADICL-seq on iPSC, NSC and NEU as previously described^35^. Briefly, we fixed 1 million iPSC, NSC and NEU cells in 1% paraformaldehyde (PFA) (Thermo Fisher Scientific, cat. no. 28906) solution in 1x PBS and stored the cell pellets at –80 °C after FA quenching and washing. For the proximity ligation of RNAs and chromatin, we resuspended the cell pellets in a lysis buffer containing 10 mM Tris-HCl pH 8.0, 10 mM NaCl, and 0.2% NP-40, followed by a mild digestion with DNase I (Thermo Fisher Scientific, cat. no. EN0525) at 37 °C for 10 minutes. We then performed end-repair with T4 DNA Polymerase (Thermo Fisher Scientific, cat. no. EP0062) and DNA Polymerase I, Large (Klenow) Fragment (Thermo Fisher Scientific, cat. no. 18012021) at room temperature for 1 hour followed by A-tailing using Klenow Fragment (3’→5’ exo–) (Thermo Fisher Scientific, cat. no. EP4021) at 37 °C for 1 hour. We added RNase H (New England BioLabs, cat. no. M0297L) to the reaction and incubated at 37 °C for 40 minutes. We then ligated the pre-adenylated RADICL-seq adapter to the RNA using T4 RNA Ligase 2, truncated KQ (New England BioLabs, cat. no. M0373L) at 20 °C overnight. The next day, we performed ligation of the adapter-ligated RNA to chromatin by incubating the cell pellets with T4 DNA Ligase (New England BioLabs, cat. no. M0202L) at room temperature for 4 hours. We then proceeded to extract genomic DNA and prepare sequencing libraries as described in ref. ^35^. We sequenced each RADICL-seq library separately on the NovaSeq 6000 platform (Illumina) using the S2 Reagent Kit v1.5 (200 cycles) (Illumina, cat. no. 20028315) with SE150 sequencing mode, aiming at generating 600 million reads per sample.

#### 4. Hi-C

We performed Hi-C on iPSC, NSC, and NEU using the Arima HiC+ kit (Arima Genomics, cat. no. A510008) and the Arima Library Prep Module (Arima Genomics, cat. no. A303011). Briefly, we fixed 2 million iPSC, NSC and NEU cells in 2% paraformaldehyde (PFA) (Thermo Fisher Scientific, cat. no. 28906) solution in 1x PBS and stored the cell pellets at –80 °C after PFA quenching and washes. We then performed in situ proximity ligation and library preparation following the manufacturer’s protocol. We sequenced the libraries on the NovaSeq 6000 platform (Illumina) using 2 lanes and the S4 Reagent Kit v1.5 (300 cycles) (Illumina, cat. no. 20028312) with PE150 sequencing mode, aiming at generating 800–1,000 million reads per library.

#### 5. GPSeq

We performed GPSeq on iPSC, NSC and NEU following an improved GPSeq protocol that we recently described^52^. Briefly, for each cell type, we fixed 0.3 million cells on 22 x 22 mm coverslips in 4% paraformaldehyde (PFA) solution (Thermo Fisher Scientific, cat. no. 28906) in 1x PBS. We performed radial chromatin digestion using DpnII for 6 different timepoints (10 sec, 30 sec, 2 min, 5 min and 30 min for NES; 10 sec, 30 sec, 2 min, 5 min, 10 min and 30 min for NPC; 10 sec, 20 sec, 30 sec, 45 sec, 2 min and 5 min for NEU). To visually confirm that radial digestion had occurred properly, we performed YFISH and analyzed the images as described in ref. ^37^. We prepared one sequencing library for each timepoint and sequenced all the libraries on the NextSeq 500/550 platform (Illumina) using High Output Kit v2.5 (75 Cycles) (Illumina, cat. no. 20024906) with SE75 sequencing mode, aiming at generating 50 million reads per library.

#### 6. scATAC-seq

We isolated nuclei from iPSC, NSC and NEU following the 10x Genomics protocol for nuclei extraction for scATAC-seq (10x Genomics, Demonstrated Protocol no. CG000169). We performed DNA tagmentation, single-cell droplet encapsulation, and cell indexing using the Chromium Next GEM Single Cell ATAC Library & Gel Bead Kit (10x Genomics, cat. no. 1000175), aiming at processing 5000 nuclei. We sequenced all the libraries on the HiSeq X platform (Illumina) using one lane of HiSeq X Ten Reagent Kit v2.5 (Illumina, cat. no. FC-501-2501) with PE150 sequencing mode, aiming at generating 150 million reads per library.

#### 7. Total RNA-seq

We extracted total RNA from iPSC, NSC and NEU using the Qiagen RNeasy Mini kit (Qiagen, cat. no. 74104) and prepared libraries using the TruSeq Stranded Total RNA Sample Prep Kit with Ribo-Zero Human/Mouse/Rat kit (Illumina, cat. no. 20020596) and TruSeq RNA Single Indexes (12 indexes, 24 samples) Set A (Illumina, cat. no. 20020492). We sequenced all the libraries on the NovaSeq 6000 platform (Illumina) using 1 lane of S4 Reagent Kit v1.5 (300 cycles) (Illumina, cat. no. 20028312) with PE150 sequencing mode, aiming at generating 300–400 millions reads per library.

#### 8. Cap Analysis of Gene Expression (CAGE)

We extracted total RNA in the same way as for total RNA-seq and prepared single-stranded CAGE libraries as previously described^39^. We sequenced all the libraries on NextSeq 1000/2000 using the P2 Reagent Kit (Illumina, cat. no. 20100986) with PE100 sequencing mode, aiming at generating 50 million reads per library.

#### 9. Immunofluorescence for neuron-specific markers

To confirm that iPSCs were properly differentiated into NSC and NEU, we performed immunofluorescence staining with the following antibodies: Rabbit NeuN Recombinant Antibody (14HCLC) (Thermo Fisher Scientific, cat. no. 711054); Guinea Pig MAP2 antibody (CiteAb, cat. No. 188004); Mouse Neuron-specific beta III Tubulin Antibody (Bio-techne R&D system, cat. No. MAB1195); goat anti-Rabbit IgG (H+L) Cross-Adsorbed Secondary Antibody Alexa Fluor 488 (Thermo Fisher Scientific, cat. no. A-11008); Goat anti-Guinea Pig IgG (H+L) highly Cross-Adsorbed Secondary Antibody Alexa Fluor 594 (Thermo Fisher Scientific, cat. no. A-11076); Goat anti-Mouse IgG (H+L) Cross-Adsorbed Secondary Antibody Alexa Fluor 647 (Thermo Fisher Scientific, cat. no. A-21235). We fixed NSC and NEU cells with 4% paraformaldehyde (Fisher Scientific, cat. no. 11481745) for 10 minutes and inactivated the unreacted formaldehyde with 1x PBS (Ambion, cat. no. AM9625), 125 mM glycine (Sigma, cat. no. 50046) for 5 min at room temperature. We then permeabilized the cells with 1x PBS, 0.5% Triton X-100 (Sigma, cat. no. T8787) for 20 minutes at room temperature. After blocking in 1x PBS, 0.1% Tween-20 (Sigma, cat. no. P9416), 5% BSA (Thermo Fisher Scientific, cat. no. AM2618) for 1 hour at room temperature, we incubated the cells in primary antibodies diluted 250 x in 1x PBS, 0.1% Tween-20, 0.1% BSA overnight at 4 °C. The next day, we washed the cells three times in 1x PBS, 0.1% Tween-20 for 5 min at room temperature. We then incubated the cells in the appropriate secondary antibody diluted at 2-4 µg/mL in 1x PBS, 0.1% Tween-20, 5% BSA for 1 hour at room temperature. We washed the cells three times in 1x PBS, 0.1% Tween-20 for 5min at room temperature, followed by incubation in 1xPBS, Hoechst 33342 (Thermo Fisher Scientific, cat. no. H3569) for 15 minutes at room temperature to stain DNA. Lastly, we mounted the coverslips with Prolong Diamond antifade mountant medium (Thermo Fisher Scientific, cat. no. P36961). ***Imaging.*** We acquired fluorescent images on a custom-built Nikon Eclipse Ti2 inverted wide-field microscope, equipped with a Plan Apochromat Lambda 60x/1.4 oil-immersion objective (Nikon) and an Andor iXon Ultra 888 EMCCD camera (Oxford Instruments). For all datasets, we collected z-stacks with 300 nm spacing between imaging planes.

#### 10. RNA FISH, DNA FISH and combined DNA&RNA FISH

##### Probe design and production

We designed DNA/RNA FISH probes targeting various intronic and exonic sequences of TIR source genes and matching TIR control genes; TIRCs and matching TIRC control loci; and chr12 and chr13 spotting probes, using the iFISH pipeline that we previously developed^40^. We appended a probe-specific barcode pair to each oligo and ordered multiple probes as oligopools (Twist Bioscience). The genomic coordinates and oligo sequences of all FISH probes used in this study are available in **Supplementary Table 3**. We amplified individual probes from the oligopools by PCR using the corresponding primers, complemented with a readout sequence at the 5’ end and a T7 promoter sequence at the 3’ end, using the PowerUp SYBR Green Master Mix (Thermo Fisher Scientific, cat. no. A25776). We purified the PCR product with AMPure XP magnetic beads (Beckman Coulter, cat. no. A63882) following the manufacturer’s instructions. We performed *in vitro* transcription (IVT) with the purified PCR product as template, using the HiScribe T7 Quick High Yield RNA Synthesis Kit (New England Biolabs, cat. no. E2050S) in presence of RNaseOUT (Thermo Fisher Scientific, cat. no. 10777019). We purified the RNA product with RNAClean XP magnetic beads (Beckman Coulter, cat. no. A63987) and converted it to single-stranded DNA (ssDNA) by reverse transcription using the Maxima H Minus Reverse Transcriptase (Thermo Fisher Scientific, cat. no. EP0752). Finally, we hydrolyzed the RNA in alkaline conditions and purified the resulting probes using the Monarch PCR & DNA Cleanup Kit (New England Biolabs, cat. no. T1030L).

##### RNA FISH

After growing cells on coverslips, we fixed them in 4% PFA (Fisher Scientific, cat. no 11481745) in PBS for 10 min at room temperature, before quenching with 125 mM glycine (Sigma Aldrich, cat. no. 4810) in PBS for 5 min. We permeabilized the cells using 0.5% Triton X-100 (Sigma Aldrich, cat. no. T8787) in PBS for 20 min at room temperature, followed by two washes with 0.05% Triton X-100 in PBS; or, alternatively, 70% ethanol for 10 min at room temperature. We incubated the cells in 0.1 N HCl (Sigma Aldrich, cat. no. 1090571000) for 5 min at room temperature, followed by two washes with 0.05% Triton X-100 in PBS. We equilibrated the cells in RNA wash buffer, composed of 25% formamide (Ambion, cat. no. AM9344) in 2x saline-sodium citrate (SSC) buffer (Ambion, cat. no. AM9765), for at least 5 min at room temperature, before proceeding with hybridization. We mixed the ssDNA FISH probes at a ratio of 1:99 with RNA hybridization buffer containing 2x SSC, 25% formamide, 10% dextran sulfate (Sigma Aldrich, cat. no. S4030), 1 mg/mL tRNA from *E. coli* (Sigma Aldrich, cat. no. 10109541001), 0.2 mg/mL BSA (Ambion, cat. no. AM2616), and 2 mM Ribonucleoside Vanadyl Complex (RVC; New England Biolabs, cat. no. S1402S), to achieve a final concentration of 1 nM per oligo in the final hybridization mix. We performed hybridization overnight at 37℃ in a humidity chamber. On the next day, we washed the coverslips twice with RNA wash buffer for 30 min at 37℃. Afterwards, we prepared the readout hybridization mix, by combining fluorescently-labeled readout oligos with the readout hybridization buffer containing 2x SSC, 10% dextran sulfate and 25% formamide, at a ratio of 1:99 to achieve a final concentration of 20 nM per readout oligo. For FISH probes targeting exons, we compensated for their relatively low number of oligos by adding a short oligo bridge, composed of two readout sequences and the reverse-complementary sequence to the 3’ barcode of the probe oligos, to the hybridization mix, thereby effectively increasing from 1 to 3 readout docking sites per oligo. We incubated the cells with the readout hybridization mix overnight at 30℃ in a humidity chamber. The next day, we washed the cells with 0.2x SSC, 0.2x Tween-20 (Sigma Aldrich, cat. no. P9416) for 2x 10 min at 45℃, then performed counterstaining with 1 ng/μL Hoechst 33342 (Thermo Fisher Scientific, cat. no. 62249) in 2x SSC for 30 min at room temperature. After washing the cells with 2x SSC, we mounted the coverslips in GLOX buffer, containing 10 mM Trolox (Sigma Aldrich, cat. no. 238813), 37 μg/mL glucose oxidase (Sigma Aldrich, cat. no. G2133), 100 μg/mL catalase (Sigma Aldrich, cat. no. C3515) in an equilibrium buffer with 10 mM Tris-HCl (Invitrogen, cat. no. 15567027), 0.4% glucose (Sigma Aldrich, cat. no. 49139) and 2x SSC, sealed the sides of the coverslips with rubber cement (Fixogum, Triolab, cat. no. LK071A) and proceeded immediately with imaging. Alternatively, we mounted the coverslips in Prolong Glass Antifade Mountant (Thermo Fisher Scientific, cat. no. P36980), let them cure for 24 hours and proceeded with imaging.

##### DNA FISH and combined DNA&RNA FISH

After growing cells on coverslips, we fixed them in 4% PFA in PBS for 10 min at room temperature, before quenching with 125 mM glycine in PBS for 5 min. We permeabilized the cells using 0.5% Triton X-100 in PBS for 20 min at room temperature, followed by two washes with 0.05% Triton X-100 in PBS. We incubated the cells in 0.1 N HCl for 5 min at room temperature, followed by two washes with 0.05% Triton X-100 in PBS. For labelling of the *NALF1* source locus, we additionally incubated the cells for 1 hour with a mix of RNase Cocktail Enzyme Mix (Invitrogen, cat. no. AM2286) and RNase I (Thermo Scientific, cat. no. EN0601), for a final concentration of 70 U/mL RNase A, 285 U/mL RNase T1 and 143 U/mL RNase I in PBS, before proceeding with the rest of the protocol. We equilibrated the cells in FPS buffer composed of 50% formamide, 50 mM sodium phosphate (Thermo Fisher Scientific, cat. no. J60158-AP) and 2x SSC buffer overnight at room temperature. The next day, we incubated the cells for 1 hour in the pre-hybridization buffer containing 50% formamide, 5x Denhardt’s solution, 1 mg/mL tRNA from *E. coli*, 100 μg/mL salmon sperm DNA (Invitrogen, cat. no. 15632011), 0.2 mg/mL BSA and 2x SSC, before proceeding with hybridization. We mixed the ssDNA FISH probes at a ratio of 1:10 with DNA hybridization buffer, composed of 2.2x SSC, 5.5x Denhardt’s solution, 11% dextran sulfate and 55% formamide, to achieve a final concentration of 0.05 nM per oligo for FISH probes against DNA, and 1 nM per oligo for FISH probes against RNA. We placed the coverslips with cells face-down on microscope slides with 10 μL of the hybridization mix and sealed the coverslips with rubber cement. We performed DNA denaturation at 75–78℃ for 2-3 min on a heating block, then transferred the slides to a humidity chamber for hybridization overnight at 37℃. On the next day, we released the coverslips in 2x SSC, 0.2% Tween-20, then washed the samples twice for 7 min at 56–65 ℃ in 0.2x SSC, 0.2% Tween-2. We rinsed the coverslips with 4x SSC, 0.2% Tween-20, followed by 2x SSC, before equilibrating the cells in RNA wash buffer, composed of 25% formamide in 2x SSC, for at least 5 min at room temperature. Afterwards, we prepared the readout hybridization mix, by combining fluorescently-labeled readout oligos with the readout hybridization buffer containing 2x SSC, 10% dextran sulfate and 25% formamide, at a ratio of 1:99 to achieve a final concentration of 20 nM per readout oligo. We incubated the cells with the readout hybridization mix overnight at 30℃ in a humidity chamber. The next day, we washed the cells twice with 0.2x SSC, 0.2x Tween-20 for 2x 10 min at 45℃, then performed counterstaining with 1 ng/μL Hoechst 33342 in 2x SSC for 30 min at room temperature. After washing the cells with 2x SSC, we either mounted the coverslips in GLOX buffer, sealed the sides of the coverslips with rubber cement and proceeded immediately with imaging; or mounted the coverslips in Prolong Glass Antifade Mountant, let this cure for 24 hours, and proceeded with imaging.

##### Imaging

We acquired fluorescent images on either a custom-built Nikon Eclipse Ti inverted wide-field microscope, equipped with a Plan Apochromat Lambda 100x/1.45 oil-immersion objective (Nikon) and an Andor iXon Ultra 888 EMCCD camera (Oxford Instruments), or on a custom-built Nikon Eclipse Ti2 inverted wide-field microscope, equipped with a Plan Apochromat Lambda 100x/1.45 oil-immersion objective (Nikon) and an Andor Sona sCMOS camera (Oxford Instruments). For all datasets, we collected z-stacks with 200-300 nm spacing between imaging planes.

#### 11. Cellular fractionation and RNA-seq

We performed cellular fractionation on 3×10^7^ SH-SY5Y cells and 3×10^7^ HCT116 cells, according to a previously published protocol^26^ with the following minor modifications. Briefly, after collecting the cytoplasmic (CYTO) and nuclear soluble (NS) fractions, we resuspended the remaining material in Ammonium Sulphate Extraction Buffer and collected the chromatin-dependent nuclear soluble fraction (CDNS). We then resuspended the insoluble nuclear material in Trizol (Invitrogen, cat. no. 15596018), 10 mM EDTA (final concentration). We precipitated the CYTO, NS and CDNS fractions with isopropanol and recovered the RNA by adding Trizol, 10 mM EDTA. Before RNA isolation, we heated all the fractions at 65 ℃ for 15 minutes. Lastly, we treated the Trizol-isolated RNA with RNase-Free DNase Set (Qiagen, cat. no. 79254) and used the resulting DNA-free RNA to prepare RNA-seq libraries using the Stranded Total RNA Prep, Ligation with Ribo-Zero Plus (Illumina, cat. no. 20040529). We sequenced all the libraries on the NovaSeq 6000 platform (Illumina) using the NovaSeq 6000 SP Reagent Kit v1.5 (200 cycles) (Illumina, cat. no. 20040719) with paired-end 100 bp sequencing mode, aiming at generating 30 million reads per sample.

### Computational Methods

The following sections are ordered based on when the corresponding analyses are first described in the Results.

#### 1. RNA-seq data analysis

We processed and aligned RNA-seq data from iPSC, NSC, and NEU using the *nf-core/rnaseq* pipeline (v3.10.1). We aligned the reads to the human GRCh38/hg38 reference genome with GENCODE v38 as gene annotation. We used the tximport (v 1.18.0) R package to generate a raw gene count table based on salmon quant.sf output files obtained from nf-core/rnaseq pipeline. We obtained the transcript-per-million (TPM)-normalized gene expression matrix (*salmon.merged.gene_tpm.tsv*) from nf-core/rnaseq pipeline. We performed differential gene expression analysis with *DESeq2* (v1.30.1) using the raw gene count table as input. We used the TPM-normalized gene expression matrix and the *DESeq2* rlog-transformed counts for visualization purposes and to plot all the corresponding figure panels.

#### 2. Gene ontology analysis

We performed gene ontology (GO) analysis using *DOSE* (v3.28.1), *clusterProfiler* (v4.10.0) and *org.Hs.eg.db* (v3.18.0) R packages. We retained only findings related to biological processes and visualized the results using *ggplot2* (v 3.5.1). For GO analysis on NEU samples, we defined as expressed the genes being expressed at levels higher than the mean expression of all protein-coding (pc) genes (DESeq rlog ≥ 9.6, corresponding to ∼0.4 quantile of all pc genes in NEU Replicate 1).

#### 3. RADICL-seq data analysis

We processed and mapped the RADICL-seq data as previously described^35^. In brief, we mapped the data to the human GRCh38/hg38 reference genome. Since correlation analysis showed a high degree of similarity between biological replicates, we merged the two replicates from each cell line. We assigned RNA tags to annotated genes while preserving strand specificity, and grouped DNA tags into 25 kb genomic bins, with each fragment represented by its central nucleotide to minimize overlap with multiple bins. We excluded all blacklisted genomic regions in the ENCODE hg38 genome blacklist and filtered out duplicate RNA-DNA contacts using the *BEDtoolsr* R package. To determine statistically significant RNA-DNA contacts, we used a modified negative binomial distribution model implemented in *CHiCANE*^73,74^. We retained only significant contacts that met a q-value threshold of 0.05 (q ≤ 0.05) for further analysis. We summed contacts of a gene in the same 100 kb genomic bin using *intersect* and *groupby* in *bedtools* (v2.30.0). Since correlation analysis showed a high degree of similarity between biological replicates, we merged the two replicates from each cell line. We classified the significant RNA-DNA contacts as either cis—if the RNA contacts a DNA locus within 5 Mb of its source locus (on the same chromosome)—or trans—if the contacted DNA locus was found beyond 5 Mb or on a different chromosome. To compute the speckle proximity score, we used RPM-normalized RADICL-seq contact frequencies within 100 kb genomic bins contacted by *MALAT1* RNA.

#### 4. Hi-C data analysis

We first processed the Hi-C data following Arima Genomics’ recommendations to trim 6 nt from the left end, and 70 nt from the right end of each read using `trimfq –b 6 –e 70` in the *seqtk* tool (https://github.com/lh3/seqtk). We then fed the remaining 75 nt from each read to the *nf-core hic* pipeline (v 2.1.0, https://nf-co.re/hic/2.1.0/) with the following parameters:

`--genome ‘hg38’`

`--restriction_site ‘[^GATC,G^ANTC]’`

`--ligation_site ‘[GATCGATC,GANTGATC,GANTANTC,GATCANTC]’`

`--digestion ‘arima’`

`--res_compartments ‘500000,250000,100000’`

`--tads_caller ‘insulation,hicexplorer’`

`--bin_size ‘1000000,500000,100000,50000,25000,10000,5000,2500,1000’`

`Since correlation analysis showed a high degree of similarity between biological replicates, we merged the two replicates from each cell line.`

To call A/B compartments, we processed the Hi-C contact maps (100 kb resolution) generated by the *nf-core hic* pipeline using a customized R script (https://github.com/wenjingk/TIR_code/tree/main/Hi-C_analysis). Briefly, we loaded and converted the ICE normalized contact map to an observed versus expected (O/E) intra-chromosomal matrix. For this, we first calculated the expected Hi-C signal for intra-chromosomal contact maps using *cooltools expected-cis* (v 0.6.1), with the balanced cool file at 100 kb resolution as input and with the options `--smooth –-aggregate-smoothed`. For each intra-chromosomal DNA-DNA contact, we then calculated the O/E value by dividing the ICE normalized value by the expected value c of the given linear genomic distance between the two DNA loci. We converted the O/E matrix to a correlation matrix using R *cor* with the `use=“pairwise.complete.obs”` option and performed Principal Component Analysis (PCA) on the correlation matrix using R *prcomp*, where the correlation matrix is filtered by removing DNA bins with missing values. After recording the PCA scores, we corrected the sign of the PC1 value when needed to obtain a positive correlation with GC-content, so that DNA bins with positive sign represent the A compartment, while DNA bins with negative sign represent the B compartment. We obtained the GC-content at 100 kb resolution using `genome gc` in *cooltools* with the coordinates of 100 kb genomic bins and the human reference genome hg38 as input.

To obtain O/E values for inter-chromosomal contact maps, we leveraged *cooltools expected-trans* with the balanced cool file (1 Mb resolution) as input, and with the default options. The O/E values are calculated by dividing the ICE normalized value by the expected value (the ‘balanced.avg.smoothed’ reported from the *cooltools expected-trans*) of the given chromosome pairs.

#### 5. CAGE data analysis

We de-multiplexed the CAGE reads and mapped them to the human GRCh38/hg38 reference genome using STAR^75^. We retained the primary alignments of Read1 with mapping quality (MAPQ) > 10 and extracted their 5′ ends as single-nucleotide transcription start sites (TSS count). To define a set of universal promoter regions, we collected TSS from a comprehensive transcriptome annotation combining GENCODE v39, FANTOM CAT^76^, and CFC-seq-derived models from iPSC, NSC, NEU, and THP-1 cells^77^. To define promoter regions, we extended the 5′ ends of all these transcript models 100 nt upstream and 25 nt downstream in a strand-specific manner, which were merged if overlapping. To generate a promoter-level count matrix, we counted the TSS within these promoter regions (n = 282,958). Subsequently, we aggregated the TSS counts to the gene level based on the annotation linking promoter regions to 174,933 genes excluding ribosomal RNAs. Lastly, we log-normalized the counts across all the CAGE libraries into counts per million (CPM) mapped reads.

#### 6. RNA-seq and RADICL-seq RNA coverage over the source gene body

We identified significant RNA-DNA contacts retained after *CHiCANE* filtering and used them to generate RADICL-seq RNA coverage profiles (bedGraph format) over the genome using *bedtools genomecov* (v 2.29.2). We then converted the RNA-seq RNA coverage from the bigWig files generated by *nf-core rnaseq* pipeline to bedGraph format using *bigWigToBedGraph*. For each gene of interest, we first generated sliding windows with window size 100 bp and step size 10 bp using the *makewindows* tool in *BEDTools* and we annotated them based on whether the windows cover exons (including 3′ and 5′ UTRs) or introns. We then intersected the windows with the RADICL-seq and RNA-seq RNA coverage profiles, and summed the coverage counts falling in each window. For each gene, we used the maximum coverage count of each exon and intron to calculate the intron vs. exon coverage ratio, which is equal to the mean of the maximum coverage counts for introns divided by the mean of the maximum coverage counts for exons.

#### 7. Intron length analysis

We obtained intron coordinates from the GENCODE v38 GTF annotation and defined intronic regions by subtracting annotated exons from the corresponding gene bodies using *BEDTools*. To annotate the RNA component, we intersected the RNA coordinates of significant RADICL-seq RNA-DNA pairs obtained from the FANTOM6 repository with intron coordinates. We assigned the DNA component of each pair to 100 kb genomic bins. We then collapsed RNA-DNA pairs by intronic regions and summed the DNA contact counts for each genomic bin.

For each of the top-55 TIR genes identified in the RADICL-Seq neuron dataset, we applied a custom R function to quantify and classify the relationship between intron length and trans-contact features, including the total number of trans contacts and the number of distinct trans-contacted DNA bins (at 100 kb resolution) formed by the corresponding intronic RNAs, into distinct growth modes. The function fitted several candidate models to each gene’s data using least-squares regression (lm or nls), including linear (*y* = *a* + *bx*), saturating (*y* = *a*(1 – *e*^-*bx*^)), exponential (*y* = *ae^bx^*), and power-law (*y* = *ax^k^*) forms, with starting parameters determined automatically. Model performance was evaluated using the Akaike Information Criterion (AIC), which balances fit accuracy against model complexity, and the model with the lowest AIC was selected as the best representation of the relationship. From the best-fitting model, an Acceleration Factor was derived as the ratio of predicted slopes at the 90th and 10th percentiles of intron length, providing a quantitative measure of how growth rate changes across the length range. An Acceleration Factor greater than 1 indicates an accelerating trend, a value near 1 indicates constant growth, and a value less than 1 indicates a saturating trend.

#### 8. FISH image analysis

##### Deconvolution

We processed all FISH datasets using our deconvolution package *Deconwolf*^41^ (v0.4.5, available at https://deconwolf.fht.org/). Briefly, we generated in silico point-spread functions (PSF) for each of the fluorescent dyes used for FISH as well as for Hoechst 33342 used for DNA staining, with the imaging parameters matching the dataset to process. As an example, we generated a PSF for images of ATTO 647, with an emission maximum at 662 nm, and imaged with an oil-immersion objective with NA 1.45, pixel size of 65 nm, and 200 nm between z-planes at acquisition, using the following parameters:

`dw_bw –-lambda 662.0 –-NA 1.450 –-ni 1.515 –-threads 4 –-resxy`

`65.0 –-resz 200.0 ‘./ATTO647_PSF.tif’`

We applied the obtained PSF to deconvolve the corresponding datasets, by processing single-color channels individually, then splitting the image stack in four along the x-y axis and using GPU acceleration to process the stacks separately before recombining them at output. As an example, we ran *Deconwolf* on an ATTO 647 image stack using the following settings, for 50 iterations with GPU acceleration:

`dw –-bq 2 –-method shbcl2 –-tilesize 1024 –-iter 50 –-threads 4`

`‘image_path_ATTO647.tif’ ‘./ATTO647_PSF.tif’`

For visualization, we used 50 iterations of *Deconwolf* to deconvolve all the images shown in the figures. For nucleus segmentation and identification of dots, we used images deconvolved for 50–100 iterations, to reduce background signals without reaching oversharpening.

##### Nuclei segmentation and dot detection

We segmented cell nuclei using either thresholding of the Hoechst 33342 channel, or manual annotation. We detected FISH dots using the ‘`df_getDots`’ function in our DOTTER^78^ analysis suite, using default values. Briefly: (i) We applied a Difference-of-Gaussians filter to the image; (ii) We extracted local maxima to identify possible dots of interest; (iii) We set a manual threshold with visual feedback, to separate false from true localization events; and (iv) We fitted a Gaussian model on the remaining dots using Maximum Likelihood, to retrieve sub-pixel localization. For intron and exon quantifications, we obtained dot counts per field of view (130 x 130 μm^2^) and normalized them to the number of nuclei. To estimate the relative proximity of loci to nuclear speckles, we identified DNA FISH dots within manually corrected nuclei masks, then measured the average *MALAT1* fluorescence intensity within 0.5 μm around each dot compared to 2.5 μm away from it and reported this ratio as the local enrichment of *MALAT1* transcripts.

##### Co-localization analysis

To quantify the extent of overlap in 3D between the top-10 TIRs identified in NEU and TIRCs or TIRC-control regions (see **Supplementary Fig. 25d, e**), we first deconvolved the images using 100 iterations of *Deconwolf* (v0.4.6 available at https://deconwolf.fht.org/). We detected FISH dots using the dot detection module incorporated in *Deconwolf* with manually set channel-specific thresholds based on visual inspection of the images. The final thresholds used were: i) 4.21 × 10³ for the top-10 TIRs (A647 channel); 1.75 × 10³ for TIRCs (A594 channel); and 4.37 × 10³ for TIRC-control regions (SpGold channel). To estimate the underlying A647 signal at the positions of the FISH dots detected in the other channels, we applied lateral smoothing on the A647 images using a Gaussian filter (σ = 4). We then interpolated the smoothed A647 images at the coordinates of the FISH dots in the A594 and SpGold images to generate the density plots shown in **Supplementary Fig. 25d, e**.

#### 9. RADICL-seq residue calculation

To calculate RADICL-seq residues for each source gene of interest, we summed up the RADICL-seq contacts made by RNA derived from the source gene, per 100 kb DNA bin. We scaled the binned contacts from the chromosome harboring the source gene using Min-Max normalization and extracted contacts from the region ranging from 5 Mb upstream to 5 Mb downstream to the highest peak of RNA-DNA contacts at the source gene. The middle point of the plot is defined by the genomic bin with the highest contacts over the gene body. To generate the background decay model, we applied a generalized linear model (GLM) with a Gamma family (`glm` function in R) to separately fit the binned contacts at two regions: from the midpoint of the source gene to 5 Mb upstream; and from of the midpoint of the source gene to 5 Mb downstream. We then measured the residues by taking the binned contact counts and subtracting the number of contacts that are expected by the decay background model. Residues were then scaled back to 0.0-1.0 for displaying purposes.

#### 10. HNRNPU fRIP-seq data analysis

To determine whether specific gene sets were enriched among the genes previously identified by RNA-seq following HNRNPU Formaldehyde crosslinking and ribonucleoprotein immunoprecipitation (fRIP-seq)^51^, we performed hypergeometric enrichment analyses using the R function `phyper`. Gene sets included the top-55 TIR source genes identified in NEU as well as autism-associated genes (scores 1, 2, and syndromic) from the SFARI database (release 08-19-2024)^79^, gene panels for intellectual disability and epilepsy (green and amber classifications) from *PanelApp* (https://panelapp.genomicsengland.co.uk/; versions 7.0 and 6.0, respectively). We defined the background as all the genes expressed in the corresponding sample based on RNA-seq data from ref. ^80^. We calculated the enrichment using a hypergeometric model that estimates the probability of observing an overlap equal to or greater than the observed count when a gene set of the same size is randomly drawn from the background. Upper-tail *P* values were computed as:

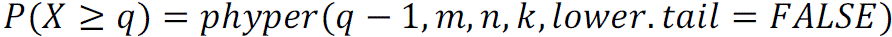

where *m* and *n* represent, respectively, the number of background genes that do and do not belong to the gene set of interest; *k* is the number of HNRNPU-associated genes; and *q* is the observed number of genes included both in the gene set of interest and in the set of HNRNPU-associated genes. A significant *P* value indicates that the gene set is overrepresented among HNRNPU-associated genes relative to chance. We applied this procedure separately to each gene list.

#### 11. Cellular fractionation and RNA-seq data analysis

We processed and aligned the RNA-seq reads from different cellular fractions of HCT116 and SH-SY5Y cells using the *nf-core/rnaseq* pipeline (v3.10.1). We aligned the reads to the human GRCh38/hg38 reference genome, with gene annotations from GENCODE (v38). We quantified read counts using *featureCounts* (v2.1.1) at three different genomic resolutions: (i) gene level, where the reads were assigned to full gene bodies; (ii) exon-intron level of protein-coding genes, where all the annotated exons and introns per gene were concatenated and counted as separate features; and (iii) individual intron level, where each annotated intron of protein-coding genes was quantified independently. We obtained the coordinates of genes, exons, and introns from the GTF annotation in GENCODE (v38). We defined intronic regions by subtracting the annotated exons from the corresponding gene body using *BEDTools* and the BAM files generated by the *nf-core/rnaseq* pipeline as input.

#### 12. scATAC-seq data analysis

We processed the scATAC-seq data using *cellranger-atac-2.0.0*, which yields three files per differentiation stage: 1) ‘peaks.bed’ contains the coordinates of scATAC-seq peaks; 2) ‘matrix.mtx’ contains the peak index, cell index and peak abundance; and 3) ‘barcodes.tsv’ contains the barcode of single cell. We obtained the coordinates of scATAC-seq peaks from ‘peaks.bed’ and calculated the height of the pseudo-bulk ATAC-seq peaks based on the number of cells sharing the peak by parsing ‘matrix.mtx’. We then assigned the pseudo-bulk ATAC-seq peaks to the 100 kb DNA bins over the genome using *bedtool intersect* and used the average height of the peaks mapped to the same DNA bin to represent the abundance of pseudo-bulk ATAC-seq signals per bin.

#### 13. GPSeq data analysis

##### GPSeq data preprocessing

We processed the GPSeq data using a customized pipeline available at https://github.com/wenjingk/GPSeqNP. We installed all necessary software and packages in a docker and singularity container. Briefly, we indexed the reference genome (GRCh38) using *bowtie2* (v 2.4.1) and identified the coordinates of the recognition site (GATC) of the restriction enzyme DpnII in the reference genome using *fastx-barber* (v 0.1.5). We then identified the 8-nt unique molecular identifier (UMI), 8-nt sample barcode, and 4-nt cut site in subsequent order from the 5′ end of the reads. We trimmed these sequences and stored them and their corresponding sequencing quality scores in the header, using *fastx-barber*. We discarded reads if the barcode or the cut site sequence did not match the expected sequence with up to 1 mismatch, or if more than 20% of the nucleotides in the UMI had a quality score below 30. We aligned the remaining reads to the reference genome using *bowtie2* with options `--very-sensitive –L 20 –-score-min L,-0.6,-0.2 –-end-to-end`. We retained only primary alignments with a mapping quality equal to or higher than 30, while discarding unmapped, multimapping, chimeric, chrM, and low mapping quality reads. Next, we grouped reads mapped to the same genomic coordinate and calculated the distance between that coordinate and the nearest cut site using the customized script *group_umis.py*. We discarded read groups that could not be confidently assigned to a nearest cut site (threshold for assigning within 20 nucleotides), using the customized script *umis2cutsite.py*. Finally, we removed duplicated reads at each cut site if they had the same UMI, using the customized script *umi_dedupl.R*. These preprocessing steps resulted in a bed file containing the number of unique reads found at each cut site, for each digestion time point.

##### GPSeq score and percentile calculation

To calculate the GPSeq score^37^, we input the bed files generated from the preprocessing and a config file sample information to the *gpseq-radical.R* script (v 0.0.9) with the following parameters:

--normalize-by lib

--chromosome-wide

--binsize 1e6:1e5,5e5:5e4,1e5:1e4,5e4:5e4,25e3:25e3

--ref-genome hg38

--chrom-tag 22:X,Y

--site-domain universe

--mask-bed “blacklist.bed”

The blacklist file contains regions with low mappability (< 80% coverage of k50 UMAP data, https://bismap.hoffmanlab.org/), including highly repeated regions (e.g., centromeres). We excluded these regions for the GPSeq score calculation. To enable comparison across cell types and account for variable ranges of the GPSeq score observed in different experiments, we calculated the GPSeq percentile as: (# of genomic bins with score less than or equal to the selected score)/(Total # of bins) × 100.

#### 14. Single-cell gene expression co-variation analysis

We downloaded gene count tables of two independent batches (replicates) of Smart-seq2 on iPSC-derived motor neurons (94 and 87 in batch-1 and –2, respectively) from GSE138120 (https://www.ncbi.nlm.nih.gov/geo/query/acc.cgi?acc=GSE138120). We first removed low-quality cells (expressing fewer than 6,000 genes) and lowly expressed genes (expressed in fewer than 56 cells in batch-1 or less than 48 cells in batch-2). We calculated correlations between gene pairs using Spearman’s rank correlation coefficient (SCC) on the filtered count tables of batch-1 (73 cells × 5,291 genes) and batch-2 (69 cells × 5,379 genes). We calculated *P* values using a z-score for the Fisher transformation of the SCC. We applied further filtering steps as previously described^53^: i. we only considered gene pairs that correlated significantly (*P* < 0.01) in both replicates; ii. we discarded gene pairs if the sign of the SCC differed between replicates or if the sign was negative; iii. we discarded riboprotein gene pairs as these tend to dominate the positive covariations. Lastly, we used the mean SCC values of the gene pairs to calculate the Covariation Enrichment Score (CES), as previously described^53^. We separately calculated the CES for different gene sets, restricting our analysis to gene pairs separated by more than 5 Mb. Specifically, we analyzed:

1) Expressed protein-coding (pc) genes within genomic bins (100 kb) that are co-contacted by TIRs produced by 1, 2, 3, 4, 5, 6, or 7 out of the top-10 TIR source genes identified in NEU. We randomly selected each combination of TIR-producing genes for 100 times, each time computing the CES of the gene set selected.
2) TIR source genes, or pc genes located within 100 kb genomic bins classified as TIRCs or TIRC control regions, or gene pairs consisting of one TIR source gene and one expressed pc gene located within TIRCs. For each gene group, we randomly selected 300 genes for 20 times, each time calculating the CES of the selected gene set.

#### 15. GWAS trait-associated SNP enrichment analysis

To check whether disease risk loci are enriched inside TIR source and TIRC genes, we applied Stratified Linkage Disequilibrium Score Regression (S-LDSC)^55^ implemented in *ldsc* (v1.0.1) available at https://github.com/bulik/ldsc. Briefly, we calculated stratified linkage disequilibrium (LD) scores from the European ancestry samples in the 1000 Genomes Project Phase 3 data with a minor allele frequency higher than 0.05, including only HapMap3 SNPs. We then partitioned the SNP heritability of neurodevelopmental disorder traits (*n* = 18), neurodegenerative disorder traits (*n* = 7), immune related traits (*n* = 6) and other traits (*n* = 6) based on the annotations used in the Baseline Model v2.2^56^, including: i) DNaseI hypersensitivity sites; ii) promoter and enhancer sites; iii) protein-coding regions; iv) untranslated regions; and v) evolutionarily conserved regions (see **Supplementary Table 4**). In addition to these publicly available annotations, we also added TIR-related annotations, including: i) TIR source gene regions; ii) intronic regions of TIR source genes; iii) TIR control genes; iv) TIRCs; and v) TIRC control regions. We downloaded the 1000 Genomes Project Phase 3 data, the HapMap3 SNPs, the baseline model, and the GWAS summary statistics of 33 traits from https://alkesgroup.broadinstitute.org/LDSCORE/. We plotted the heritability enrichment (HE) of the selected annotations using R (v 4.0.5) and *ggplot2* (v 3.3.5) if the enrichment of the trait-annotation pairs was significant (*P* < 0.00001). The output of SNP heritability partitioning can be found in the Source Data (see Data Availability statement).

#### 16. De novo variant analysis

##### Data source

We utilized de novo variant (DNV) data from two autism cohorts supported by the Simons Foundation Autism Research Initiative (SFARI): the Simons Foundation Powering Autism Research for Knowledge (SPARK) and the Simons Simplex Collection (SSC). Both cohorts include individuals with autism and their families recruited in the United States, with detailed phenotypic and genomic data available through the SFARI Base (https://sfari.org/sfari-base). In the SPARK cohort, genome sequencing (GS) was performed on the Illumina NovaSeq 6000 platform using DNA extracted from saliva, followed by variant calling using GATK (v3.5) and GLnexus (v1.4.1). We downloaded SPARK GS data version 1.1 data and identified de novo variants using Slivar (v0.2.8) and GATK (v4.1.4.1), followed by stringent quality control filtering as previously described^81^. In SSC, GS was performed on the Illumina HiSeq X Ten platform using DNA extracted from blood, followed by variant calling using GATK (v4.1.0) and DeepVariant (v0.10). More details are available in ref. ^82^. We filtered out individuals with DNV counts beyond three times the standard deviation from the mean DNV count in each cohort separately. We included in our analysis 3,493 autistic and 2,188 non-autistic individuals in the SPARK cohort and 2,334 autistic and 1,843 non-autistic individuals in the SSC cohort.

##### Statistical analysis

To compare the de novo variant (DNV) burden between the introns of the top-55 TIR genes and control introns, we identified matched control introns from GENCODE (v38), selecting from genes expressed in NSC or NEU (transcripts per million >1, based on RNA-seq) and matching each TIR intron by length (± 10%) and GC-content (± 5%). We repeated this random matching process 100 times to generate reference intron sets. We excluded TIR introns without suitable matches based on expression criteria (10 for NSC and 4 for NEU) leaving 1,651 introns for further analysis. We then compared the DNV burden between TIR introns and matched control introns, separately for autistic and non-autistic individuals, using Generalized Estimating Equation (GEE) models, with DNV count modelled as a function of intron group (TIR vs. matched) and family ID included as a clustering variable to account for within-family correlation. For each cohort, we quantified the proportion of individuals harboring at least one DNV in each intron region, counting each individual only once per intron regardless of multiple DNVs. We evaluated the statistical significance of the difference of individual frequency between autism and non-autism using the Fisher’s exact test. We adjusted the *P* values for multiple hypothesis testing by applying the Benjamini-Hochberg method, separately for the SPARK and SSC cohorts.

## Code Availability

All the custom code used for processing and analyzing the data described in this study as well as to reproduce all the plots in the main and Supplementary Figures is available on GitHub at https://github.com/wenjingk/TIR_code. The code for DNV analysis is available at https://github.com/Tammimies-Lab/TIR_DNV.

