## Supplementary Information for "Chromatin-associated intronic RNAs from long genes form introsomes that shape nuclear architecture in neuronal cells"

1. Supplementary Figures pg. 2
2. Supplementary Tables pg. 62

### 1. Supplementary Figures

#### Supplementary Figure 1

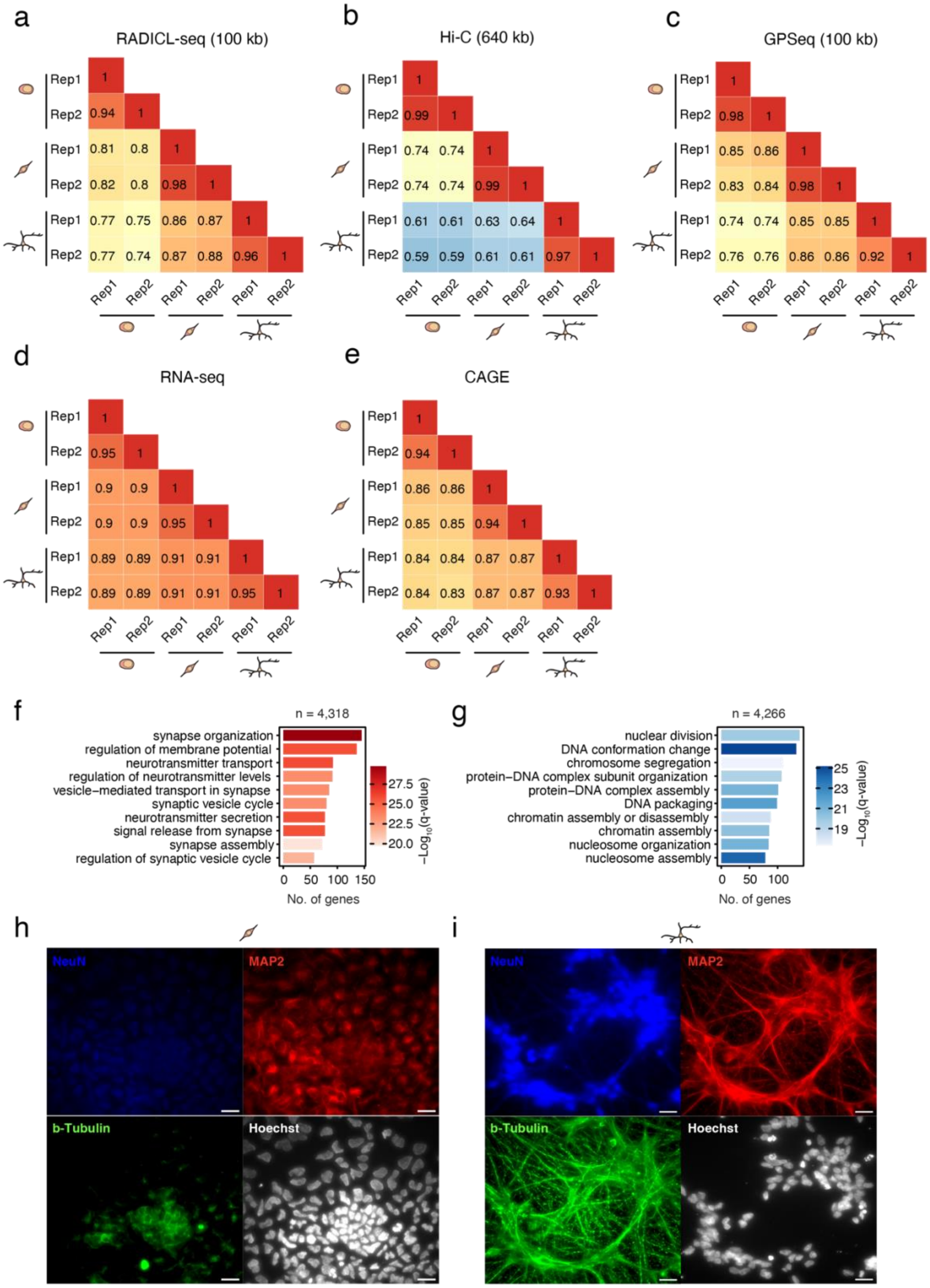

**Supplementary Figure 1. Validation of the assays and model system used in this study**. (**a-e**) Correlation (Spearman's correlation coefficient) between different biological replicates (Rep), for different assays, at the resolution indicated in brackets. For RNA-seq and CAGE, the correlation was calculated between the respective gene count tables. (**f**) Enriched gene ontology (GO) terms in biological processes for the genes upregulated (based on RNA-seq) in NEU compared to iPSC. *n*, number of upregulated genes. (**g**) As in (f) for genes downregulated in NEU compared to iPSC. (**h, i**) Maximum intensity projection of z-stack widefield microscopy images exemplifying the expression of various neuronal cell markers in NSC (h) and NEU (i), visualized by immunofluorescence. DNA was stained with Hoechst 33342. Scale bars, 20 μm. A link to the Source Data and code to regenerate the plots displayed in this figure is provided in the Data Availability and Code Availability statements.

#### Supplementary Figure 2

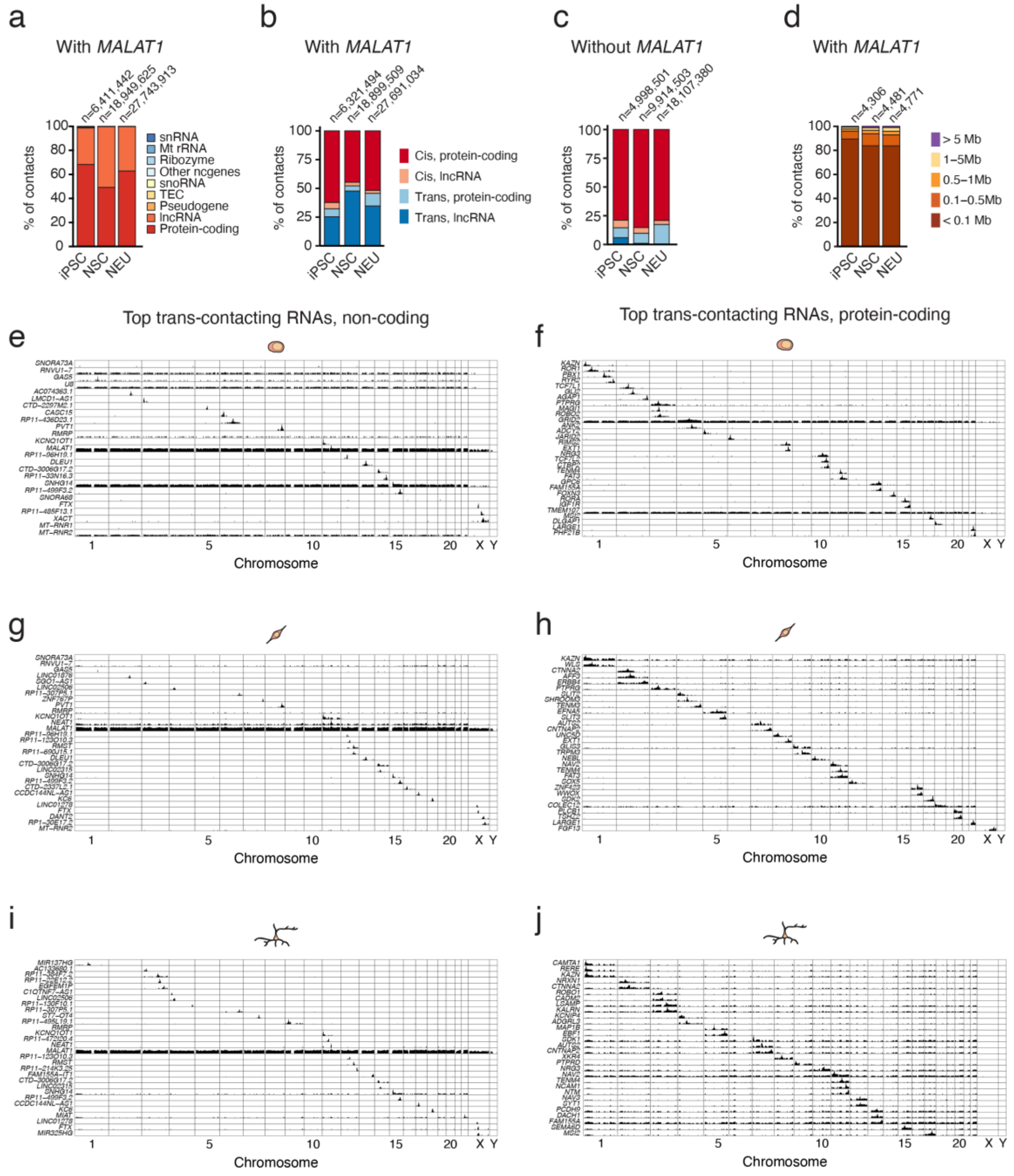


**Supplementary Figure 2. Genomic distribution of RNA-DNA contacts detected by RADICL-seq.** (**a**) Proportion of RNA-DNA contacts detected by RADICL-seq in each of the types/classes of RNA shown. *n*, number of RADICL-seq contacts, including *MALAT1* RNA contacts, detected in each cell type. (**b**) Proportion of RADICL-seq contacts representing contacts in cis or trans (the latter defined as contacts with DNA loci at a linear genomic distance ≥ 5 Mb from the source gene or on another chromosome). Only contacts of RNA transcribed from protein-coding and lncRNA genes (including *MALAT1*) are included. *n*, number of RNA-

DNA contacts detected in each cell type. (**c**) As in (b) excluding RNA-DNA contacts involving *MALAT1* lncRNA. (**d**) Proportion of RNA-DNA contacts mediated by lncRNAs (including *MALAT1*), for different genomic distances between the RNA source locus and the contacted loci. *n*, number of lncRNA genes producing RNA-DNA contacts detected by RADICL-seq in each cell type. (**e–j**) Genome-wide RADICL-seq contact profiles (100 kb resolution) for the top-30 trans-contacting RNAs derived from non-coding genes (e, g, i) and from protein-coding genes (f, h, j), in iPSC (e, f), NSC (g, h), and NEU (i, j). A link to the Source Data and code to regenerate the plots displayed in this figure is provided in the Data Availability and Code Availability statements.

#### Supplementary Figure 3

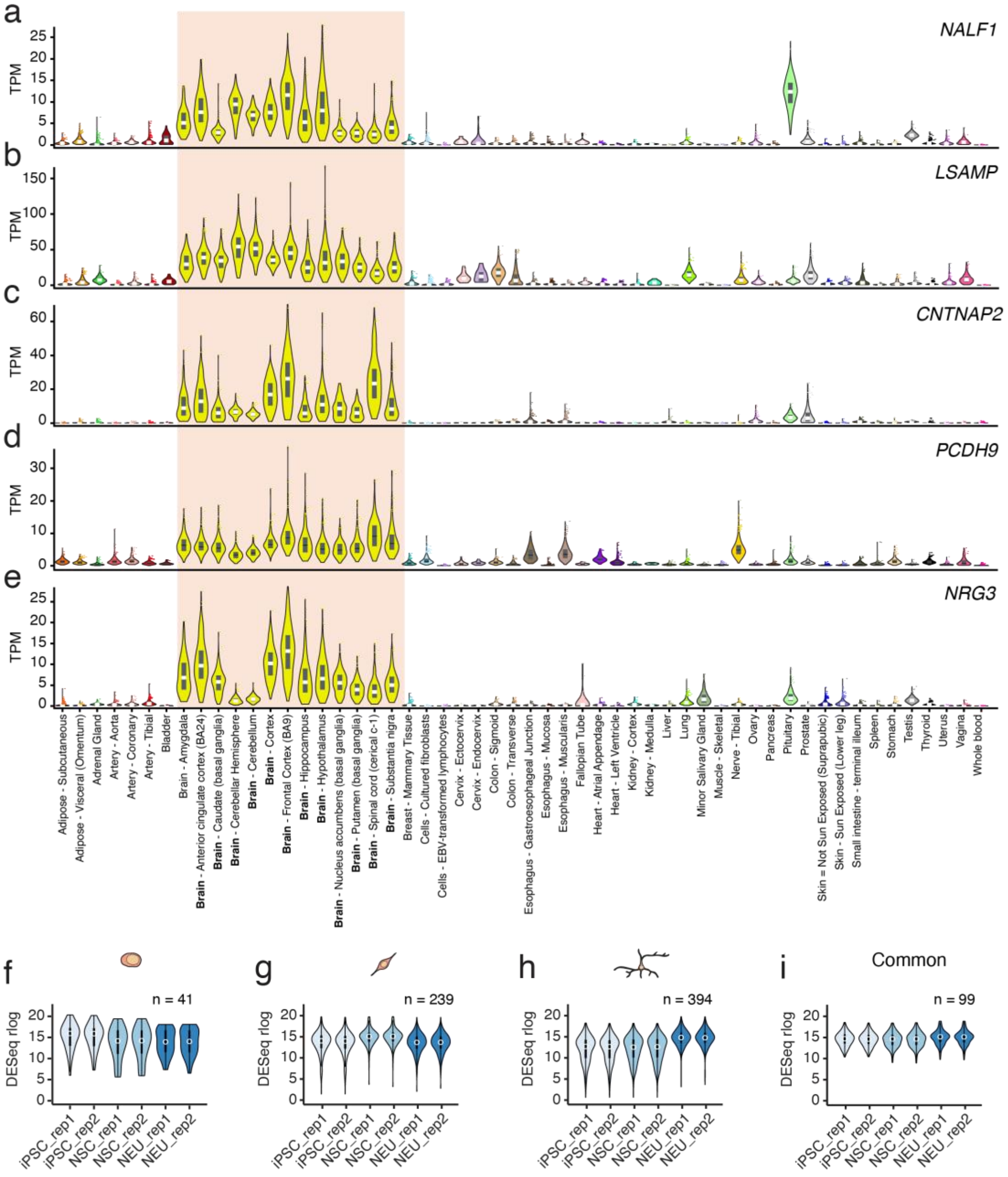


**Supplementary Figure 3. Source genes of trans-contacting RNAs show cell type-specific expression.** (**a-e**) Expression levels across different tissues of 5 genes selected from the top-10 neuronal trans-contacting RNA source genes, whose RNAs form trans contacts (> 5 Mb from the source locus) with the highest number of distinct 100 kb genomic bins. The peach rectangle highlights high levels of expression across different brain regions. The plots have been adapted from 'bulk tissue gene expression' plots downloaded from the GTEx Portal (release v10, RNA-seq data). TPM, transcripts per million. (**f-i**) Distribution of differential gene expression of cell type-specific (f-h) and common (i) trans-contacting RNA source genes calculated by DESeq for different cell types and biological replicates (rep). *n*, number of trans-contacting RNA source genes. Common trans-contacting RNAs in (i) were detected in all three cell types. In all the plots, violins extend from minimum to maximum, boxplots extend from the 25$^{th}$ to the 75$^{th}$ percentile, white dots represent the median, whiskers extend from –1.5×IQR to +1.5×IQR from the closest quartile. IQR, inter-quartile range. A link to the Source Data and code to regenerate the plots displayed in this figure is provided in the Data Availability and Code Availability statements.

Supplementary Figure 4

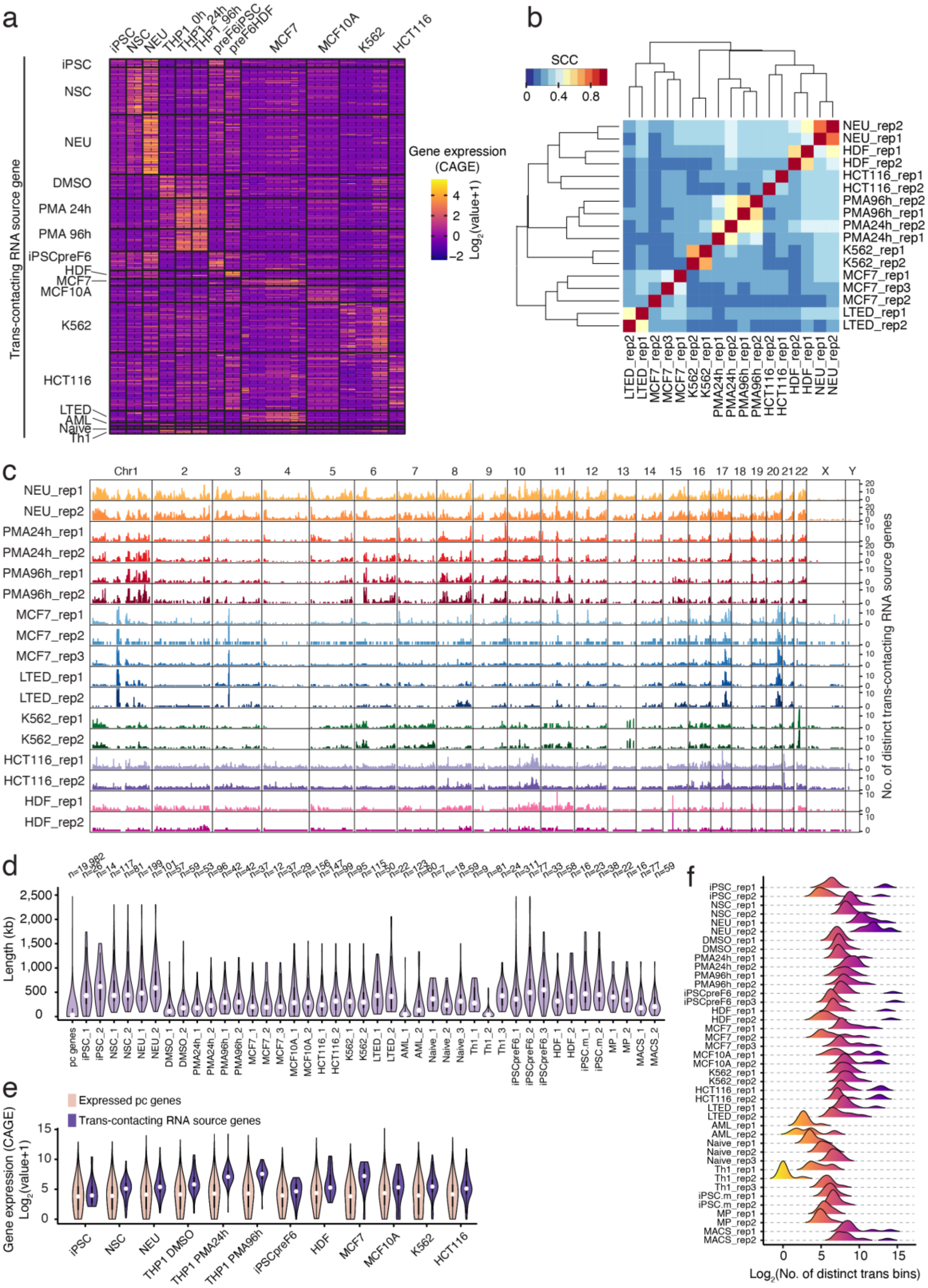


**Supplementary Figure 4. Trans-contacting RNA landscapes are cell type-specific. (a)** Gene expression measured by CAGE for trans-contacting RNA source genes, across all the cell types profiled by RADICL-seq in FANTOM6. Source genes were defined as protein-coding genes producing RNAs that contact more than 10 distinct genomic bins (100 kb resolution) in trans (≥ 5 Mb from the source locus) in each sample, based on RADICL-seq data. Vertical dashed black lines separate biological replicates of the same cell type. Each row represents one trans-contacting RNA source gene identified in the corresponding cell type. Only trans-contacting RNA source genes that were detected in all biological replicates are shown. Gene expression is scaled across samples. (**b**) Hierarchically clustered correlation heatmap showing pairwise correlations of trans contact counts between the indicated cell types and biological replicates (rep) profiled by RADICL-seq in FANTOM6. Note that the correlation is calculated based on the number of distinct trans-contacting RNA source genes. SCC, Spearman's correlation coefficient. (**c**) Genome-wide distribution (100 kb resolution) of RADICL-seq trans contacts across the indicated cell types and biological replicates (rep). (**d**) Distributions of the length of trans-contacting RNA source genes identified by RADICL-seq in the indicated cell types and biological replicates (rep). *n*, number of genes. (**e**) Distributions of gene expression levels (from CAGE) of all expressed protein-coding (pc) genes and trans-contacting RNA source genes identified by RADICL-seq in the indicated cell types and biological replicates (rep). (**f**) Distribution of the number of 100 kb genomic bins contacted in trans by RNAs from the top-10 trans-contacting RNA source genes identified by RADICL-seq in the indicated cell types and biological replicates (rep). Source genes were defined as genes producing trans-contacting RNAs forming contacts with the highest number of distinct 100 kb genomic bins. In (d) and (e), violins extend from minimum to maximum, boxplots extend from the $25^{th}$ to the $75^{th}$ percentile, white dots represent the median, whiskers extend from –1.5×IQR to +1.5×IQR from the closest quartile. IQR, inter-quartile range. A link to the Source Data and code to regenerate the plots displayed in this figure is provided in the Data Availability and Code Availability statements.

Supplementary Figure 5

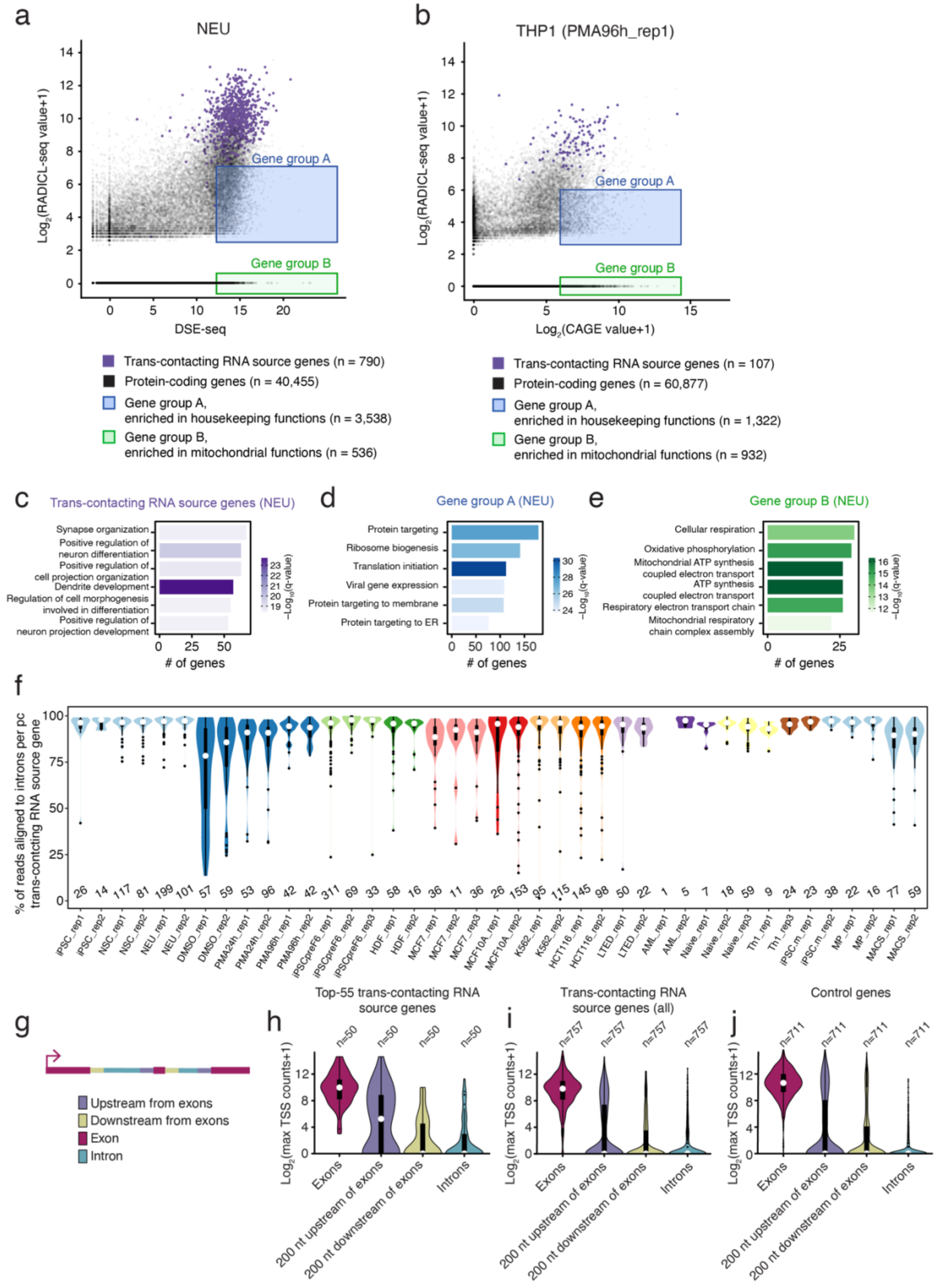


**Supplementary Figure 5. Trans-contacting RNAs mainly derive from introns of moderate-to-highly expressed cell type specific genes.** (**a**) Scatterplot showing the relationship between RNA-DNA contact frequencies measured by RADICL-seq and gene expression levels measured by RNA-seq, for protein-coding genes expressed in neurons (NEU). Each dot represents a gene. Purple dots: trans-contacting RNA source genes defined as genes whose RNAs contact more than one distinct 100 kb genomic bin in trans, in NEU. Gene group A: highly expressed genes with relatively few RNA-DNA contacts. Gene group B: highly expressed genes not producing trans-contacting RNAs. *n*, number of genes. (**b**) As in (a) for the indicated replicate (rep) of THP1-derived macrophages treated with Phorbol myristate acetate (PMA) for 96 hours (see **Supplementary Table 1**) and gene expression measured by CAGE. (**c-e**) Gene Ontology terms enriched in each of the three gene classes shown in violet, blue, and green in (a). (**f**) Distributions of the proportion of intron-derived reads for each trans-contacting RNA source gene in the indicated cell types and biological replicates (rep), as detected by RADICL-seq. Source genes were defined as protein-coding genes whose RNAs contact more than 10 distinct 100 kb genomic bins in trans (≥5 Mb from the source locus), based on RADICL-seq. The numbers below each violin plot indicate the number of source genes analyzed in each group. (**g**) Scheme of different regions along the gene body used for the analysis of CAGE data in (h-j). (**h-j**) Distributions of promoter-like transcription start site (TSS) peaks from CAGE, across different parts of the gene body schematically shown in (g), for the top-55 trans-contacting RNA source genes identified in NEU (h), all trans-contacting RNA source genes in NEU (i), and control genes (j). *n*, number of genes. A link to the Source Data and code to regenerate the plots displayed in this figure is provided in the Data Availability and Code Availability statements.

Supplementary Figure 6

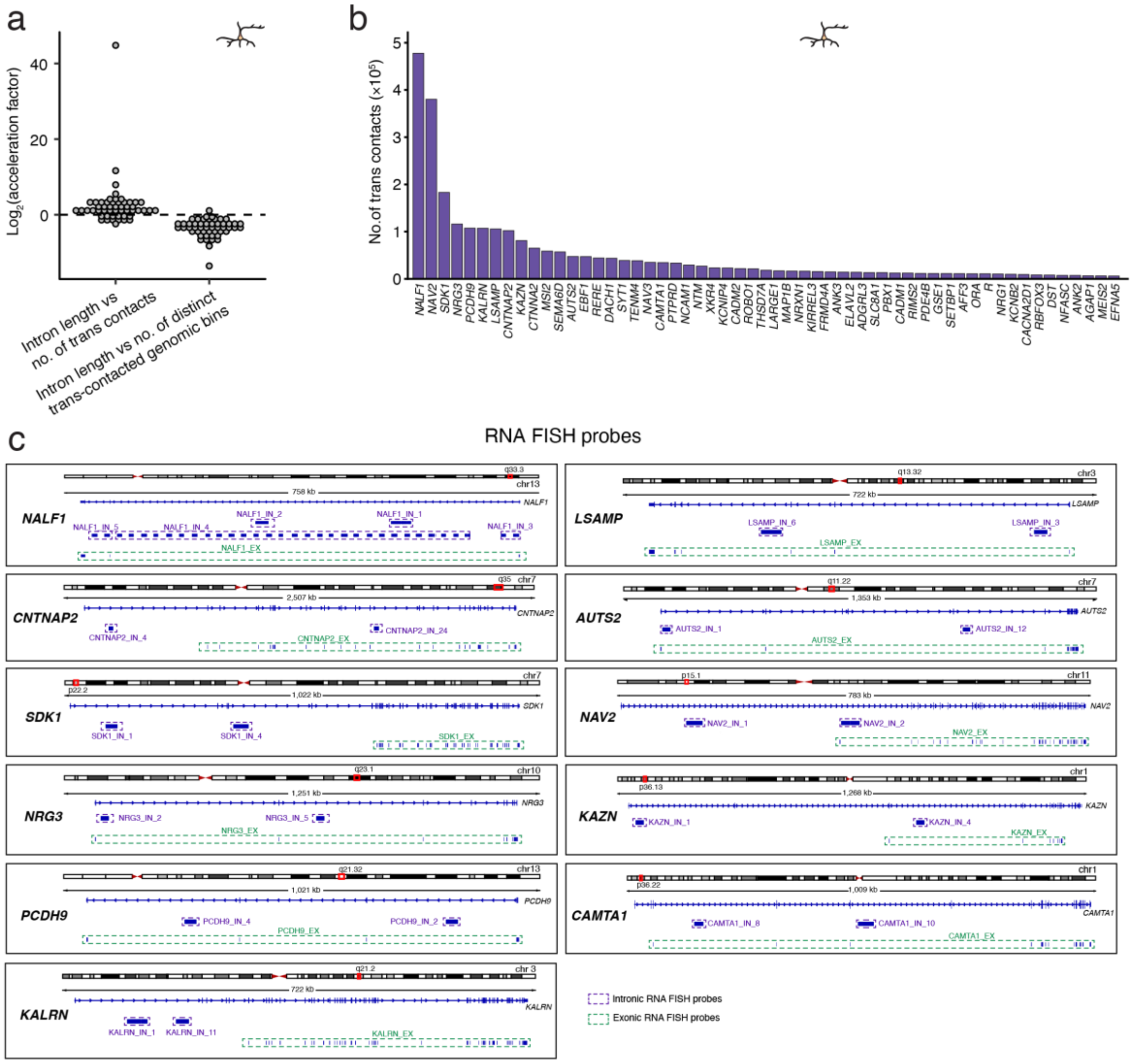


**Supplementary Figure 6. Location of the RNA FISH probes used to visualize the top TIRs identified in NEU.** (**a**) Distribution of the acceleration factor calculated for the best-fitting model of the relationship between intron length versus number of trans contacts or number of trans-contacted genomic bins (100 kb resolution), for the top-55 TIRs identified by RADICL-seq in NEU (see **Fig. 2a, b** for examples of scatterplots showing these relations and the corresponding fitted curves). Each dot corresponds to one gene. Linear, saturating, exponential, and power-law growth models were used for fitting. The acceleration factor quantifies the change in growth rate across different intron lengths in the best-fitting model: values >1 indicate accelerating growth, values ~1 indicate constant growth, and values <1 indicate saturating behavior. (**b**) Ranked number of RADICL-seq trans contacts for the top-55 TIRs identified in NEU. (**c**) Scheme of the genomic location and span of the RNA FISH probes used to visualize the indicated top TIR source genes identified in NEU. Dashed purple boxes: probes targeting the indicated intronic (IN) regions. Dashed green boxes: probes targeting the indicated exons (EX). In each dashed box, the vertical blue bars indicate individual 40-nt oligos in the corresponding probe. The nomenclature of the probes is the same as in **Supplementary Table 3**. Note: for intronic (IN) probes, the number in the probe name represents the probe number and not the number of the corresponding intron. The red boxes on top of each chromosome ideogram correspond to the gene locus shown below (blue track).

Supplementary Figure 7

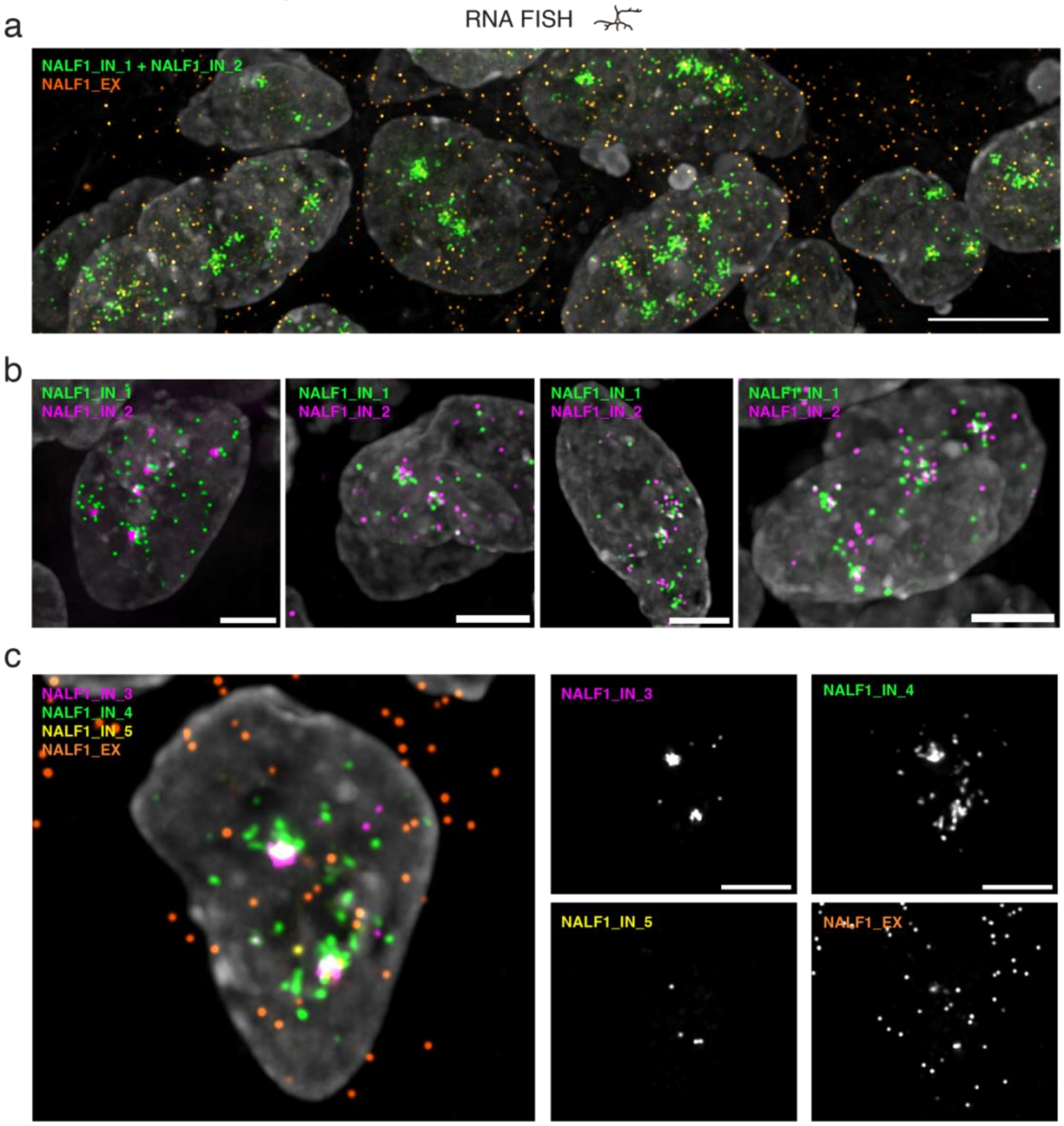


**Supplementary Figure 7.** ***NALF1*** **intronic RNA forms distinct patterns in the nucleus of neurons.** (**a**) Maximum intensity projection of a z-stack widefield microscopy image exemplifying the nuclear pattern of TIRs originating from *NALF1* introns (green) together with *NALF1* exonic RNA (orange) in NEU cells, visualized using the indicated RNA FISH probes. Gray, DNA stained by Hoechst 33342. Scale bar, 5 µm. (**b**) As in (a) using different *NALF1* intronic probes. (**c**) As in (a) using a different set of *NALF1* intronic and exonic probes. See **Supplementary Table 3** for the list of oligos composing the FISH probes shown in this figure. All the images in the figure were deconvolved using Deconwolf (https://deconwolf.fht.org/). A link to the Source Data and code to regenerate the plots displayed in this figure is provided in the Data Availability and Code Availability statements.

Supplementary Figure 8

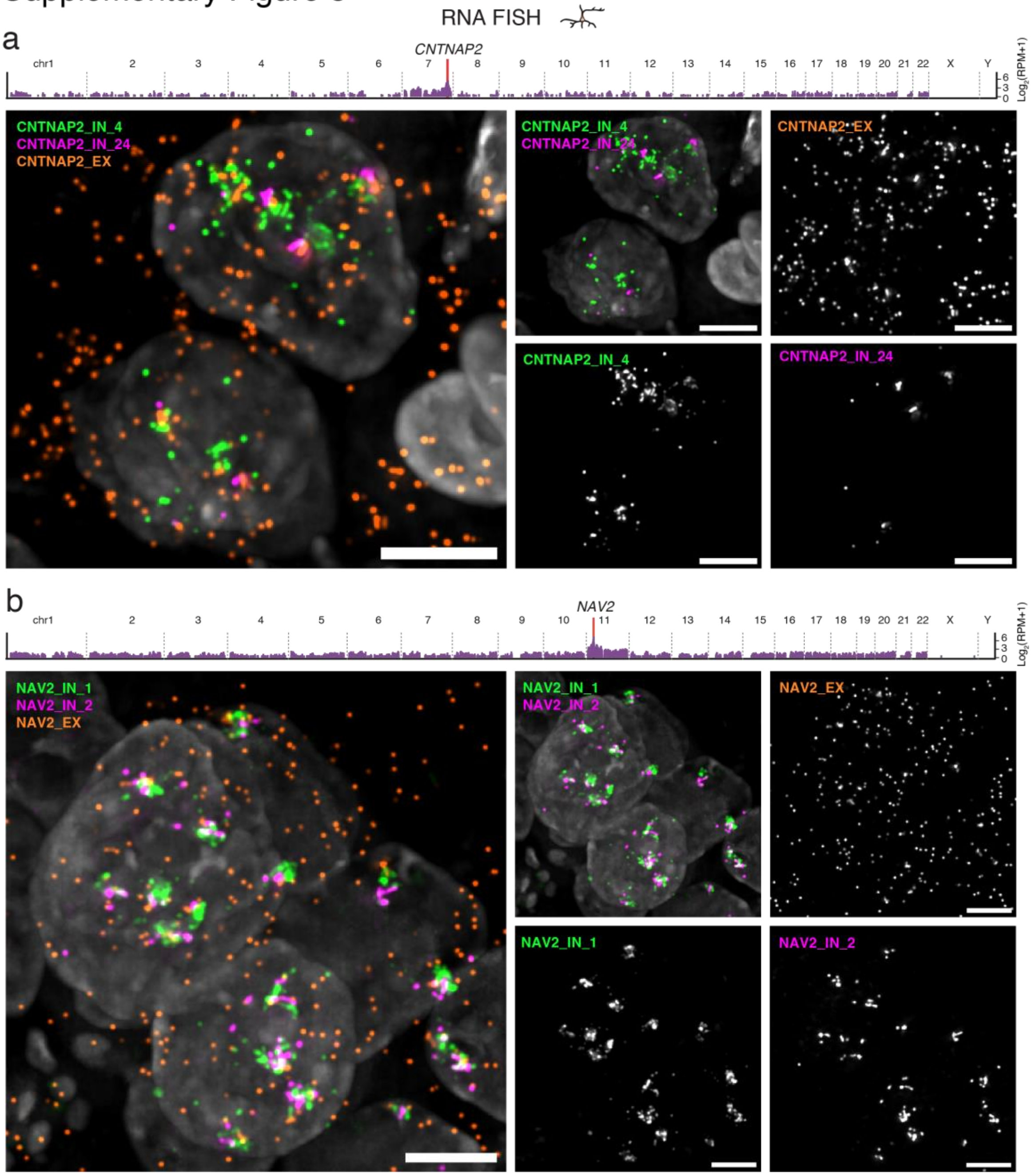


**Supplementary Figure 8. TIRs form cloud-like clusters in the nucleus of neuronal cells.** (**a**) Maximum intensity projection of a z-stack widefield microscopy image exemplifying the nuclear pattern of TIRs (green and magenta) and of exonic RNAs (orange) originating from *CNTNAP2* in NEU cells, visualized by RNA FISH. Gray, DNA stained by Hoechst 33342. Scale bar, 5 µm. The top bar plot displays the genome-wide distribution of trans contacts (100 kb resolution) made by RNAs derived from the indicated source gene (red bar), as detected by RADICL-seq. RPM, reads per million. (**b**) As in (a) using RNA FISH probes targeting intronic and exonic regions of the TIR source gene *NAV2*. See **Supplementary Table 3** for the list of oligos composing the FISH probes shown in this figure. All the images in the figure were deconvolved using Deconwolf (https://deconwolf.fht.org/). A link to the Source Data and code to regenerate the plots displayed in this figure is provided in the Data Availability and Code Availability statements.

Supplementary Figure 9

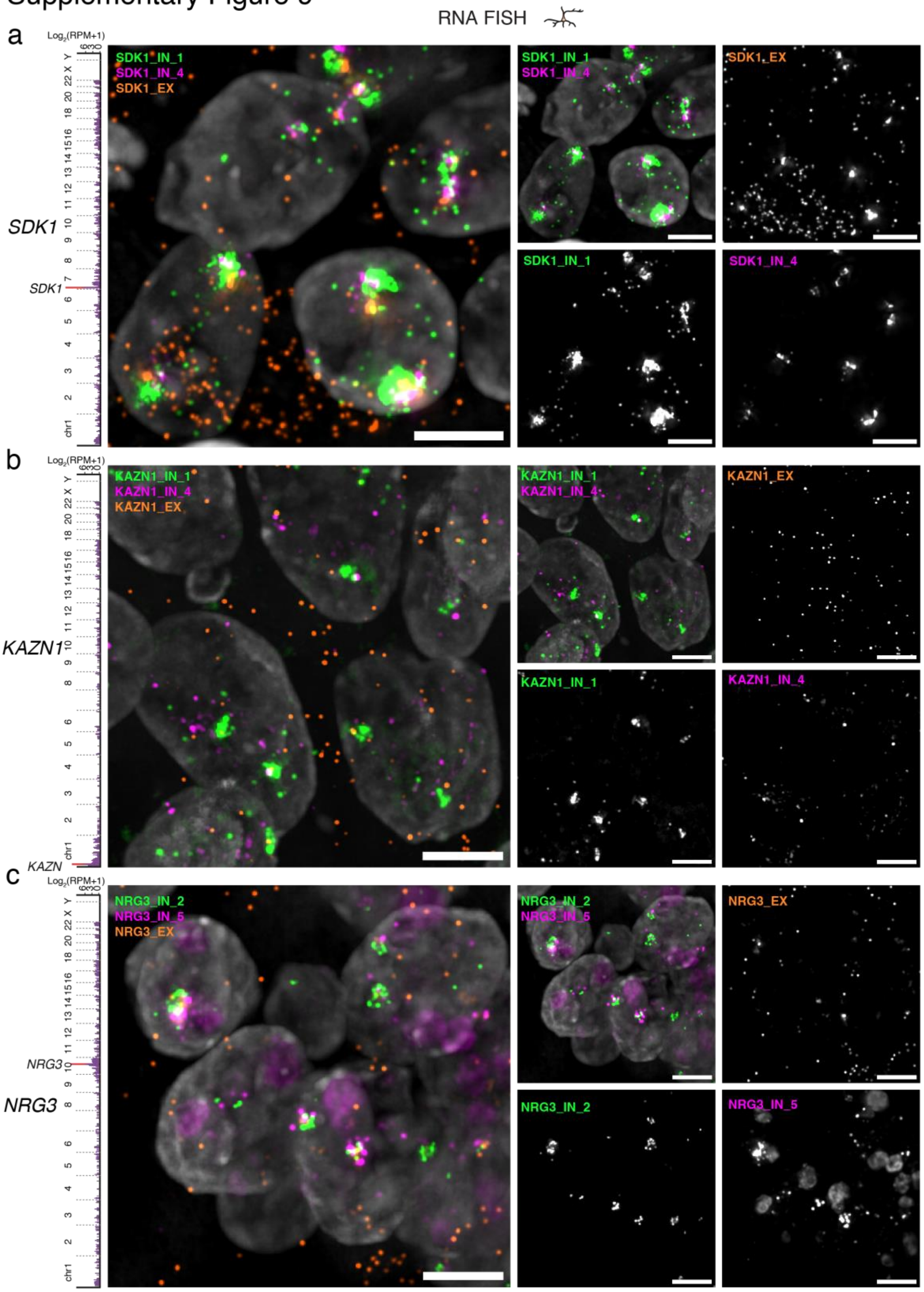


**Supplementary Figure 9. TIRs form cloud-like clusters in the nucleus of neuronal cells.** (**a**) Maximum intensity projection of a z-stack widefield microscopy image exemplifying the nuclear pattern of TIRs (green) and exonic RNAs originating from *SDK1* in NEU cells, visualized by RNA FISH. Gray, DNA stained by Hoechst 33342. Scale bar, 5 µm. The left bar plot displays the genome-wide distribution of trans contacts (100 kb resolution) made by RNAs derived from the indicated source gene (red bar), as detected by RADICL-seq. RPM, reads per million. (**b-c**) As in (a) using RNA FISH probes targeting intronic and exonic regions of the TIR source genes *KAZN* (b) and *NRG3* (c). See **Supplementary Table 3** for the list of oligos composing the FISH probes shown in this figure. All the images in the figure were deconvolved using Deconwolf (https://deconwolf.fht.org/). A link to the Source Data and code to regenerate the plots displayed in this figure is provided in the Data Availability and Code Availability statements.

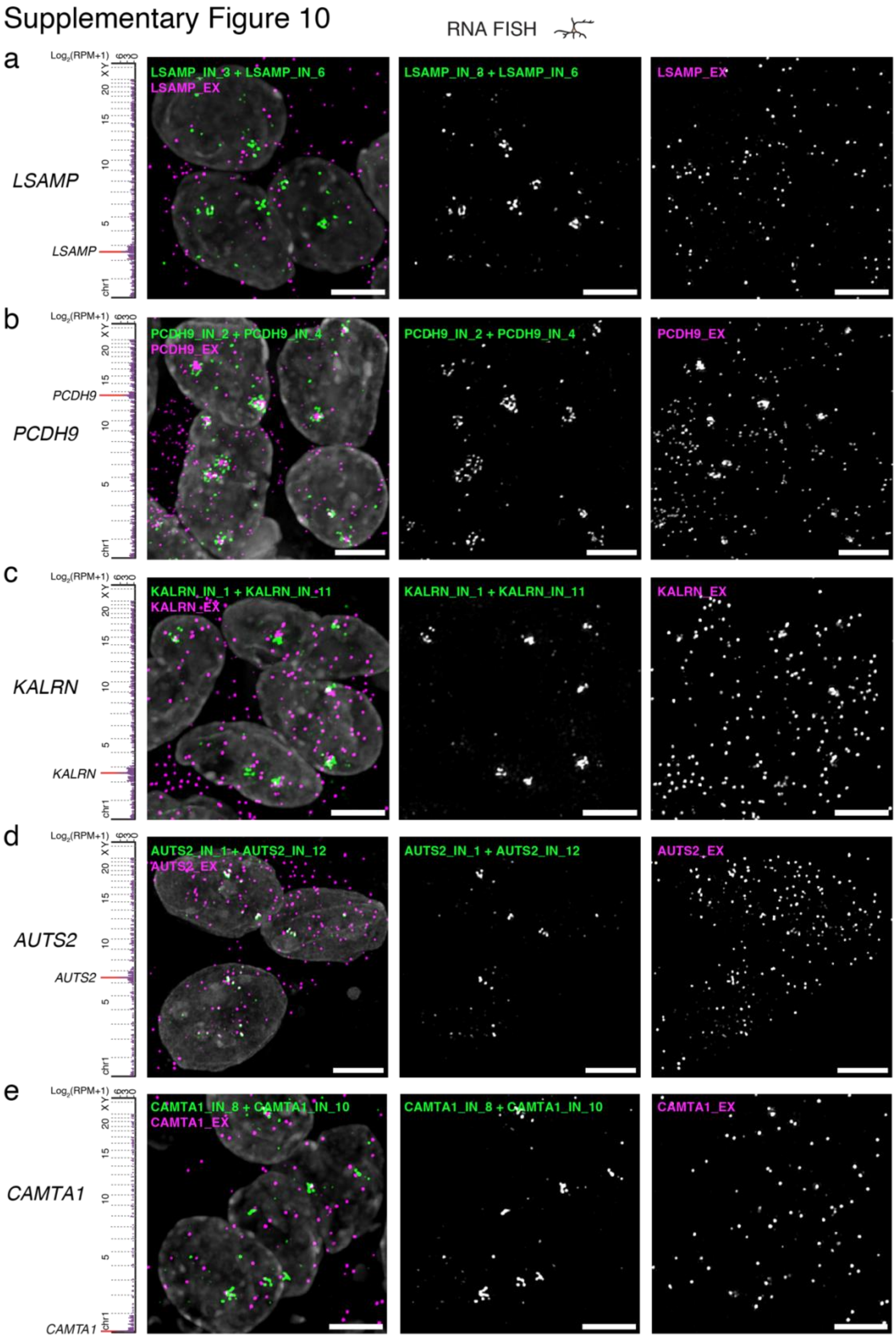


**Supplementary Figure 10. TIRs form cloud-like clusters in the nucleus of neuronal cells.** (**a**) Maximum intensity projection of a z-stack widefield microscopy image exemplifying the nuclear pattern of TIRs (green) and exonic RNAs originating from *LSAMP* in NEU cells, visualized by RNA FISH. Gray, DNA stained by Hoechst 33342. Scale bar, 5 µm. The left bar plot displays the genome-wide distribution of trans contacts (100 kb resolution) made by RNAs derived from the indicated source gene (red bar), as detected by RADICL-seq. RPM, reads per million. (**b-e**) As in (a) using RNA FISH probes targeting intronic and exonic regions of the TIR source genes *PCDH9* (b), *KARLN* (c), *AUTS2* (d) and *CAMTA1* (e). In all the images, the magenta dots represent the exons of the corresponding gene. See **Supplementary Table 3** for the list of oligos composing the FISH probes shown in this figure. All the images in the figure were deconvolved using Deconwolf (https://deconwolf.fht.org/). A link to the Source Data and code to regenerate the plots displayed in this figure is provided in the Data Availability and Code Availability statements.

Supplementary Figure 11

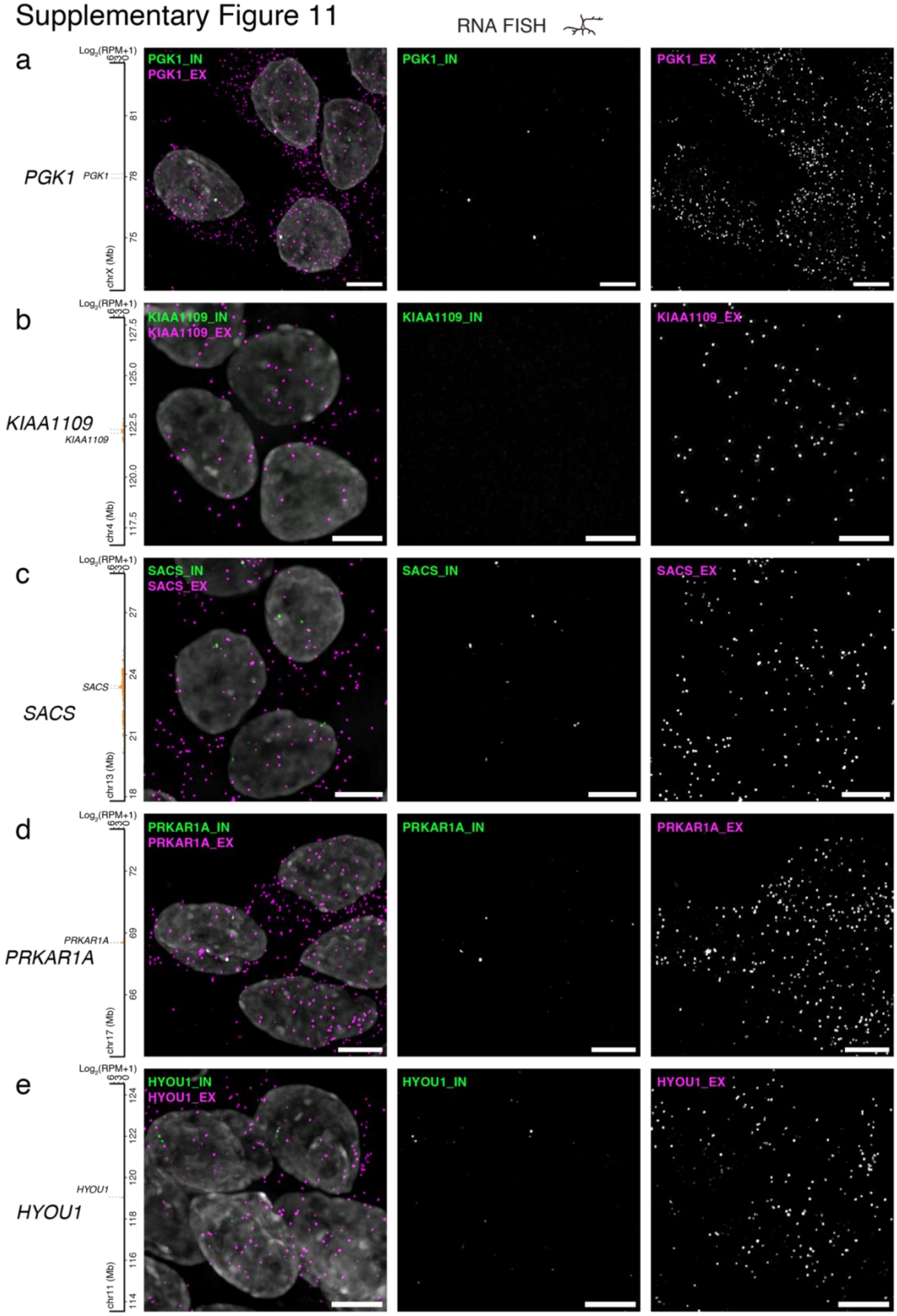


**Supplementary Figure 11. TIR control genes produce intronic RNAs that do not accumulate in the nucleus of neurons.** (**a**) Maximum intensity projection of a z-stack widefield microscopy image exemplifying the nuclear pattern of intronic (green) and exonic RNAs (magenta) originating from the TIR control gene *PGK1* in NEU cells, visualized by RNA FISH. Gray, DNA stained by Hoechst 33342. Scale bars, 5 µm. The left bar plot displays the distribution along the indicated chromosome (chr) of RNA-DNA contacts (100 kb resolution) made by RNAs derived from the indicated TIR control gene (dashed bar), as detected by RADICL-seq. RPM, reads per million. (**b-e**) As in (a) using RNA FISH probes targeting the introns (green) and exons (magenta) of the TIR control genes *KIAA1109* (b), *SACS* (c), *PRKAR1A* (d), and *HYOU1* (e). See **Supplementary Table 3** for the list of oligos composing the FISH probes shown in this figure. All the images in the figure were deconvolved using Deconwolf (https://deconwolf.fht.org/). A link to the Source Data and code to regenerate the plots displayed in this figure is provided in the Data Availability and Code Availability statements.

### Supplementary Figure 12

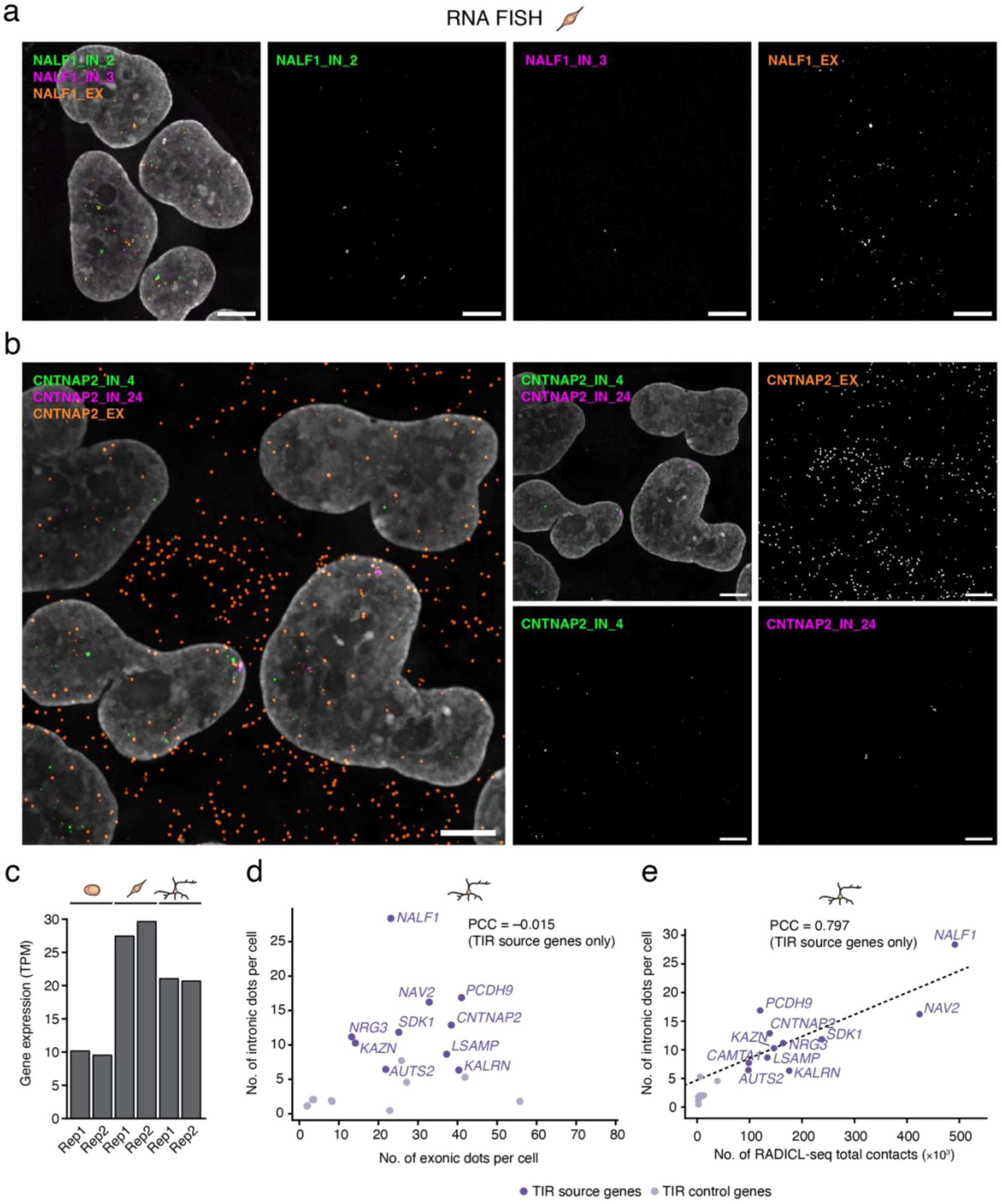


**Supplementary Figure 12. Neuron-specific TIRs do not accumulate in the nucleus of NSC.** (**a**) Maximum intensity projection of a z-stack widefield microscopy image exemplifying the nuclear pattern of intronic and exonic RNAs originating from the TIR source gene *NALF1* in NSC cells, visualized by RNA FISH. Gray, DNA stained by Hoechst 33342. Scale bar: 5 µm. (**b**) As in (a) using RNA FISH probes targeting the intron and exons of the TIR source gene

*CNTNAP2*. (**c**) *CNTNAP2* gene expression in the indicated cell type and biological replicate (Rep), as measured by RNA-seq. TPM, transcripts per million. (**d**) Quantification of the number of intronic and exonic RNA FISH dots per cell, for the TIR source and TIR control genes shown in **Supplementary Fig. 8-11**, in NEU cells. PCC, Pearson's correlation coefficient. (**e**) Correlation between the number of discrete intronic RNA FISH signals (dots) per cell, in NEU cells, and the total number of DNA contacts formed by the RNA produced from the corresponding gene as measured by RADICL-seq, for different TIR source and TIR control genes, in NEU cells. Dashed black line: linear regression. In (d) and (e), the FISH dot counts represent averages calculated from 335–943 cells across 8–12 fields of view per condition, and from 1–2 biological replicates for each gene. See **Supplementary Table 3** for the list of oligos composing the FISH probes shown in this figure. All the images in the figure were deconvolved using Deconwolf (https://deconwolf.fht.org/). A link to the Source Data and code to regenerate the plots displayed in this figure is provided in the Data Availability and Code Availability statements.

#### Supplementary Figure 13

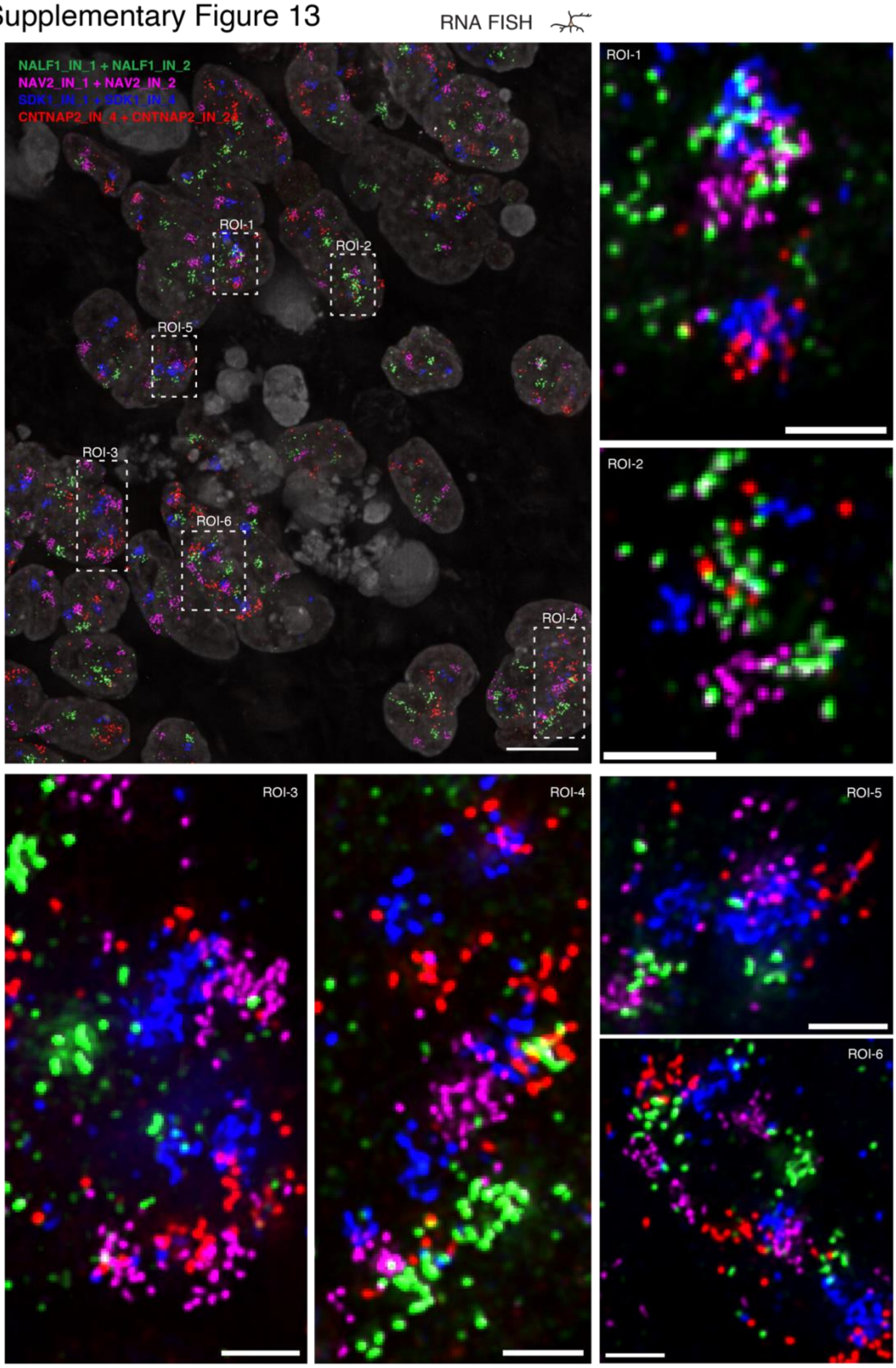


**Supplementary Figure 13. TIR clouds show highly variable localization patterns.** Large top-left image: maximum intensity projection of a z-stack widefield microscopy image exemplifying the nuclear pattern of TIRs produced from the *NALF1*, *NAV2*, *SDK1*, and *CNTNAP2* source genes in NEU cells, visualized by RNA FISH. Gray, DNA stained by Hoechst 33342. The regions of interest (ROI) marked by dashed rectangles are magnified in the corresponding images on the right and bottom. Scale bars: 10 µm (large image) and 2 µm (ROI zoom-ins). See **Supplementary Table 3** for the list of oligos composing the FISH probes shown in this figure. All the images in the figure were deconvolved using Deconwolf (https://deconwolf.fht.org/). A link to the Source Data and code to regenerate the plots displayed in this figure is provided in the Data Availability and Code Availability statements.

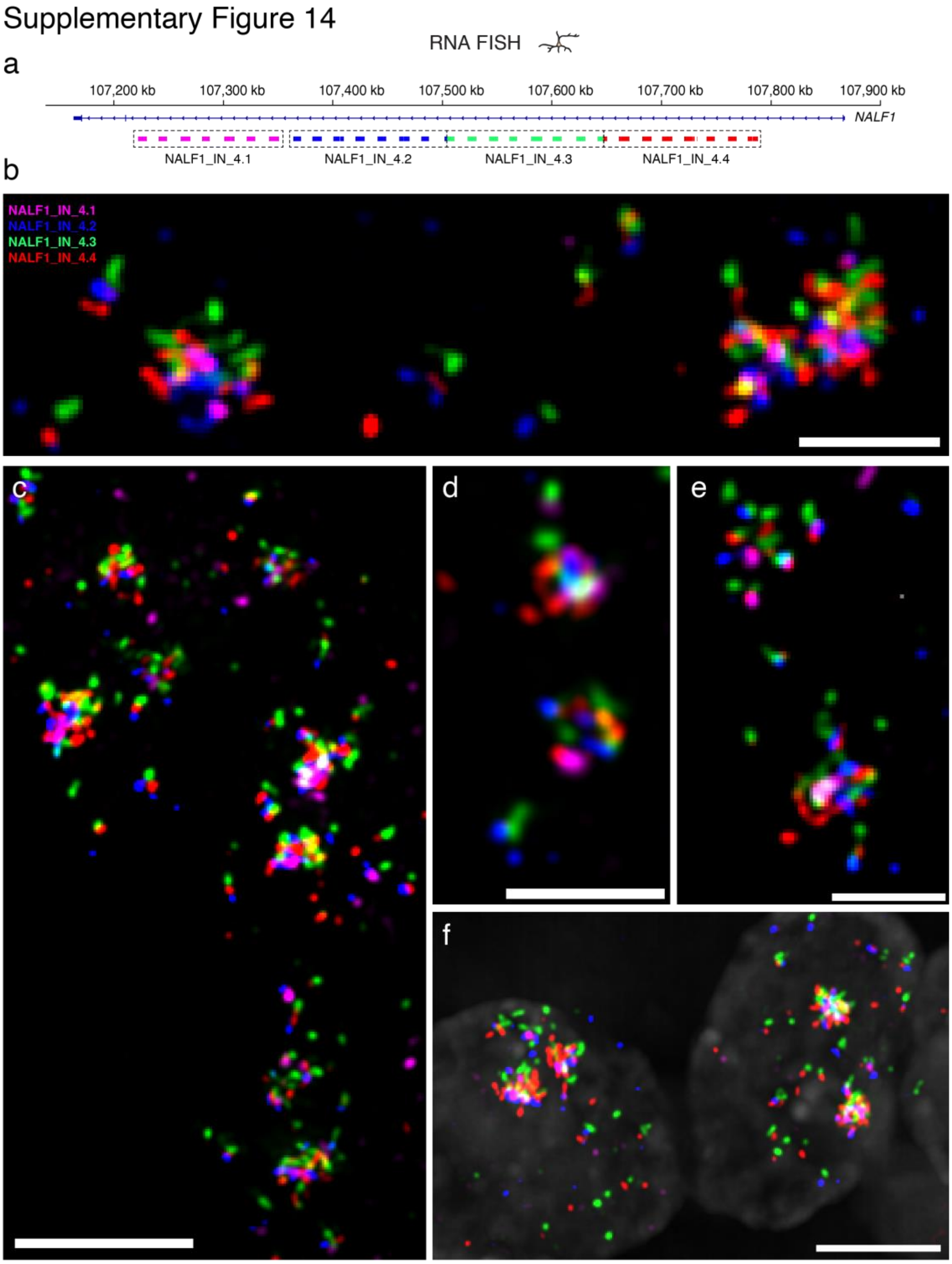


**Supplementary Figure 14. *NALF1* TIRs are exceptionally long RNA molecules. (a)** Scheme of the genomic location and span of the RNA FISH probes used to visualize a more than 600 kb-long intronic region in the *NALF1* TIR source gene. In each dashed box, the vertical bars indicate individual oligos in the corresponding probe. Each probe contains a subset of the oligos in the NALF1_IN_4 probe shown in **Supplementary Fig. 6c**. Note: the number in each probe name represents the probe number and not the number of the corresponding intron. (**b-f**) Maximum intensity projections of z-stack widefield microscopy images exemplifying the nuclear pattern of *NALF1* TIRs in NEU cells, visualized using the RNA FISH probes shown in (a). Gray, DNA stained by Hoechst 33342. Scale bars: 5 µm (c, f) and 2 µm (b, d, e). See **Supplementary Table 3** for the list of oligos composing the FISH probes shown in this figure. All the images in the figure were deconvolved using Deconwolf (https://deconwolf.fht.org/). A link to the Source Data and code to regenerate the plots displayed in this figure is provided in the Data Availability and Code Availability statements.

Supplementary Figure 15

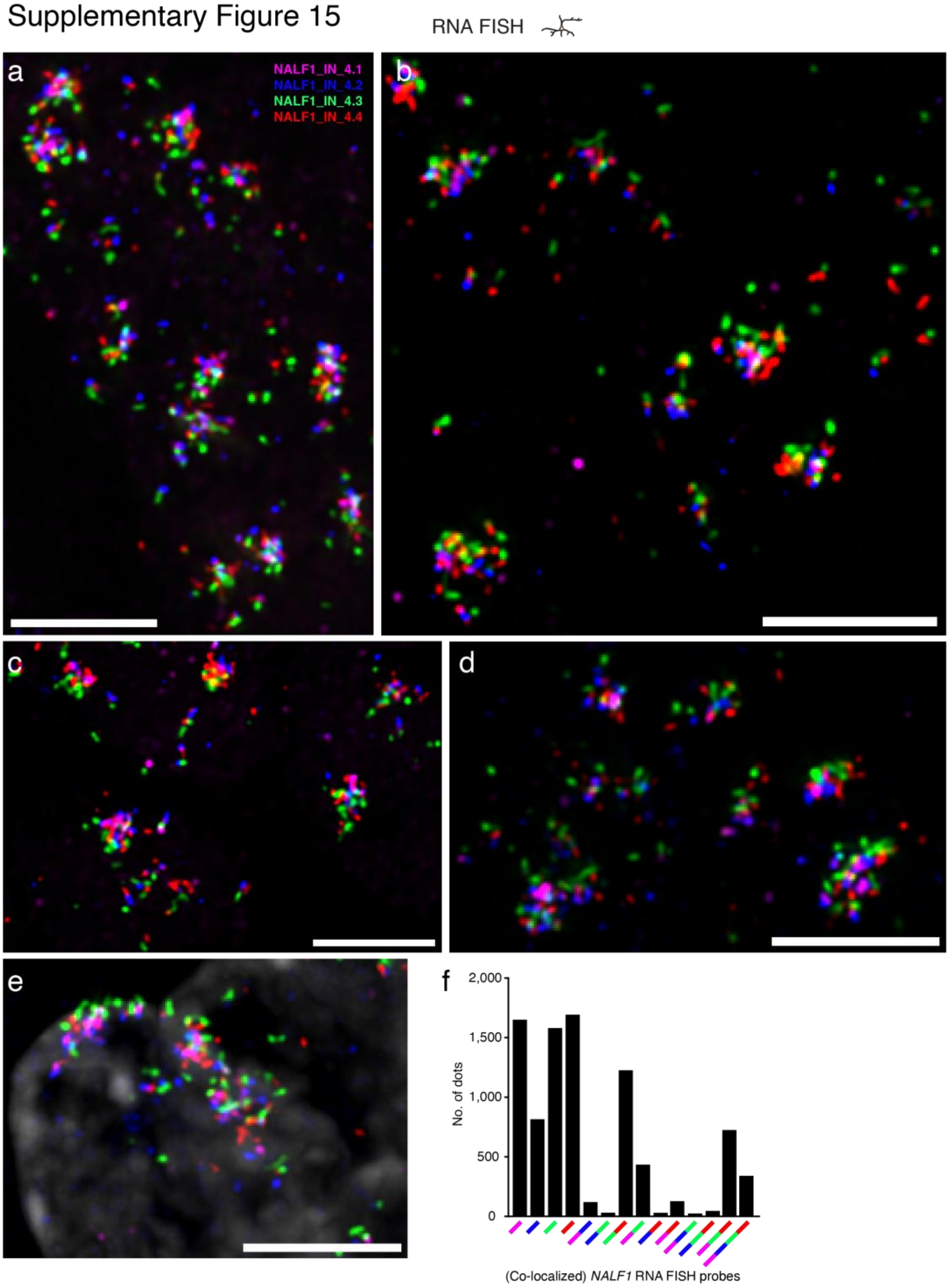


**Supplementary Figure 15. *NALF1* TIRs are exceptionally long RNA molecules. (a-e)** Maximum intensity projections of z-stack widefield microscopy images exemplifying the nuclear pattern of *NALF1* TIRs in NEU cells, visualized using the RNA FISH probes shown in **Supplementary Fig. 14a**. Gray, DNA stained by Hoechst 33342. Scale bars: 5 µm. **(f)** Quantification of the number of individual or co-localized RNA FISH dots detected in the dataset of which the image shown in (a-e) and the images shown in **Supplementary Fig. 14b-f** are representative examples. The colored segments below each bar indicate the type of co-localization event detected. The color of each segment is the same as the color of the corresponding RNA FISH probe shown in **Supplementary Fig. 14a**. See **Supplementary Table 3** for the list of oligos composing the FISH probes shown in this figure. All the images in the figure were deconvolved using Deconwolf (https://deconwolf.fht.org/). A link to the Source Data and code to regenerate the plots displayed in this figure is provided in the Data Availability and Code Availability statements.

Supplementary Figure 16

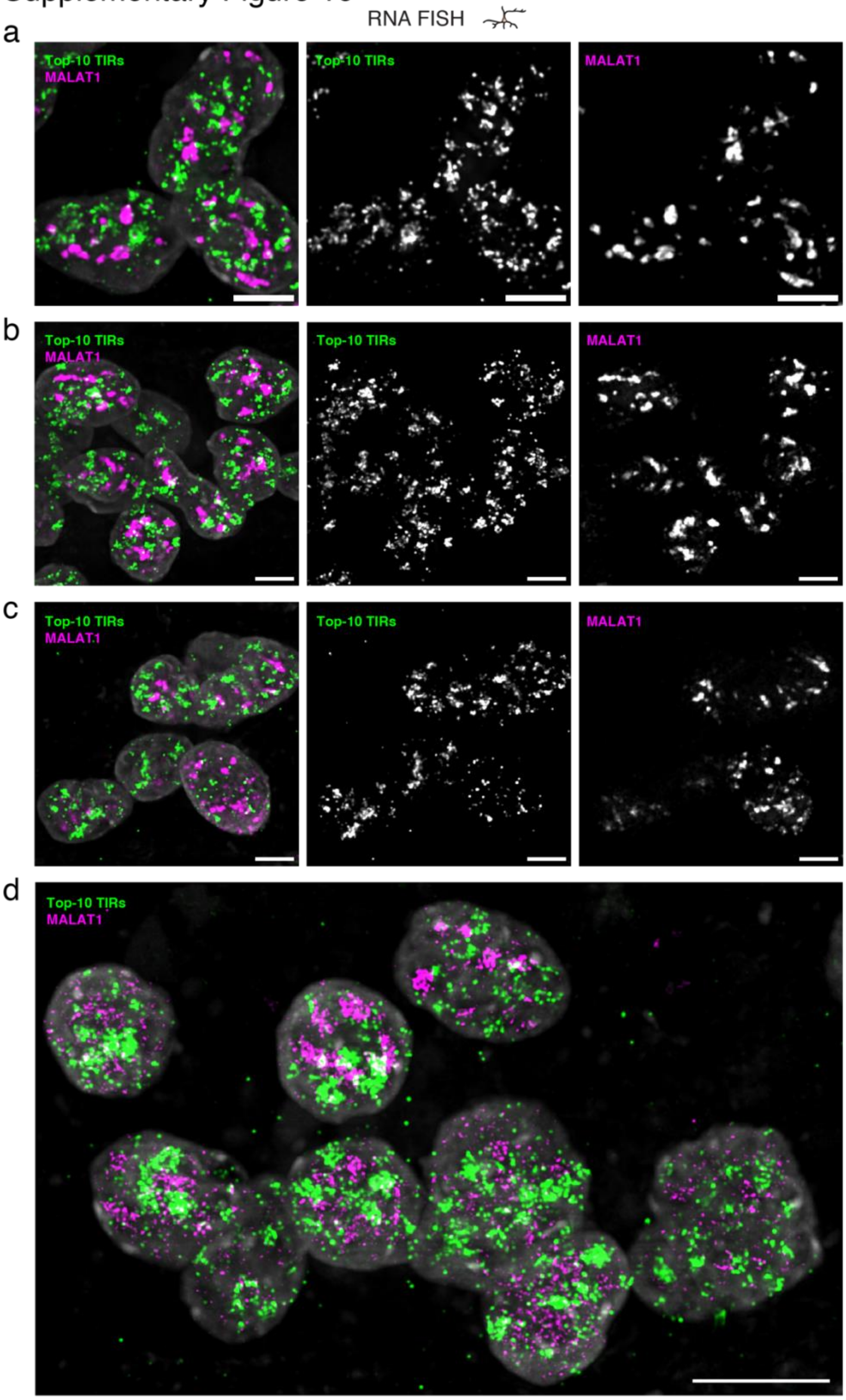


**Supplementary Figure 16. TIR clouds are spatially separated from speckles. (a-d)** Maximum intensity projections of z-stack widefield microscopy images of NEU cells highlighting the absence of co-localization between *MALAT1* RNA (magenta, marking speckles) and intronic RNAs originating from the top-10 TIR source genes (green), visualized by RNA FISH in NEU cells. All the intronic (IN) RNA FISH probes depicted in **Supplementary Fig. 6c** (except probes targeting *CAMTA1*) were pooled together and labeled with the same color (green). Gray, DNA stained by Hoechst 33342. Scale bars: 5 µm. See **Supplementary Table 3** for the list of oligos composing the FISH probes shown in this figure. All the images in the figure were deconvolved using Deconwolf (https://deconwolf.fht.org/). A link to the Source Data and code to regenerate the plots displayed in this figure is provided in the Data Availability and Code Availability statements.

Supplementary Figure 17

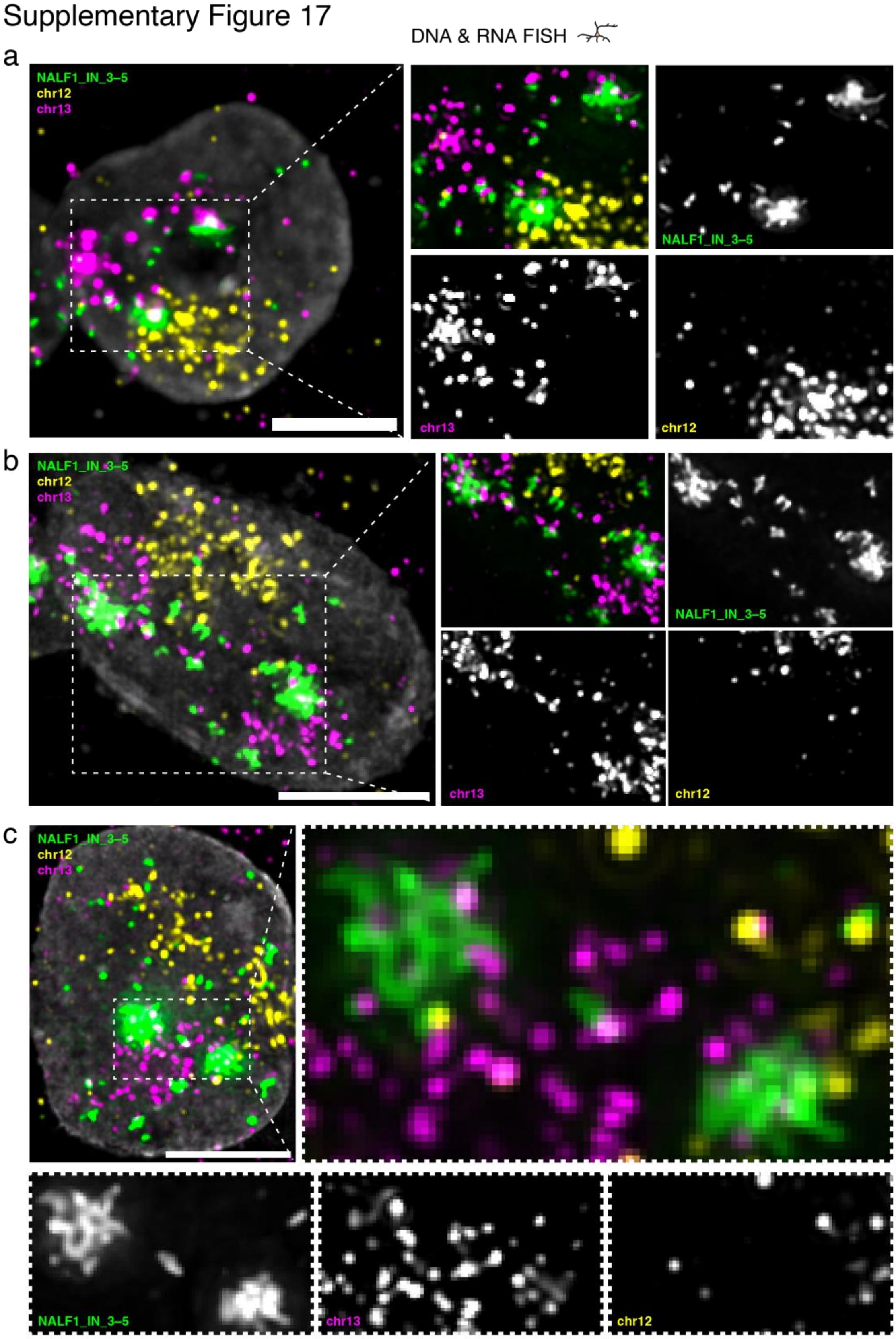


**Supplementary Figure 17. TIRs clouds extend beyond the chromosomal territory of their source gene.** (**a-c**) Maximum intensity projections of z-stack widefield microscopy images exemplifying the reciprocal positioning of *NALF1* TIRs (green) together with chromosome (chr) 13 (magenta), which harbors the *NALF1* gene, and chr12 (yellow), which engages in extensive trans contacts with the top-55 TIR source genes but does not contain any of the top-10 TIR source gene, visualized by combined DNA and RNA FISH in NEU cells. Gray, DNA stained by Hoechst 33342. Scale bars: 5 µm. The regions marked by dashed rectangles are magnified in the corresponding images on the right (a-c) and bottom (c). See **Supplementary Table 3** for the list of oligos composing the FISH probes shown in this figure. All the images in the figure were deconvolved using Deconwolf (https://deconwolf.fht.org/). A link to the Source Data and code to regenerate the plots displayed in this figure is provided in the Data Availability and Code Availability statements.

Supplementary Figure 18

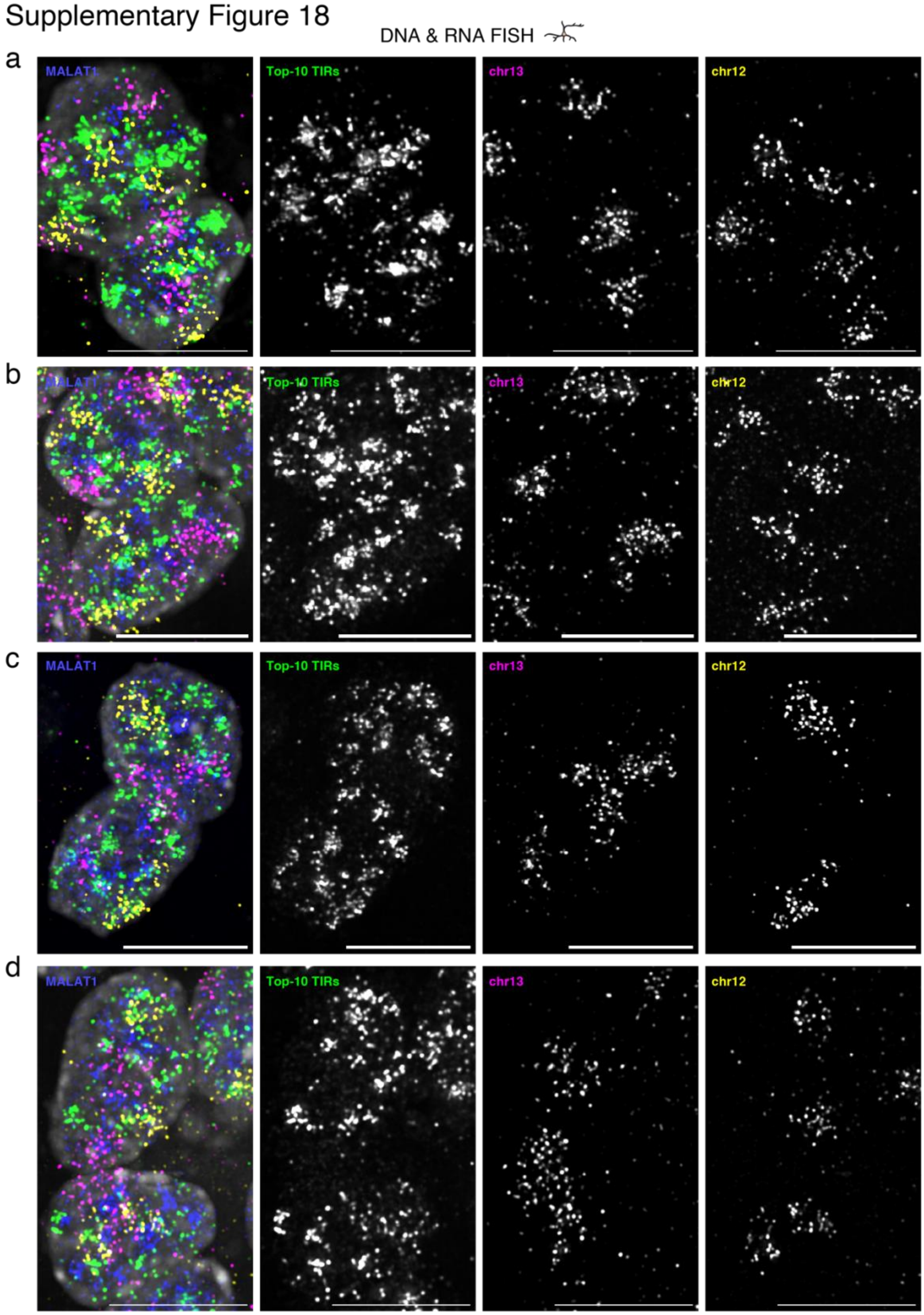


**Supplementary Figure 18. TIRs from the top-10 TIR source genes mingle more frequently with chr 12 than with chr 13. (a-d)** Maximum intensity projections of z-stack widefield microscopy images exemplifying the nuclear pattern of TIRs originating from the introns of the top-10 TIR source genes (green), together with chromosome (chr) 13 (magenta), which harbors the *NALF1* gene, and chr12 (yellow), which engages in extensive trans contacts with the top-55 TIR source genes but does not contain any of the top-10 TIR source genes, visualized by combined DNA and RNA FISH in NEU cells. Gray, DNA stained by Hoechst 33342. Scale bars: 10 µm. See **Supplementary Table 3** for the list of oligos composing the FISH probes shown in this figure. All the images in the figure were deconvolved using Deconwolf (https://deconwolf.fht.org/). A link to the Source Data and code to regenerate the plots displayed in this figure is provided in the Data Availability and Code Availability statements.

Supplementary Figure 19

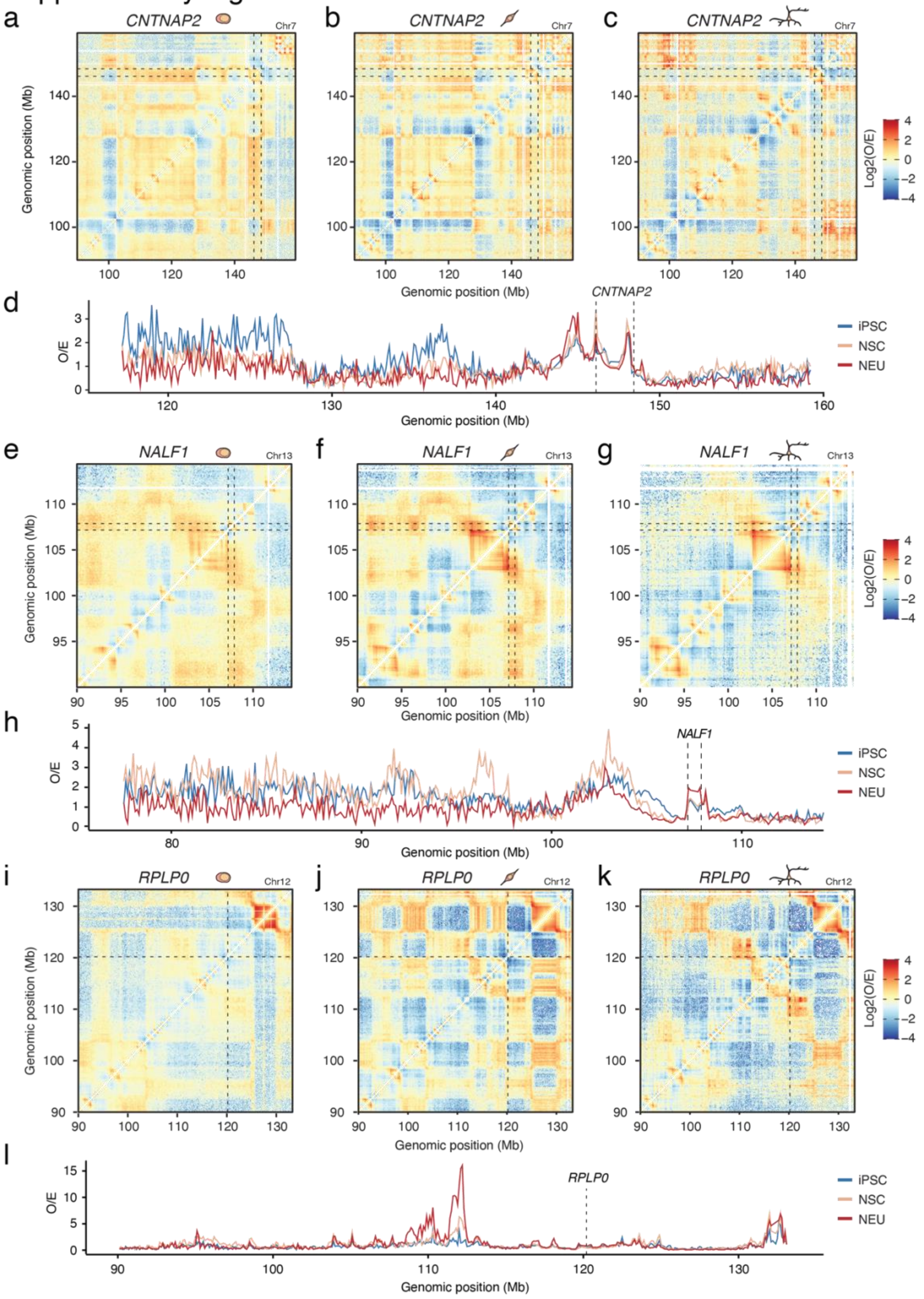


**Supplementary Figure 19. Long-range connectivity of TIR source loci weakens during neuronal differentiation.** (**a-c**) Observed-over-expected (O/E) ratio of DNA-DNA contacts detected by Hi-C (100 kb resolution) for a portion of chromosome (Chr) 7 encompassing the *CNTNAP2* TIR source gene, in iPSC, NSC, and NEU cells. The dashed black lines mark the genomic location of the *CNTNAP2* gene and its pair-wise contacts. (**d**) Hi-C O/E values calculated along the dashed black lines in (a-c), using the *CNTNAP2* gene as viewpoint. (**e-g**) As in (a-c) but for the *NALF1* TIR source gene located on Chr13. (**h**) As in (d) but referring to (e-g). (**i-k**) As in (a-c) but for the *RPLP0* TIR control gene located on Chr12. (**l**) As in (d) but referring to (i-k).

Supplementary Figure 20

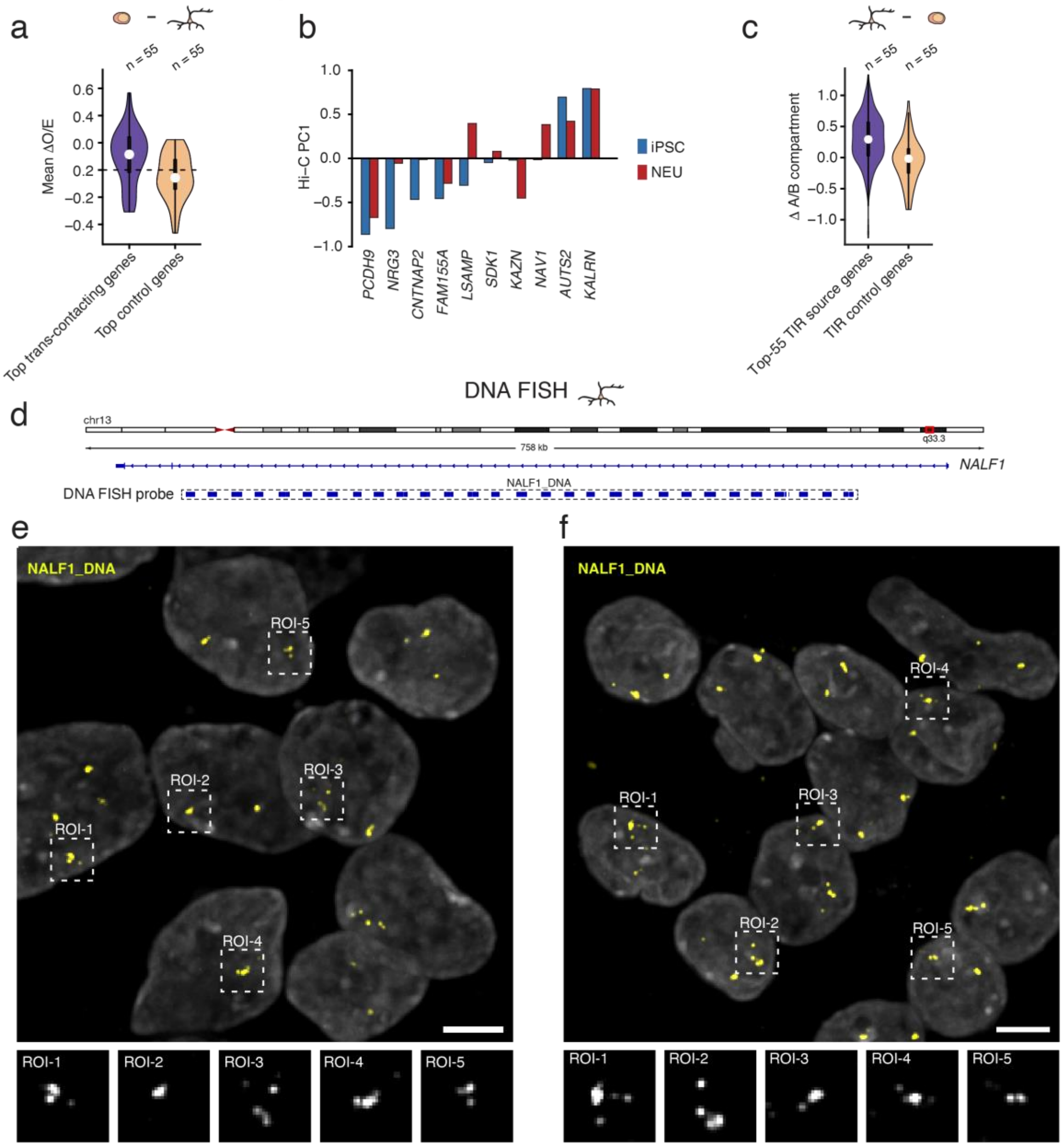


**Supplementary Figure 20. The *NALF1* TIR source locus shows partial unwinding in the nucleus of NEU cells.** (**a**) Distributions of the mean difference in the observed-over-expected (O/E) ratio of DNA-DNA contacts detected by Hi-C (100 kb resolution) within a ±30 Mb genomic window around the top-55 TIR source genes identified in NEU and matched TIR control genes (i.e., genes comparably expressed as TIR source genes based on RNA-seq and having their RNAs interacting in cis), calculated as iPSC minus NEU. Positive values indicate higher connectivity in iPSC compared to NEU. *n*, number of genes. (**b**) Mean Hi-C first eigenvector value (PC1) of 100 kb genomic bins overlapping with the gene bodies of the indicated top-10 TIR source genes, in iPSC and NEU cells. Positive values indicate A compartment (active chromatin), while negative values indicate B compartment (inactive chromatin). (**c**) Distributions of the difference in the Hi-C first eigenvector value for the top-55 TIR source genes identified in NEU and for matched TIR control genes, calculated as NEU minus iPSC. Positive values indicate a shift towards compartment A or away from compartment B in NEU compared to iPSC. *n*, number of genes. (**d**) Scheme of the genomic location and span of the DNA FISH probe used to visualize the TIR source gene *NALF1*. The vertical blue bars in the dashed box indicate individual oligos in the probe. The nomenclature of the probe is the same as in **Supplementary Table 3**. (**e, f**) Maximum intensity projections of z-stack widefield microscopy images exemplifying the staining pattern of the *NALF1* source locus (yellow) in NEU cells, visualized by DNA FISH using the probe described in (a). Gray, DNA stained by Hoechst 33342. Note that, before probe hybridization, the sample was treated with a combination of RNases to exclude binding of the DNA FISH probe to *NALF1* RNA or other RNA species. Scale bar: 5 μm. In (a) and (c), violins extend from minimum to maximum, boxplots extend from the 25$^{th}$ to the 75$^{th}$ percentile, white dots represent the median, whiskers extend from –1.5×IQR to +1.5×IQR from the closest quartile. IQR, inter-quartile range. All the images in the figure were deconvolved using Deconwolf (https://deconwolf.fht.org/). A link to the Source Data and code to regenerate the plots displayed in this figure is provided in the Data Availability and Code Availability statements.

Supplementary Figure 21

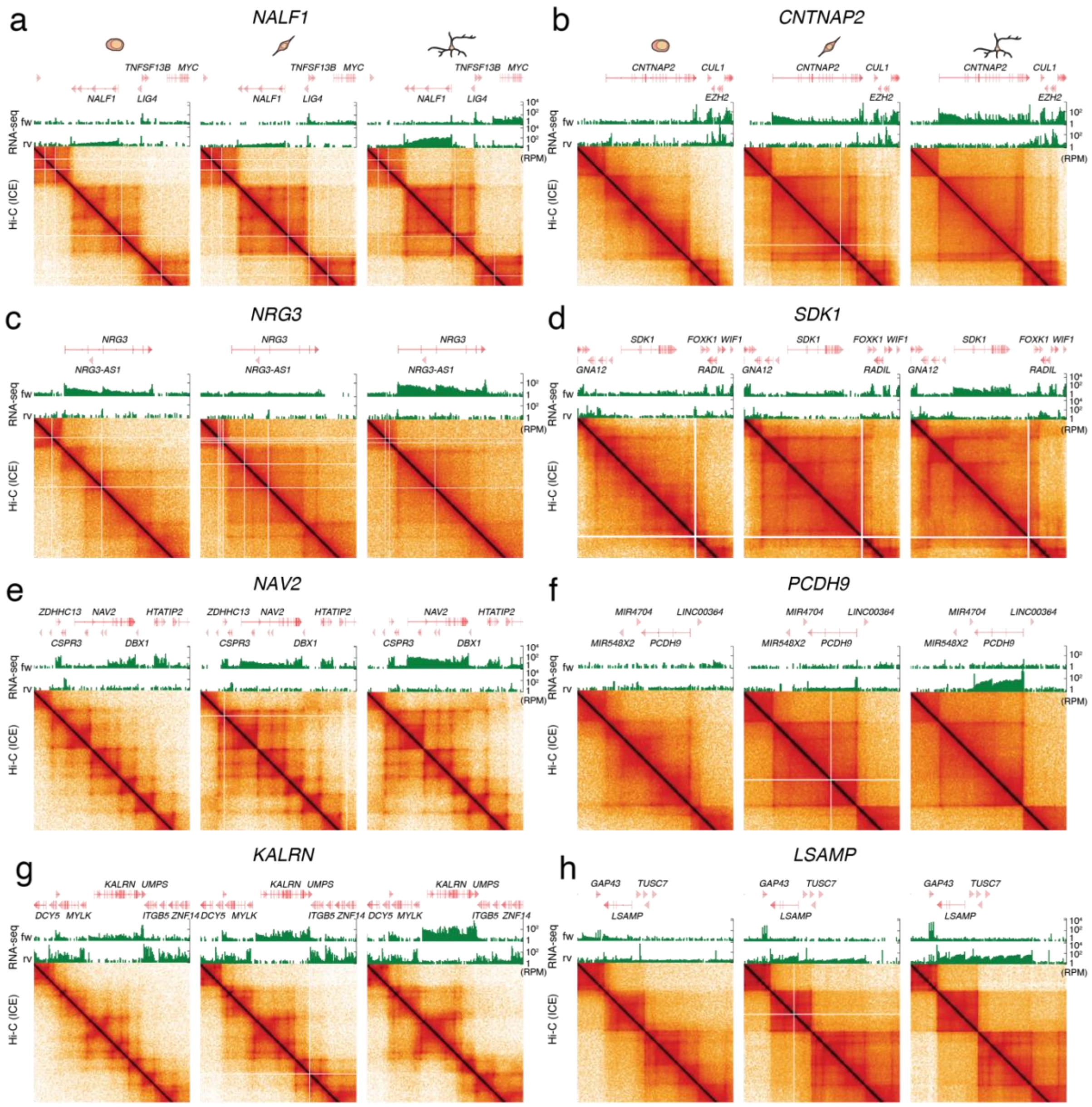


**Supplementary Figure 21. Well-defined TAD structures form around TIR source loci during neuronal differentiation.** (**a-h**) Hi-C contact matrix views in the neighborhood of eight of the top TIR source genes, in iPSC, NSC, and NEU. Top, gene tracks. Middle, RNA-seq coverage (green tracks) on the forward (fw) and reverse (rv) strand. Bottom, balanced Hi-C contact matrices with log-transformed scale. ICE, iterative correction and eigenvector decomposition. The Hi-C matrices were visualized in HiGlass using the following resolutions: *NALF1*, 8 kb; *NRG3*, 8 kb; *SDK1*, 16 kb; *NAV2*, 8 kb; *PCDH9*, 16 kb; *KALRN*, 8kb; *LSAMP*, 16 kb. A link to the Source Data and code to regenerate the plots displayed in this figure is provided in the Data Availability and Code Availability statements.

#### Supplementary Figure 22

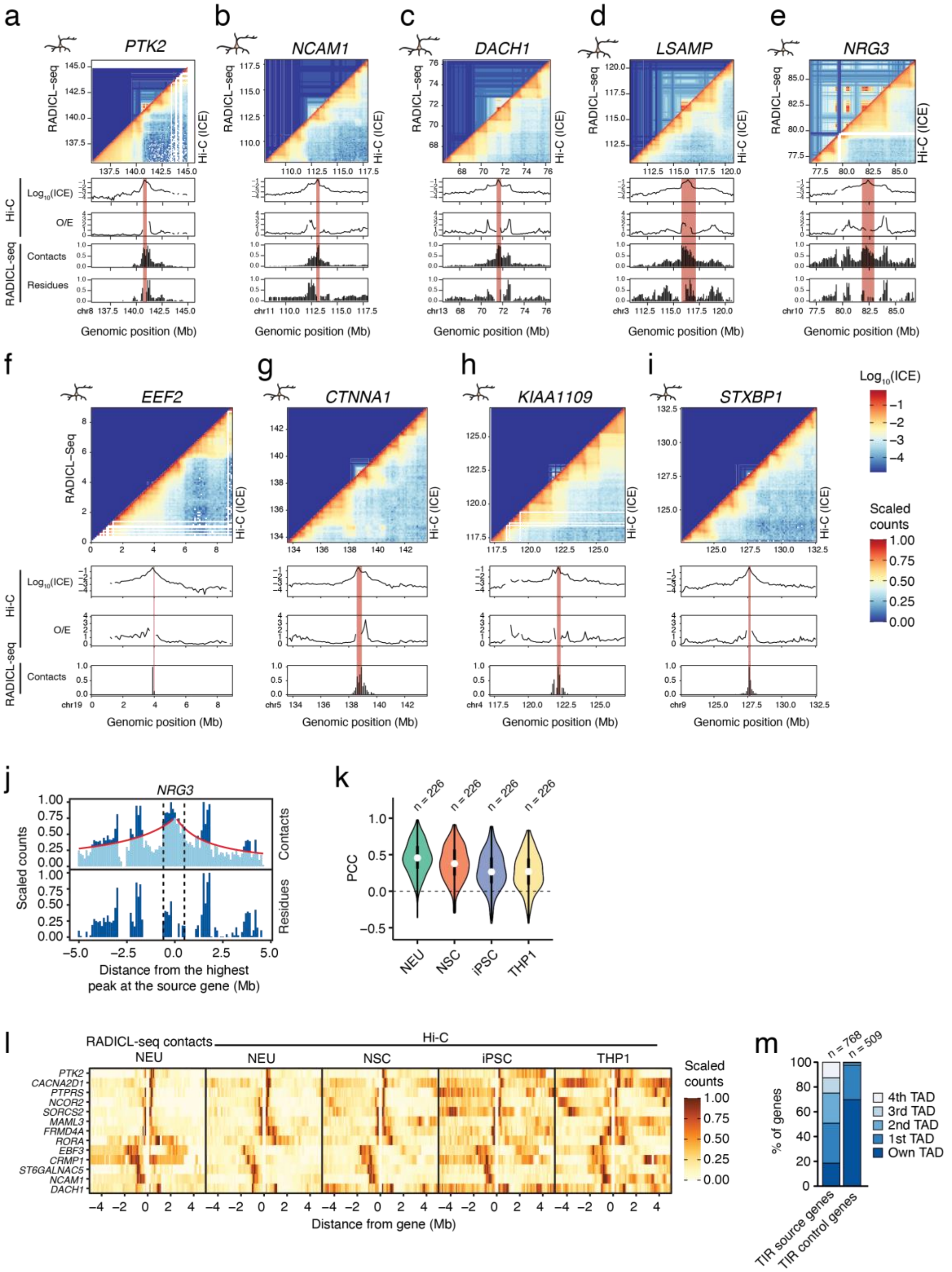


**Supplementary Figure 22. RADICL-seq RNA-DNA contact maps reflect the local chromatin structure.** (**a-e**) Side-by-side comparison between RNA-DNA contacts detected by RADICL-seq and DNA-DNA contacts measured by Hi-C, for three genes (*PTK2*, *NCAM1*,

*DACH1*) with a high correlation (PCC > 0.8) between RADICL-seq residues and Hi-C observed-over-expected (O/E) values in NEU, and for two of the top-10 TIR source genes identified by RADICL-seq in NEU (*LSAMP*, *NRG3*). Top heatmaps (100 kb resolution): putative DNA-DNA contact map (upper-left triangle) inferred by the percentiles of the shared RADICL-seq RNA contacts between the indicated DNA regions, and DNA-DNA Hi-C balanced contact map (bottom-right triangle). Bottom tracks: Hi-C and RADICL-seq tracks around the indicated genes used as viewpoint (red vertical bars). ICE, iterative correction and eigenvector decomposition. (**f-i**) As in (a-e) but for four control genes with expression levels comparable to TIR source genes, but whose RNA only engages in cis contacts. In the bottom tracks, RADICL-seq residues are not shown since TIR control genes, by definition, do not have trans contacts. (**j**) RADICL-seq contacts (top) and residues (bottom) (100 kb resolution) around the *NRG3* TIR source gene (vertical dashed lines). Residue calculation normalizes the total number of contacts identified by the number of contacts expected according to a background-decay-over-distance model (red line). The RADICL-seq residue signal is any signal above the model. (**k**) Distributions of the Pearson's correlation coefficient (PCC) calculated between the RADICL-seq residues within a genomic region ±5 Mb around the source loci of the TIRs identified by RADICL-seq in NEU and the O/E ratio of DNA-DNA contacts detected by Hi-C (100 kb resolution) in the same region, in each of the indicated cell types. Only TIR genes contacting in trans more than 80 distinct 100 kb genomic bins were included in the analysis to ensure sufficient data points for model fitting. Violins extend from minimum to maximum, boxplots extend from the 25$^{th}$ to the 75$^{th}$ percentile, white dots represent the median, whiskers extend from –1.5×IQR to +1.5×IQR from the closest quartile. IQR, inter-quartile range. (**l**) RADICL-seq (scaled counts, leftmost panel) and Hi-C O/E values (100 kb resolution) calculated in a 5 Mb window centered on 13 TIR source genes displaying a high correlation (PCC > 0.8) between RADICL-seq residues and Hi-C O/E values in NEU. (**m**) Percentage of genes that produce RNA forming DNA contacts within the same topologically associating domain (TAD) of the TIR source gene or extending to neighboring TADs, for the top-55 TIRs identified in NEU or for TIR control genes. *n*, number of genes. A link to the Source Data and code to regenerate the plots displayed in this figure is provided in the Data Availability and Code Availability statements.

#### Supplementary Figure 23

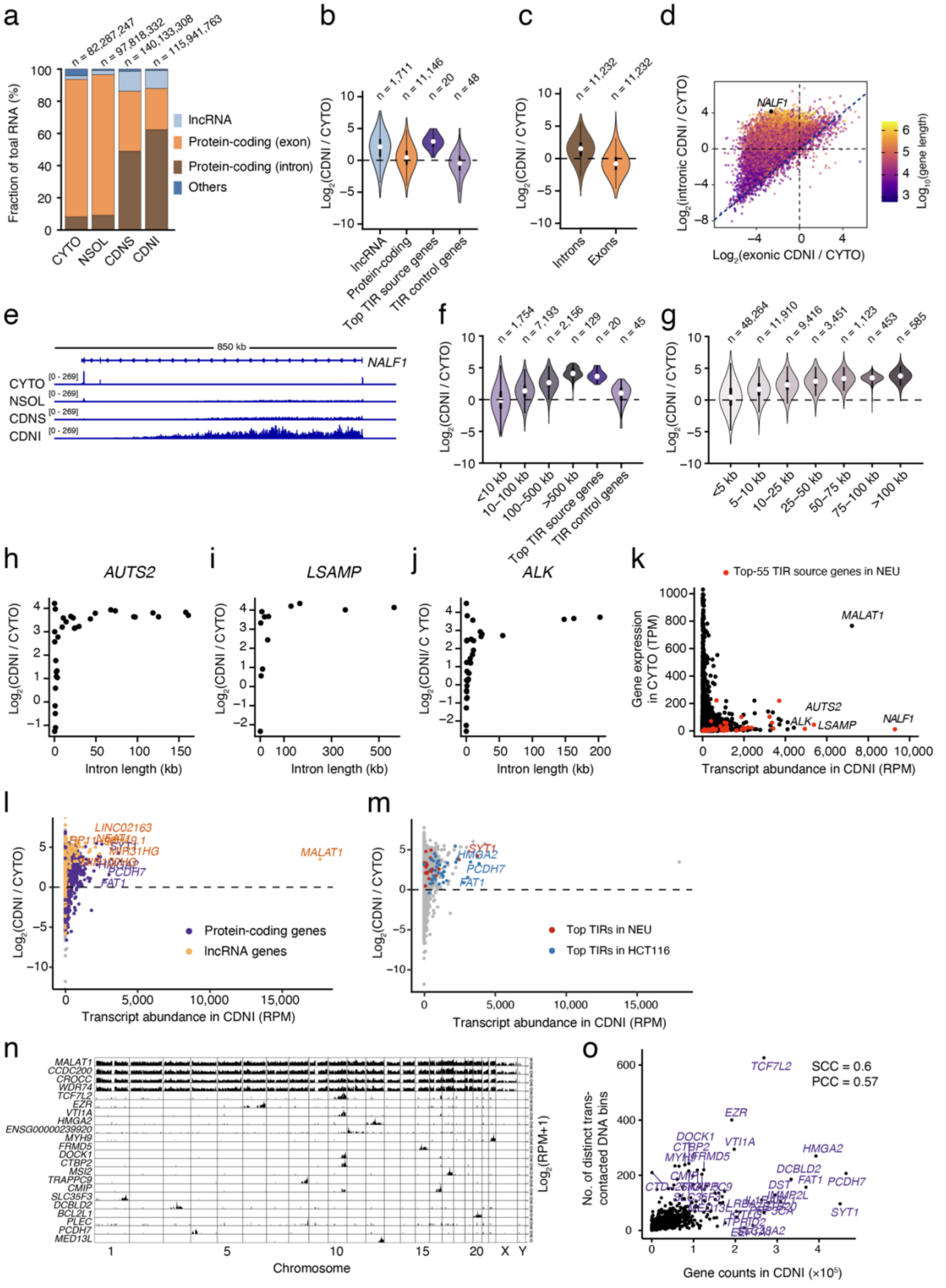


**Supplementary Figure 23. TIRs are enriched in the chromatin-depleted nuclear insoluble RNA fraction.** (**a**) Percentages of Total RNA-seq reads mapping to exonic or intronic regions of protein-coding genes, lncRNA genes, or other annotated genes, in four subcellular RNA fractions obtained from HCT116 cells. CYTO, cytosolic fraction. NSOL, nuclear soluble fraction. CDNS, chromatin-depleted nuclear soluble fraction. CDNI, chromatin-depleted nuclear insoluble fraction. *n*, number of reads. (**b**) Distributions of the gene enrichment in the CDNI fraction relative to the CYTO fraction (quantified as the fold change in RNA counts between the CDNI and the CYTO fraction) for the indicated gene sets, in HCT116 cells. *n*, number of genes in each set. RNA reads were quantified at the gene level. (**c**) As in (b) summing all the RNA reads mapping to intronic or exonic regions of each protein-coding gene, in HCT116 cells. (**d**) Relationship between the intronic and exonic RNA enrichment in the CDNI fraction relative to the CYTO fraction, in SH-SY5Y cells. For each gene, the total number of reads aligned to intronic vs. exonic regions was calculated. Each dot represents a gene, and the dots are colored based on the corresponding gene length. The *NALF1* TIR source gene is highlighted. (**e**) Integrative Genomics Viewer tracks displaying the RNA-seq coverage across the *NALF1* TIR source gene body (located on the reverse strand), for different cellular fractions of SH-SY5Y cells. (**f**) As in (b) with RNA reads quantified as summed intronic counts per protein-coding gene, in HCT116 cells. Gene sets are grouped by gene length, in addition to the top TIR source genes and matched control genes identified in NEU. *n*, number of genes. (**g**) As in (b) with RNA reads quantified at the level of individual introns of protein-coding genes, in SH-SY5Y cells. Introns are grouped by intron length, in addition to the introns of the top TIR source genes and the introns of matched control genes identified in NEU. *n*, number of introns. (**h-j**) Relationship between intron length and RNA enrichment in the CDNI fraction relative to the CYTO fraction, for the three genes with the highest RNA abundance in the CDNI fraction in SH-SY5Y cells. Note that these genes are part of the top-55 TIR source genes identified in NEU. *NALF1* is not shown because it contains only two introns. Each dot represents one intron. (**k**) Relationship between gene expression in the CYTO fraction (quantified as transcripts per million, TPM) and RNA abundance in the CDNI fraction (quantified at the gene level as reads per million, RPM), in SH-SY5Y cells. (**l**) Relationship between RNA abundance and gene enrichment in the CDNI fraction relative to the CYTO fraction, in HCT116 cells. Each dot represents one gene. (**m**) As in (l) with the top TIR source genes identified in NEU and HTC116 highlighted. (**n**) Genome-wide distributions of RADICL-seq contacts formed by RNAs derived from *MALAT1* and the top-21 TIR source genes identified in the HCT116 Replicate (Rep) 1 sample (see **Supplementary Table 1**). Note that the widespread contacts visible in the *CCDC200*, *CROCC*, and *WDR74* tracks are attributable to snRNA loci overlapping with those protein-coding genes. (**o**) Correlation between the RNA abundance in the CDNI fraction and the number of distinct 100 kb genomic bins contacted in trans by RNAs from the same genes (based on RADICL-seq), in HCT116 cells. SCC, Spearman's correlation coefficient. PCC, Pearson's correlation coefficient.

#### Supplementary Figure 24

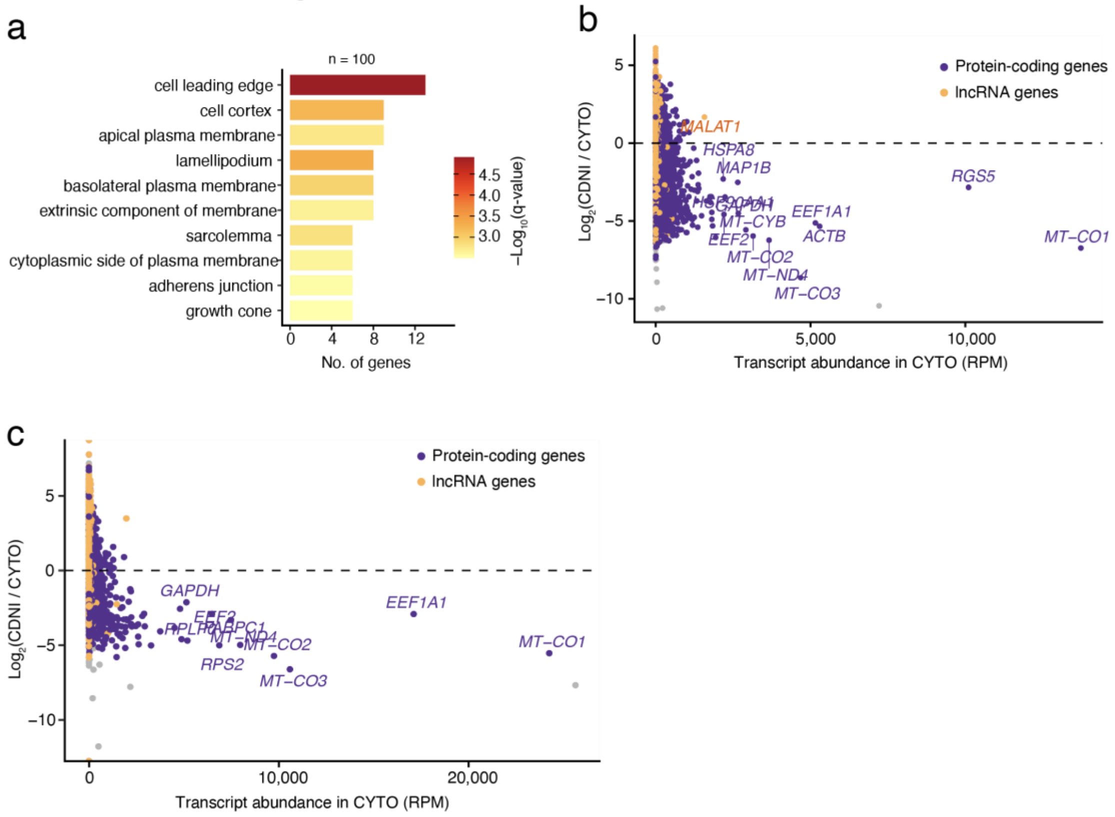


**Supplementary Figure 24. TIRs are enriched in the chromatin-depleted nuclear insoluble RNA fraction.** (**a**) Enriched gene ontology biological process terms based on the 100 most abundant genes (*n*) in the chromatin-depleted nuclear insoluble fraction (CDNI) relative to the cytosolic fraction (CYTO), in HTC116 cells. (**b**) Relationship between the RNA abundance (quantified at the gene level as reads per million, RPM) in the CYTO fraction and the gene enrichment in the CDNI fraction relative to the CYTO fraction (quantified as the fold change in RNA counts between the CDNI and the CYTO fraction, at gene level), in SH-SY5Y cells. Each dot represents one gene. (**c**) As in (b) but for HCT116 cells. A link to the Source Data and code to regenerate the plots displayed in this figure is provided in the Data Availability and Code Availability statements.

#### Supplementary Figure 25

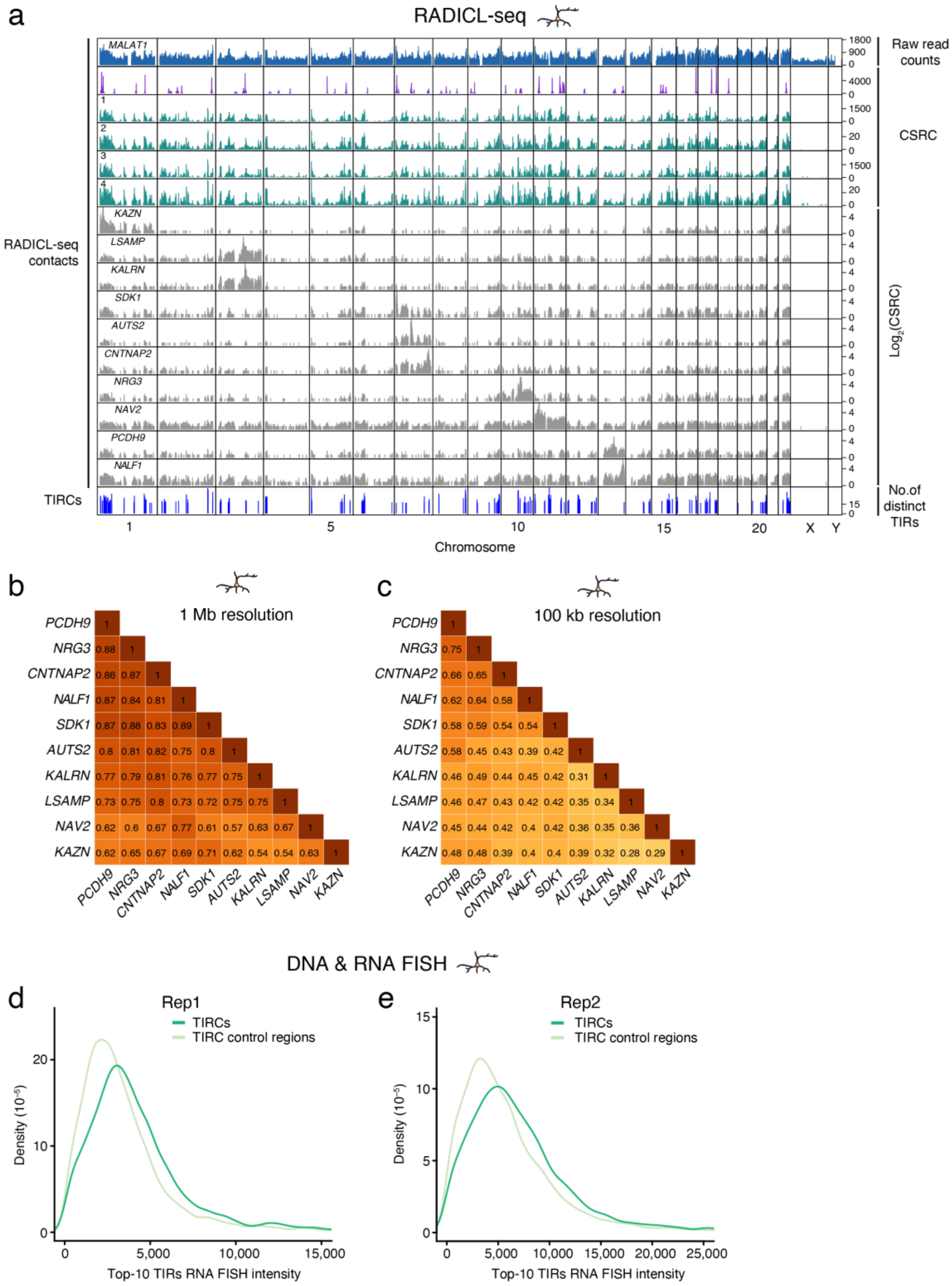


**Supplementary Figure 25. TIRs originating from different source genes contact a common set of TIR-contacted regions (TIRCs).** (**a**) Genome-wide distribution (100 kb bins) of: number of RADICL-seq contacts per bin, for *MALAT1* RNA (blue track); number of cis contacts per bin, for the top-55 TIR genes identified in NEU (purple track); number of trans contacts per bin, for the top-55 (green track #1) and 790 (green track #3) TIR genes; number of TIR source genes engaging in trans contacts per bin, for the top-55 (green track #2) and 790 (green track #4) TIR genes; number of trans contacts per bin, for the top-10 TIR genes in NEU (grey tracks); and TIRC regions defined as 100 kb genomic bins contacted by 16 or more distinct TIR species (bottom blue track, see **Computational Methods**, section 3). CSRC, CHiCANE significant read counts. (**b, c**) Pearson´s correlation coefficient matrix of genome-wide RADICL-seq trans contacts of the top-10 TIRs identified in NEU, at 1 Mb (b) and at 100 kb (c) resolution. (**d, e**) Probability density distributions of the RNA FISH signal intensity for TIRs originating from the top 10-TIR source genes overlapping with 10 TIRCs or 10 TIRC control loci detected by DNA FISH in NEU cells, in two biological replicates (Rep). All the intronic (IN) RNA FISH probes depicted in **Supplementary Fig. 6c** (except probes targeting *CAMTA1*) were pooled together and labeled with the same color. A link to the Source Data and code to regenerate the plots displayed in this figure is provided in the Data Availability and Code Availability statements.

#### Supplementary Figure 26

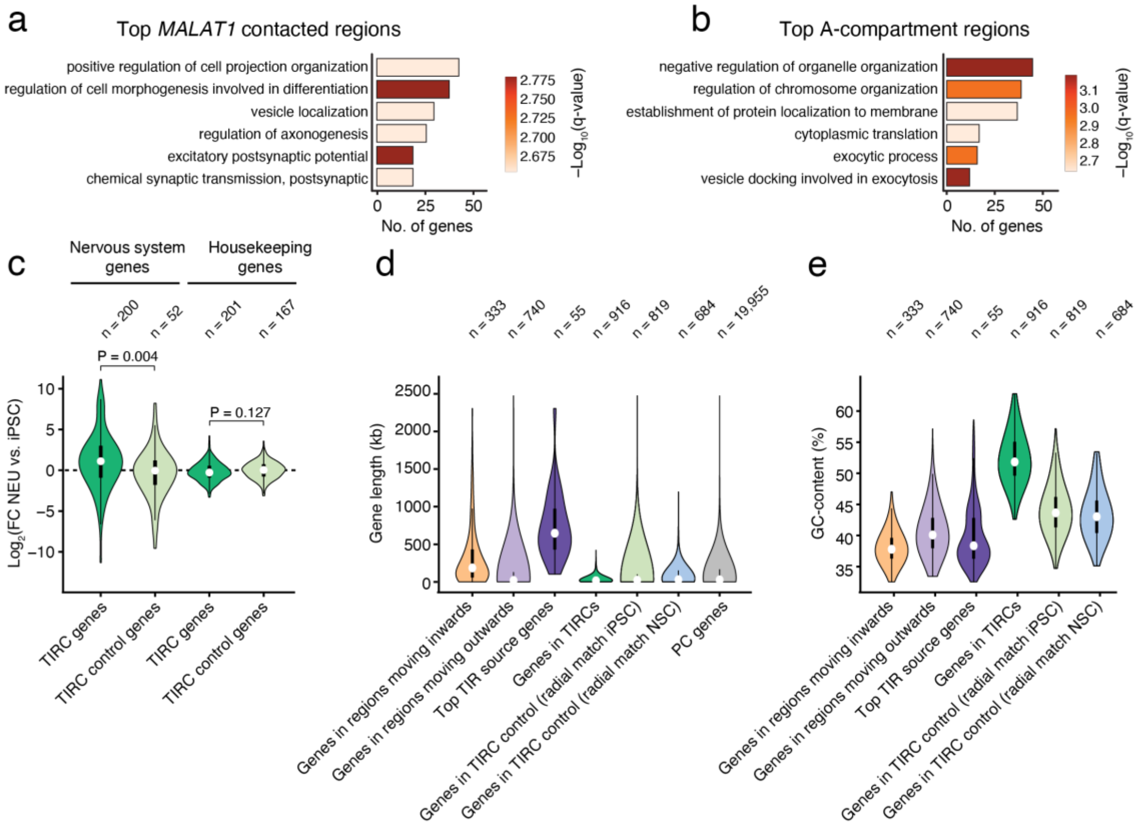


**Supplementary Figure 26. TIRCs are enriched in short active genes associated with neuronal functions.** (**a-b**) Enriched gene ontology biological process terms for expressed genes included in the top *MALAT1* contacted regions (a) and in A compartment regions displaying the highest Hi-C eigenvector values (b), in NEU. (**c**) Distributions of the fold change in gene expression measured by RNA-seq between NEU and iPSC, for nervous system genes and housekeeping genes located within TIRCs and TIRC control regions, respectively. Nervous system genes were defined based on Gene Ontology terms GO:0050877 and GO:0007399 from Gene Ontology (https://www.informatics.jax.org/vocab/gene_ontology/). The list of housekeeping genes was obtained from DOI: 10.1016/j.tig.2013.05.010. (**d**) Distributions of the gene length in the indicated gene groups. Outward- and inward-moving bins represent the top-1000 100 kb genomic bins ranked by the change in the ranked GPSeq score (rGS) from iPSC to NEU. Outward-moving bins show the largest decrease while inward-moving bins show the largest increase in the rGS (see **Supplementary Fig. 28a**). PC genes, protein-coding genes. (**e**) Distributions of the GC-content of 100 kb genomic bins overlapping the indicated gene groups or genomic regions. In (c-e), violins extend from minimum to maximum, boxplots extend from the 25th to the 75th percentile, white dots represent the median, whiskers extend from –1.5×IQR to +1.5×IQR from the closest quartile. IQR, inter-quartile range. A link to the Source Data and code to regenerate the plots displayed in this figure is provided in the Data Availability and Code Availability statements.

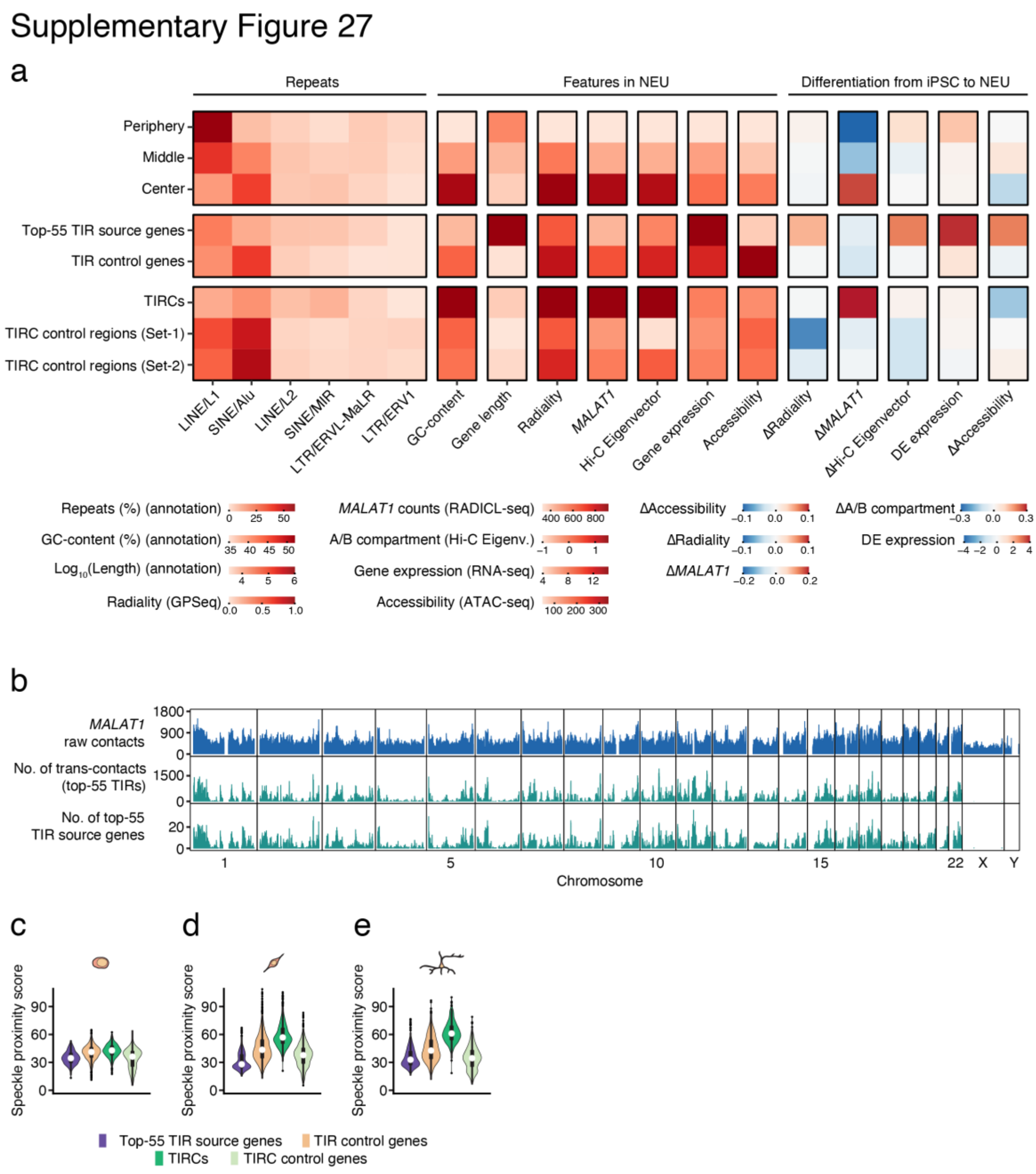


**Supplementary Figure 27. Genomic and epigenomic characterization of TIR source genes and TIRCs.** (**a**) Summary of the relative intensity of genomic features across the indicated region groups. Delta (Δ) indicates the difference in feature intensity between NEU and iPSC. Radial nuclear zones were defined based on the ranked GPSeq score (rGS) in NEU: periphery (rGS < 0.2), middle (rGS: 0.55–0.65), and center (rGS > 0.9). TIRC control regions (set 1 and 2): 100 kb genomic bins selected based on the absence of trans RNA contacts and on radial positioning similar to that of TIRCs in iPSC (Set-1) or NSC (Set-2). (**b**) Genome-wide profiles of *MALAT1* RNA trans contacts detected by RADICL-seq (top track), cumulative trans contacts made by any of the top-55 TIRs identified in NEU with each 100 kb genomic bin (middle track), and number of TIR source genes producing RNAs contacting each 100 kb bin (bottom track). (**c-e**) Distributions of the nuclear speckle proximity score inferred from the normalized (based on reads per million) *MALAT1* trans contacts for the indicated gene groups and genomic regions, in iPSC (c), NSC (d), and NEU (e). Violins extend from minimum to maximum, boxplots extend from the $25^{th}$ to the $75^{th}$ percentile, white dots represent the median, whiskers extend from $-1.5\times$IQR to $+1.5\times$IQR from the closest quartile. IQR, inter-quartile range. A link to the Source Data and code to regenerate the plots displayed in this figure is provided in the Data Availability and Code Availability statements.

#### Supplementary Figure 28

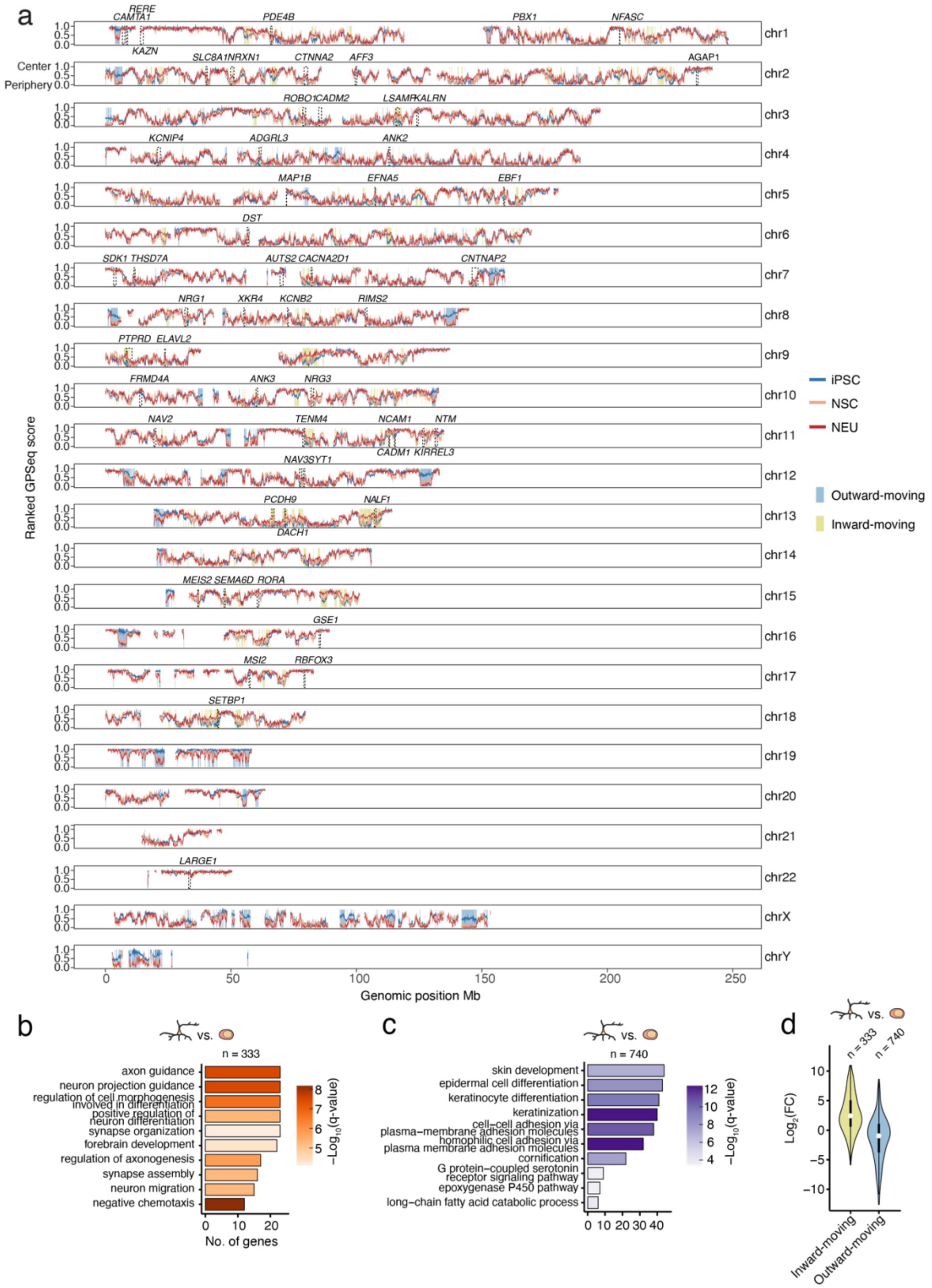


**Supplementary Figure 28. Genome-wide radial re-organization during neurodifferentiation**. (**a**) Genome-wide distribution of the ranked GPSeq score (rGS, 100 kb resolution) calculated from GPSeq, in iPSC, NSC, and NEU. The blue and yellow rectangles indicate the top-1000 outward- and inward-moving genomic bins, respectively, between iPSC and NEU. These bins were defined by ranking 100 kb genomic bins according to the change in their ranked GPSeq score (rGS) between iPSC and NEU. Outward-moving bins show the largest decrease while inward-moving bins show the largest increase in rGS. The dashed boxes mark the gene bodies of the top 55 TIR source genes identified in NEU by RADICL-seq. (**b**) Enriched gene ontology biological process terms based on the expressed genes (*n*) within the genomic bins moving inwards between iPSC and NEU shown in (a). (**c**) As in (b) for the outward-moving genomic bins shown in (a). (**d**) Distributions of the $Log_2$ fold change ($Log_2FC$) in gene expression within inward-moving and outward-moving genomic regions defined in (a), calculated from RNA-seq data using DESeq2 (NEU versus iPSC; positive $Log_2FC$ indicates upregulation in NEU). *n*, number of genes. Violins extend from minimum to maximum, boxplots extend from the $25^{th}$ to the $75^{th}$ percentile, white dots represent the median, whiskers extend from –1.5×IQR to +1.5×IQR from the closest quartile. IQR, inter-quartile range. A link to the Source Data and code to regenerate the plots displayed in this figure is provided in the Data Availability and Code Availability statements.

#### Supplementary Figure 29

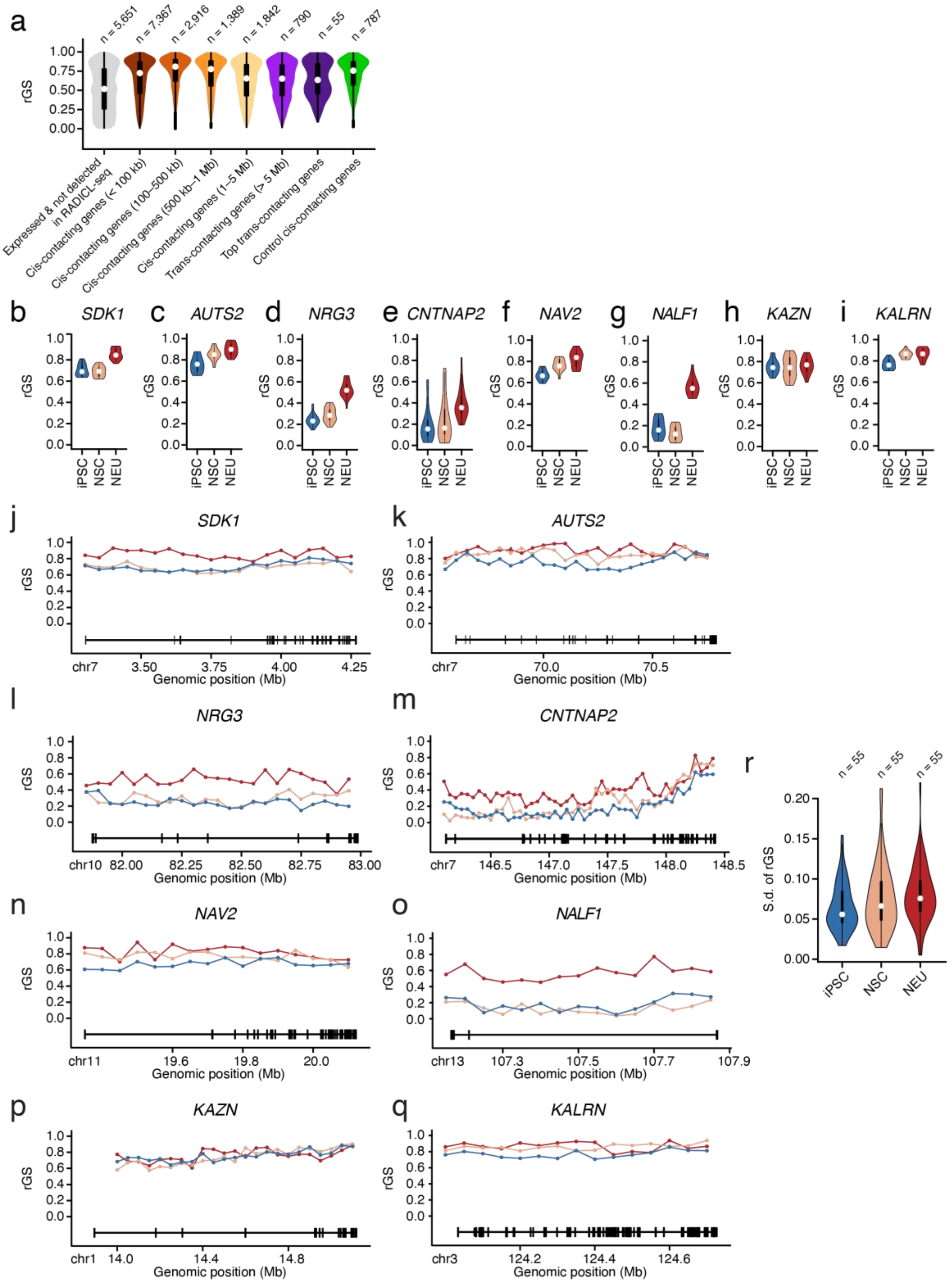


**Supplementary Figure 29. Radial arrangement of TIR source genes.** (**a**) Distribution of the ranked GPSeq score (rGS, 100 kb resolution) for different gene groups classified based on the expression levels, as detected by RNA-seq, and on the type of RNA-DNA contacts, as detected by RADICL-seq. *n*, number of genes in each group. (**b-q**) rGS shown as aggregated distributions (violin plots) or along the gene body (dotted line plots, 50 kb resolution) for 8 out of the top-10 TIRs source genes, in iPSC, NSC and NEU. The black tracks under the dotted line plots schematically represent the gene body (horizontal lines) and exons (vertical lines). *SDK1*, *AUTS2*, *NRG3*, *CNTNAP2*, *NAV2*, *KAZN*, and *KALRN* are on the positive strand. *NALF1* is on the negative strand. (**r**) Distribution of the standard deviation (S.d.) of the rGS per gene, in iPSC, NSC and NEU, for the top-55 TIR source genes identified by RADICL-seq in NEU. In all the figure, violins extend from minimum to maximum, boxplots extend from the 25$^{th}$ to the 75$^{th}$ percentile, white dots represent the median, whiskers extend from $-1.5\times IQR$ to $+1.5\times IQR$ from the closest quartile. A link to the Source Data and code to regenerate the plots displayed in this figure is provided in the Data Availability and Code Availability statements.

Supplementary Figure 30

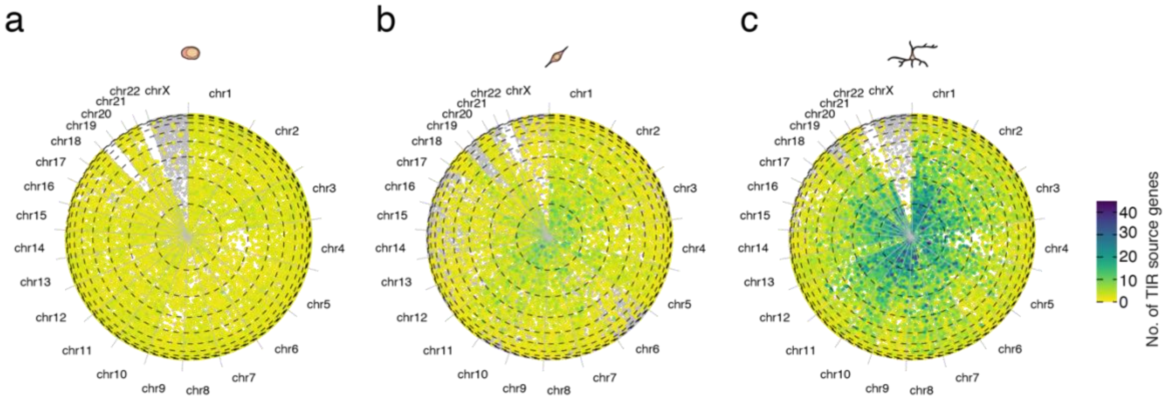


**Supplementary Figure 30. Genomic regions engaging with multiple distinct TIRs preferentially localize to central nuclear regions.** (**a**) 'Pizza plots' showing the number of TIRs (expressed as number of distinct source genes) contacting 100 kb genomic bins radially placed based on GPSeq data from iPSC (a), NSC (b), and NEU (c). TIRs were identified independently in each cell type. A link to the Source Data and code to regenerate the plots displayed in this figure is provided in the Data Availability and Code Availability statements.

Supplementary Figure 31

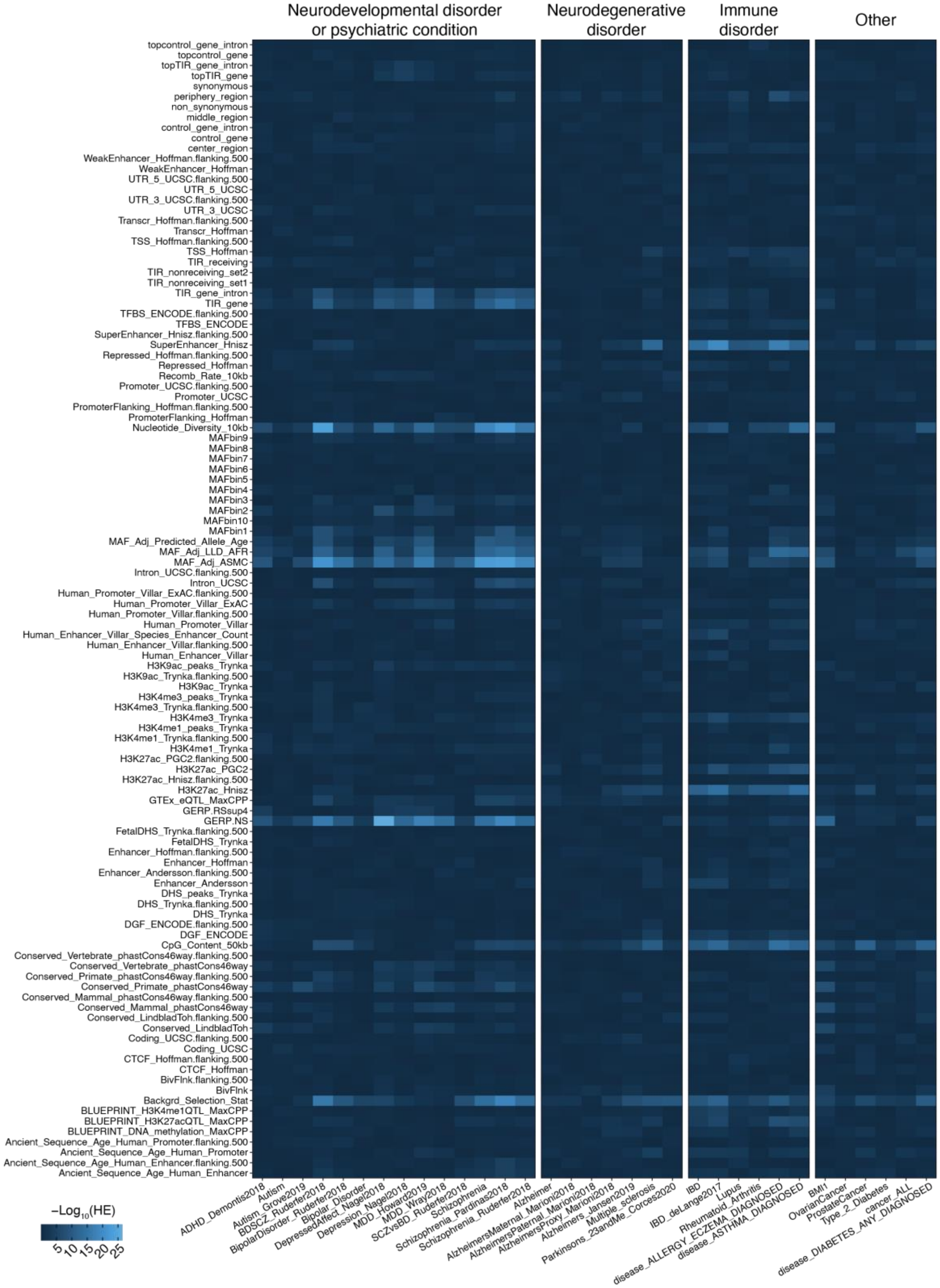


**Supplementary Figure 31. TIR source genes are significantly associated with risk loci for mental, behavioral, and neurodevelopmental disorders.** Heatmap showing the P value of the heritability enrichment (HE) of SNPs identified across multiple GWAS datasets grouped by trait analyzed (x-axis). The annotation groups are listed on the y-axis. Traits and annotations are based on the S-LDSC results from 33 GWAS studies (see **Supplementary Table 4**). A link to the Source Data and code to regenerate the plots displayed in this figure is provided in the Data Availability and Code Availability statements.

Supplementary Figure 32

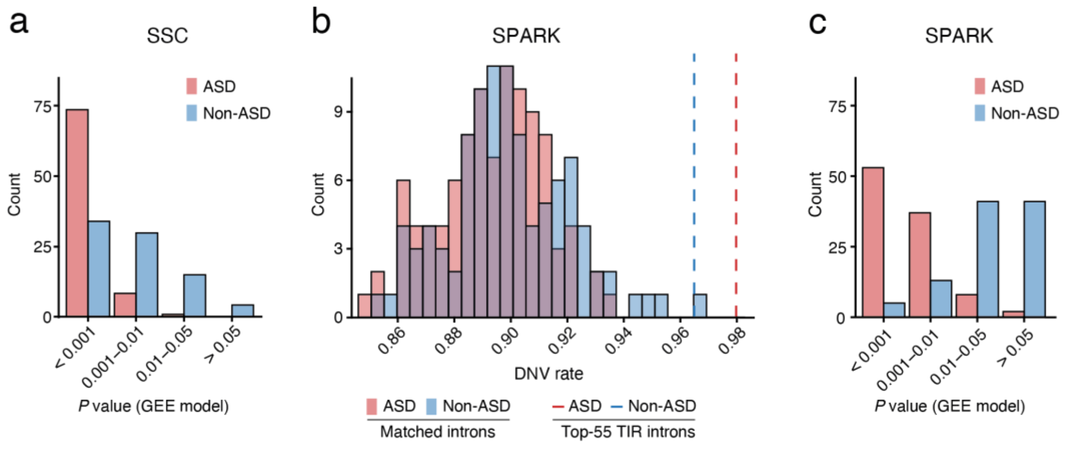


**Supplementary Figure 32. Enrichment of de novo variants (DNV) in two autism cohorts.** (**a**) Counts of significance level (*P* values) in 100 Generalized Estimating Equation (GEE) models assessing the correlation between DNV counts and intron set type (introns of the top-55 TIR source genes detected by RADICL-seq in NEU versus randomly matched introns from neuronally expressed genes), in the Simons Simplex Collection (SSC) cohort. (**b**) Distribution of the DNV event rate in 100 randomly matched intron sets (from genes expressed in NEU) in the Simons Foundation Powering Autism Research for Knowledge (SPARK) cohort, for the top-55 TIR introns. Vertical lines represent DNV event rates in the introns of the top-55 TIR source genes detected by RADICL-seq in NEU. (**c**) As in (a) for the SPARK cohort. A link to the Source Data and code to regenerate the plots displayed in this figure is provided in the Data Availability and Code Availability statements.

#### 2. Supplementary Tables

Because of their large size, alle tables are provided as separate Excel files.

**Supplementary Table 1.** List of sequencing datasets analyzed in this study.

**Supplementary Table 2.** List of the top-55 TIRs identified in this study.

**Supplementary Table 3.** List of DNA and RNA FISH probes used in this study.

**Supplementary Table 4.** List of GWAS datasets used for stratified linkage disequilibrium score regression (S-LDSC) analysis in this study.

**Supplementary Table 5.** Coefficients of the GEE models assessing the association between DNV burden and autism status, adjusting for sex and clustering by family ID.
